# Extracellular Delivery of Functional Mitochondria Rescues the Dysfunction of CD4^+^ T Cells in Aging

**DOI:** 10.1101/2021.02.21.432151

**Authors:** Colwyn A. Headley, Shalini Gautam, Angelica Olmo-Fontanez, Andreu Garcia-Vilanova, Varun Dwivedi, Anwari Akhter, Alyssa Schami, Kevin Chiem, Russell Ault, Hao Zhang, Hong Cai, Alison Whigham, Jennifer Delgado, Amberlee Hicks, Philip S. Tsao, Jonathan Gelfond, Luis Martinez-Sobrido, Yufeng Wang, Jordi B. Torrelles, Joanne Turner

## Abstract

Mitochondrial dysfunction alters cellular metabolism, increases tissue oxidative stress, and may be principal to the dysregulated signaling and function of CD4^+^ T lymphocytes in the elderly. In this proof of principle study, we investigated whether the transfer of functional mitochondria into CD4^+^ T cells that were isolated from old mice (aged CD4^+^ T cells), could abrogate aging-associated mitochondrial dysfunction, and improve the aged CD4^+^ T cell functionality. Our results show that the delivery of exogenous mitochondria to aged non-activated CD4^+^ T cells led to significant mitochondrial proteome alterations highlighted by improved aerobic metabolism and decreased cellular mitoROS. Additionally, mito-transferred aged CD4^+^ T cells showed improvements in activation-induced TCR-signaling kinetics displaying markers of activation (CD25), increased IL-2 production, enhanced proliferation *ex vivo*. Importantly, immune deficient mouse models (RAG-KO) showed that adoptive transfer of mito-transferred naive aged CD4^+^ T cells, protected recipient mice from influenza A and *Mycobacterium tuberculosis* infections. These findings support mitochondria as targets of therapeutic intervention in aging.

## 1. Introduction

Biological aging positively correlates with acute and chronic microbial infections and diseases that are of increased severity, poor response to vaccines, increased incidence of autoimmunity, and is accompanied by chronic low-grade inflammation [1-4]. CD4^+^ T cells bridge communication between innate and adaptive arms of the immune system, and their dysfunction contributes to the aggravation of these aging-associated changes. The collective pool of naïve, memory, and effector CD4^+^ T cells succumb to aging-associated perturbations. For example, thymic involution stunts the generation of properly educated naïve CD4^+^ T cells [5-8]. CD4^+^ memory T cells generated later in life function at subpar levels compared to those generated in younger years [5, 8-10]. Moreover, the kinetics of CD4^+^ effector T cells (*i.e.* activation and resolution) in aged mice are enervated [11, 12].

Mitochondrial activity orchestrates a significant number of CD4^+^ T cell signaling events, including shifts in metabolism and ATP production, intracellular Ca^2+^ mobilization, gene induction (NFAT, NF-κB, and AP-1), and production of reactive oxygen species (ROS) [13-23]. Perturbations in T cell intracellular signaling networks are likely central to the collective dysregulation and dysfunction observed in T cells of the elderly [24-27]. Indeed, redox dysregulation due to aging-associated mitochondrial dysfunction might manifest as disturbances in T cell function including homeostasis, activation, proliferation, and their ability to modulate of inflammation [28].

The optimum levels of ROS necessary for the induction of CD4^+^ T cell signaling cascades are mainly generated as byproducts of oxidative phosphorylation (Ox-Phos). Ox-Phos occurs on the inner membrane of mitochondria via the electron transport chain (ETC) [15, 29-32]. Superoxide anions (O_2_ ^•^-) and hydrogen peroxide (H O) are the principal mitochondrial ROS (mitoROS) essential for CD4^+^ T cell redox signaling [15, 23, 33]. Due to its permeability, H_2_O_2_ serves as a secondary messenger molecule [33-36]. For instance, exposed thiol moieties (-SH) on proteins can be oxidized by H_2_O_2_, altering protein conformation to favor either increases or decreases in enzymatic activities [37, 38]. Concurrently, mitoROS contribute to the formation of other increasingly destructive ROS. For example, O ^•^- can rapidly react with nitric oxide (NO) to form peroxynitrite (ONOO-), while H_2_O_2_ can react with O ^•^-, transition metals, lipids, and proteins found within mitochondria and cells, propagating the generation of hydroxyl (OH•), and peroxyl (ROO•) radicals [30-32, 39-42]. Therefore, shifts in the concerted generation and elimination of mitoROS, as seen in many aged cells, tissues, and organs, contribute to abnormal mitochondrial function which in turn perpetuates redox dysregulation and subsequent T cell dysfunction [43, 44].

Cellular energy demand is dynamic, and mitochondria must continually fuse and fragment in response to available nutrients and biological stimuli [45-47]. Increases in mitoROS impact mtDNA stability; continual fusion and fission events may also propagate the inter-mitochondrial dissemination of damaged mtDNA [45, 48-52]. Improper translation and assembly of mtDNA encoded proteins, in turn, hinders proper ETC assembly. Often this is associated with increased mitochondrial electron leaks and excess formation of mitoROS, favoring a pro-oxidant intracellular environment [53-55]. In short, mitochondrial dysfunction manifests as quantifiable abnormalities in the assembly of the ETC, mitochondrial membrane potential, excessive production of mitoROS, and abnormal production of ATP via Ox-Phos [24, 53, 56]. These perturbations impact the metabolic fitness and the redox-sensitive signaling pathways of CD4^+^ T cells [13, 25, 53, 57-60].

Therapeutic interventions that can simultaneously improve mitochondrial biogenesis and re-establish redox equilibrium inCD4^+^ T cells may abrogate aging-associated T cell dysfunction [22, 27, 28, 61]. This proof-of-concept study assessed whether the exogenous delivery of functional mitochondria to CD4^+^ T cells (mito-transfer) from old mice and elderly humans could re-establish the cellular redox balance and positively impact CD4^+^ T cell function. Compared to non-manipulated CD4^+^ T cells, mito-transfer remodeled the mitochondrial proteome of CD4^+^ T cells; improving their aerobic metabolism resulting in decreased mitoROS. Mito-transferred CD4^+^ T cells from old mice also displayed enhanced intracellular phosphorylation of key proteins essential to TCR signal transduction, increased cellular production of cytokines including IL-2, increased the number of cells expressing CD25, and improved cellular proliferation after *ex vivo* activation. Importantly, the adoptive transfer of mito-transferred naïve CD4^+^ T cells from old mice into Rag1-KO mice protected mice against influenza A and *Mycobacterium tuberculosis (M.tb)* infections. These results open the possibility for future translational studies into the immunological and therapeutic implications of directed mito-transfer in aged lymphocytes to re-establish their loss of function.

## 2. Methods

### Mice

Specific pathogen-free C57BL/6 young (2 to 4 months) and old (18 to 24 months) mice were obtained from the National Institute on Aging, Charles River Laboratories (Wilmington, MA, USA), and The Jackson Laboratories (JAX, Bar Harber, ME, USA). Rag1-KO (B6.129S7-Rag1^tm1Mom/^J, 10 weeks old) mice were obtained from JAX. Mice were housed in groups of 5 on individually ventilated cages and fed a standard chow diet, *ad libitum*, for the duration of the study. All procedures were approved by the Texas Biomedical Institutional Laboratory Animal Care and Use Committee (IACUC) under protocols 1608 MU and 1735 MU.

### Isolation of CD4^+^ T cells from mice

Young and old mice were euthanized by CO_2_ asphyxiation, and spleens surgically excised. Isolated spleens were individually placed in 6 well plates containing 4 ml of chilled (4°C) complete DMEM and gently hand homogenized using sterile syringe stoppers. Spleen homogenate was passed through a 70 µm microfilter to remove large debris. The resulting single-cell suspensions were centrifuged for 5 min at 300 x *g* at 4°C and suspended in Gey’s lysis buffer (8 mM NH_4_Cl, 5 mM KHCO_3_) for 1 min to lyse erythrocytes, and then neutralized with an equivalent volume of complete DMEM. Splenocytes were washed and suspended in separation buffer (10 mM D-Glucose, 0.5 mM MgCl_2_, 4.5 mM KCl, 0.7 mM Na_2_HPO_4_, 1.3 mM NaH_2_PO_4_, 25 mM NaHCO_3_, 20 mM HEPES, 5 µM DETC, 25 µM deferoxamine) containing appropriate antibody-magnetic particles for positive selection (α-CD4), or negative selection (α-CD8, α-CD11b, α-CD109, α-CD24, α-CD45, α-CD49b, α-Ly-6G, α-χδ TCR, α-TER-119). The resulting population of CD4^+^ T cells (85-95% purity) was used for downstream experiments.

### Quantitation of intracellular ROS

For flow cytometry-based ROS experiments, CD4^+^ T cells were incubated in media containing MitoSOX (5 µM) or CellROX (5 µM) for 30 min at 37°C, to measure mitochondrial and cytosolic ROS, respectively. Cells were washed twice with phosphate-buffered saline (PBS, pH 7.4) and stained with antibodies targeting surface proteins of interest (*i.e.* α-CD3, α-CD4, and α-CD69; 25 µg/ml) for 20 min before fixation in 2% paraformaldehyde. Flow cytometry-based ROS and other flow-based quantifications were performed on a BD-LSR II (BD Biosciences, NJ, USA), BD FACs Symphony, Cyan or CytoFLEX LX (Beckman Coulter, CA, USA) flow cytometers. Data were analyzed using FlowJo (FlowJo LLC-BD, NJ, USA) and GraaphPad Prism ver. 9 (Graphpad Software, CA, USA).

For electron paramagnetic resonance (EPR)-based ROS experiments, CD4^+^ T cells were probed for mitochondria-specific O_2_ ^•^- (*mt*O ^•^-) using Mito-Tempo-H. Briefly, CD4^+^ T cells were incubated in EPR buffer (10 mM D-Glucose, 0.5 mM MgCl, 4.5 mM KCl, 0.7 mM Na_2_HPO_4_, 1.3 mM NaH_2_PO_4_, 25 mM NaHCO_3_, 20 mM HEPES, 5 µM DETC, 25 µM deferoxamine), containing 100 µM Mito-Tempo-H, for ∼45 min at 37°C. EPR spectra were obtained on a Bruker EMXnano ESR system (Bruker Corporation, MA, USA) using the following parameters: microwave frequency, 9.6 GHz; center field, 3430.85 G; modulation amplitude, 6 G; microwave power at 25.12 mW; conversion time of 71.12 ms; time constant of 20 ms; sweep width of 100 G; receiver gain at 1.0 × 10^5^; and a total number 5 of scans. The Two-Dimensional (2-D) spectra were integrated, baseline corrected, and analyzed using spin-fit and GraphPad software.

### Isolation of mouse embryonic fibroblasts

Mouse embryonic fibroblasys (MEFs) were isolated from 12-14 days post-coitum C57BL/6 mice as described [62]. Briefly, mouse embryos were aseptically isolated, minced, and digested in 0.05% trypsin and crude DNase solution for 30 min at 37°C, 5% CO_2_, in a humidified incubator. Primary MEFs were cultured in sterile T-150 flasks (Corning, NY, USA), in 50:50 F12/DMEM (Corning) supplemented with 10% FBS, 1% HEPES, 2% NEAA, 1 mM sodium pyruvate, 2 mM L-glutamine, and 100 µg/ml primocin. MEFs were sub-passaged as necessary, with MEFs from passages 2-3 being used for all experiments.

### Mitochondrial isolation and transfer

Mitochondria were isolated from donor MEFs and centrifuged into CD4^+^ T cells using a published protocol [63]. Briefly, MEFs were homogenized in SHE buffer [250 mM sucrose, 20 mM HEPES, 2 mM EGTA, 10 mM KCl, 1.5 mM MgCl_2_, and 0.1% defatted bovine serum albumin (BSA)], containing complete Minitab protease inhibitor cocktail (Sigma Alrich, MO, US). The cell homogenates weree centrifuged at 800 x *g* for 3 min at 4°C to remove cellular debris. The resulting supernatant was recovered and centrifuged at 10,000 x *g* for 5 min to pellet isolated mitochondria. The CD4^+^ T cells and isolated mitochondria were suspended in 150 μL cold PBS in V-bottom 96 well plates, and plates were centrifuged at 1,400 x *g* for 5 min at 4°C. After centrifugation, cells were washed 2-3 times with cold PBS before further experimentation. Independent of mito-transfer, all experimental groups were centrifuged accordingly.

### Determination of cardiolipin content

Mitochondrial cardiolipin content was determined using a commercially available cardiolipin assay kit (BioVision Inc, CA, USA) per the manufacturer’s instructions. Briefly, isolated mitochondria were stained with a cardiolipin-specific fluorometric probe 1,1,2,2-tetrakis[4-(2-trimethylammonioethoxy)-phenyl]ethene tetrabromide, and quantified against a cardiolipin standard curve.

### Transmission electron microscopy (TEM)

CD4^+^ T cells were fixed in PBS containing 4% glutaraldehyde and 1% paraformaldehyde for 1 h, after which, cells were rinsed in PBS for 20 min. Samples were fixed in 1% Zetterqvist’s buffered osmium tetroxide for 30 min followed by an alcohol gradient (70%, 95%, 100%) and propylene oxide dehydration (10 min per solution). Samples were blocked in 1:1 propylene oxide/resin for 30 min, and 100% resin for 30 min (at 25 psi). Samples were then polymerized overnight at 85°C and stained with 1% uracil acetate and alkaline lead citrate. Images were captured using a JEM-1400 transmission electron microscope (JEOL, MA, USA) at 25,000X and 50,000X magnification, and analyzed using FIJI software (ImageJ ver. 1.52p, NIH, MD, USA) [64].

### Extracellular flux-based assays

Kinetic analysis of cellular glycolysis and oxygen consumption and the rate of ATP production was accomplished through the use of the Extracellular Flux Analyzer XF^e^96 (Agilent/Seahorse, CA, USA). Briefly, 2.5 × 10^5^ CD4^+^ T cells were plated onto poly-D-Lysine pre-coated (0.1 mg/ml) XF^e^96 microplates. Using real-time injections, the rates of mitochondrial respiration, glycolysis, and ATP production were measured via oxygen consumption rate (OCR, pmol/min) and extracellular acidification rate (ECAR, mPh/min).

For mitochondrial stress and ATP production assays, cells were placed in XF assay media supplemented with 5.5 mM D-glucose, 2 mM L-glutamine, and 1 mM sodium pyruvate. Sequential injections of oligomycin (1 µM; O), carbonyl cyanide-4-(trifluoromethoxy)phenylhydrazine (1.5 µM; FCCP), and rotenone: antimycin (1 µM, R/A) were used for the mitochondrial stress test, while injections of O and R/A alone were used to determine ATP production rate.

For the glycolysis stress test, cells were assayed in D-Glucose deficient XF media containing L-glutamine, and injections of glucose (10 mM), oligomycin (1 µM), and 2-deoxyglucose (50 mM; 2-DG) were used. For the glycolytic rate assay, cells were assayed in complete XF media and sequential injections of R/A (1 µM) and 2-DG were used.

For acute CD4^+^ T cell activation assays, cells were placed in XF assay media supplemented with 5.5 mM glucose, 2 mM L-glutamine, and 1 mM sodium pyruvate. Sequential injections of PMA/Ionomycin and 2-DG were used to determine the kinetics of oxygen consumption and extracellular acidification after CD4^+^ T cell activation.

### Fluorescent glucose uptake assay

CD4^+^ T cells were incubated in D-glucose-free DMEM for 20 min before the addition of complete DMEM supplemented with 50 µM 2-deoxy-2-[(7-nitro-2,1,3-benzoxadiazol-4-yl)amino]-D-glucose (2-NBDG), a fluorescent glucose analog, for 30 min. Cells were washed twice in cold PBS, surface stained (α-CD4, 25 µg/ml), fixed in 2% paraformaldehyde and intracellular levels of 2-NBDG were quantified by flow cytometry.

### Cellular ATP levels

Cellular ATP was determined using a commercially available luminescent ATP detection kit (Abcam) per the manufacturer’s instructions. Briefly, 0.5 × 10^6^ CD4^+^ T cells were mixed with the kit’s detergent and ATP substrate followed by a 10 min incubation at RT. Samples were read using a luminescence microplate reader (Spectra Max M2, Molecular Devices, CA, USA). Absolute ATP concentrations were calculated using an ATP standard curve.

### GLUT-1 expression

Cellular GLUT1 expression was measured by intracellular staining. Briefly, CD4^+^ T cells were fixed and permeabilized using a commercially available kit (BD Biosciences) before intracellular staining with an anti-GLUT1 antibody (Abcam, ab195020, 1:100 dilution). CD4^+^ T cells were washed twice and MFI of anti-GLUT1 antibody was detected by flow cytometry.

### Quantitation of mitochondrial mass and mitochondrial viability/membrane potential

Cellular mitochondrial mass and potential were determined using commercially available probes MitoView Green (MTV-G; Biotium, CA, USA) and Mitotracker Deep Red (MTDR; Life Technologies, CA, USA), respectively. Antibody labeled (α-CD4) CD4^+^ T cells were fixed in 2% paraformaldehyde before MTV-G staining (50 nM) for 30 min at RT or CD4^+^ T cells were stained with MTDR (50 nm) for 20 min before fixation in 2% paraformaldehyde. MFI of MTV-G and MTDR^+^ CD4^+^ T cells was determined by flow cytometry.

### Proteomics

Cell preparations and extracted mitochondria were lysed in a buffer containing 5% SDS/50 mM triethylammonium bicarbonate (TEAB) in the presence of protease and phosphatase inhibitors (Halt; Thermo Scientific) and nuclease (Pierce™ Universal Nuclease for Cell Lysis; Thermo Scientific). Aliquots normalized to 65 µg protein (EZQ™ Protein Quantitation Kit; Thermo Scientific) were reduced with tris(2-carboxyethyl) phosphine hydrochloride (TCEP), alkylated in the dark with iodoacetamide, and applied to S-Traps (mini; Protifi) for tryptic digestion (sequencing grade; Promega) in 50 mM TEAB. Peptides were eluted from the S-Traps with 0.2% formic acid in 50% aqueous acetonitrile and quantified using Pierce™ Quantitative Fluorometric Peptide Assay (Thermo Scientific).

On-line high-performance liquid chromatography (HPLC) separation was accomplished with an RSLC NANO HPLC system [Thermo Scientific/Dionex: column, PicoFrit™ (New Objective; 75 μm i.d.) packed to 15 cm with C18 adsorbent (Vydac; 218MS 5 μm, 300 Å)], using as mobile phase A, 0.5% acetic acid (HAc)/0.005% trifluoroacetic acid (TFA) in water; and as mobile phase B, 90% acetonitrile/0.5% HAc/0.005% TFA/9.5% water; gradient 3 to 42% B in 120 min; flow rate at 0.4 μl/min. Data-independent acquisition mass spectrometry (DIA-MS) was conducted on an Orbitrap Fusion Lumos mass spectrometer (Thermo Scientific). A pool was made of all of the HPLC fractions (experimental samples), and 2 µg peptide aliquots were analyzed using gas-phase fractionation and 4-*m/z* windows (120k resolution for precursor scans, 30k for product ion scans, all in the Orbitrap) to create a DIA-MS chromatogram library [65] by searching against a panhuman spectral library [66]. Subsequently, experimental samples were blocked by replicate and randomized within each replicate. Injections of 2 µg of peptides were employed. MS data for experimental samples were also acquired in the Orbitrap using 12-*m/z* windows (staggered; 120k resolution for precursor scans, 30k for product ion scans) and searched against the DIA-MS chromatogram library generated. Scaffold DIA (ver. 1.3.1; Proteome Software) was used for all DIA data processing.

### Western blots

Cells or extracted mitochondria were lysed by sonication in RIPA lysis buffer (25 mM Tris-HCl, pH 7.6, 150 mM NaCl, 1% NP-40, 1% sodium deoxycholate, 0.1% SDS, ThermoFisher) containing Minitab protease inhibitors (Sigma). The total protein quantity per sample was determined by BCA assay. Proteins (20 µg/sample) were resolved on 4–15% Mini-PROTEAN® TGX™ Precast Protein Gels (Biorad) and transferred to PVDF membranes (Biorad). To quantify ETC proteins, membranes were incubated overnight with the Total OX-PHOS Rodent Western Blot Antibody Cocktail (Abcam; 1:500) in 1% milk at 4°C. To quantify isolated mitochondria purity, membranes were incubated with Cytochrome C (Cell signaling; 1:500), α-COX IV (Abcam, 1:500), and α-PCNA (Abcam; 1:500). Membranes were washed and incubated with mouse IgG kappa-binding protein conjugated to horseradish peroxidase (m-IgGk-HRP, 1:4000) (Santa Cruz technologies) in 5% milk for 1 h at 4°C. Membranes were then stripped and probed for β-actin levels. Images were captured on a UVP ChemStudio 815 system (Analytik, Jena, DE).

### Quantitation of intracellular thiols

Intracellular reduced glutathione levels were determined using commercially available fluorometric thiol probe thiol tracker Violet (Thermo Fisher) and the total glutathione (GSH) Detection Assay Kit (Abcam, MA, USA). CD4^+^ T cells were stained with thiol-tracker violet (10 µM) for 30 min at 37°C. Cells were washed twice with PBS, stained with surface antibodies of interest (*i.e.* α-CD3, α-CD4, and α-CD69; 25 µg/ml) for 20 min, fixed in 2% paraformaldehyde, and data collected by flow cytometry. The total GSH detection assay was performed according to the manufacturer’s directions.

### Cytokine detection

The EliSpot and Luminex assays were used to measure the cellular production of cytokines. For the EliSpot assay, 5.0 × 10^4^ CD4^+^ T cells were serially diluted onto either α-IL-2 or α-IFNɣ sterile coated PVDF 96 well plates (BD Biosciences, NJ, USA). The CD4^+^ T cells were then activated for 24 or 72 h in complete DMEM. After allotted time points, the EliSpot microplates were developed following the manufacturer’s protocol and analyzed using an ImmunoSpot analyzer (Cellular Technology Limited, OH, USA). Individual spots are representative of individual cytokine-producing cells.

Briefly, 2.5 × 10^5^ CD4^+^ T cells were cultured in 96 well plates containing PMA/Ionomycin for 24 or 72 h, after which media were collected and measured for the presence of cytokines IL-2, IFNɣ, IL-4, IL-5, IL-10, IL-17/IL-17A, TNFα, and the chemokine CCL5/RANTES following the manufacture’s protocol.

### Proliferation Assay

CD4^+^ T cell proliferation was determined by the Click-it EdU detection assay (Thermo Fisher) [67]. Briefly, 2.5 × 10^5^ CD4^+^ purified T cells were activated for 72 h in 600 µL cDMEM containing 50 µM 5-Ethynyl-2’-deoxyuridine (EdU) and αCD28 (0.05 µg/ml), in sterile 48 well plates precoated with 400 µL of αCD3 (0.5 µg/ml in PBS). EdU incorporation into DNA from proliferating cells was detected by a fluorescent bioconjugate and flow cytometry. The assay was performed per the manufacturer’s instructions.

### Adoptive transfer of naïve CD4^+^ T cells and infection of mice with influenza A or *M. tb*

Naïve CD4^+^ T cells from old mice were isolated by negative magnetic bead selection. Non-manipulated naïve CD4^+^ T cells or mito-transferred naïve CD4^+^ T cells (∼ 2.0 × 10^6^ cells) were tail vein injected into Rag1-KO (n= 4 to 5 mice per group) in a final volume of 200 µl. A control group of Rag1-KO (n= 2 to 4 mice per group) were tail-vein injected with 200 µl of PBS.

### Determination of influenza A-induced morbidity and mortality

Before and after infection with influenza A, mice were weighed and monitored daily for body weight change (% morbidity). Mice that lost 20% of their initial body weight were humanely euthanized (% mortality) [68, 69].

### Determination of *M. tb* bacterial burden

At 21 days post-*M.tb* infection, the lung (local infection), and spleen (systemic infection) of infected mice were excised and homogenized in sterile PBS (Sigma). Homogenized organs were serially diluted and plated onto 150 mm × 20 mm sterile plates of 7H11 agar supplemented with OADC (oleic acid, albumin, dextrose, catalase) and incubated for 14 to 21 days at 37°C. *M.tb* CFU was counted and expressed as Log_10_ CFU per organ [70, 71].

### Confocal Microscopy

-*Z*-stack confocal images were obtained on a Leica Stellaris-8 microscope (Leica Camera AG, Wetzlar, GER) equipped with an HC PL APO CS2 63x/1.40 OIL objective. Fluorophores were detected using UV laser diode 405 nm (for α-CD4 Pacific Blue; Biolegend) and solid-state lasers 488nm (for Turbo-GFP tagged hexokinase 1 mitochondria (HK-GFP-mito); Horizon Discovery, MA, USA) and 641 nm (RedDot™2 nuclear stain (RD-2); Biotium), with pinhole set to 5.5 AU, in XYZ scan mode, using a scan speed of 400 Hz, 1x Zoom, and 1.575 µs pixel dwell time. Post-processing was performed using the LAS X LIGHTNING ver.4.3.0.24308 (Leica Camera AG). Images were processed using ImageJ software.

### mt-DNA copy number

The quantification of mitochondrial DNA (mt-DNA) copy number in CD4^+^ T cells was done using the commercially available mouse mitochondrial DNA copy number kit (Detroid R&D, MI, USA), following the manufacturer’s instructions.

### Bioinformatic Analysis

Gene ontology analysis was performed using DAVID [72, 73]. The Bonferroni, Benjamini, and false discovery rate (FDR) were used for multiple test corrections[74]. Functional pathway and network analysis of differentially expressed proteins were performed using Ingenuity Pathway Analysis (IPA) (Qiagen, Redwood City, CA, USA). The pathway and network score were based on the hypergeometric distribution and calculated with the right-tailed Fisher’s exact test.

## 3. Results

### Mito-centrifugation delivers functional mitochondria to CD4^+^ T cells

Our initial experiments determined whether mito-centrifugation could deliver functional donor mitochondria to CD4^+^ T cells from old mice. Mitochondria isolated from MEFs (5.0 × 10^6^ cells) were pre-stained with Mitotracker Deep Red (MTDR) and transferred into CD4^+^ T cells isolated from old mice. Approximately 30 min after mito-transfer, CD4^+^ T cells were probed with MitoSOX Red and MTV-G, surface labeled with α-CD4, fixed in 4% PFA, and analyzed by flow cytometry. No more than 10% of CD4^+^ T cells from old mice were MTDR^+^ after mito-transfer indicating that these received donor mitochondria **(Fig. 1. A-C)**. Compared to non-manipulated CD4^+^ T cells from old mice (*i.e.* MTDR^-^ CD4^+^ T cells), the relative MFI of MitoSOX Red in the subset of CD4^+^ T cells from old mice that received mitochondria via mito-centrifugation significantly decreased, indicating a reduction in mitoROS production **(Fig. 1. D-E).** Further, compared to MTDR^-^ CD4^+^ T cells, the relative MFI of MTV-G in the subset of MTDR^+^ CD4^+^ T cells significantly increased, indicating an increase in mitochondrial mass **(Fig. 1. F).** These data suggested that mito-centrifugation transfered functional donor mitochondria to CD4^+^ T cells. Although the efficiency of this transfer at this volume was low (≤ 10%) **(Fig. 1. A-B),** these initial experiments hinted that mito-transfer significantly decreased mitoROS levels of CD4^+^ T cells in old mice. Next, we sought to improve and standardize a protocol for mito-transfer to CD4^+^ T cells.

**Fig. 1.**
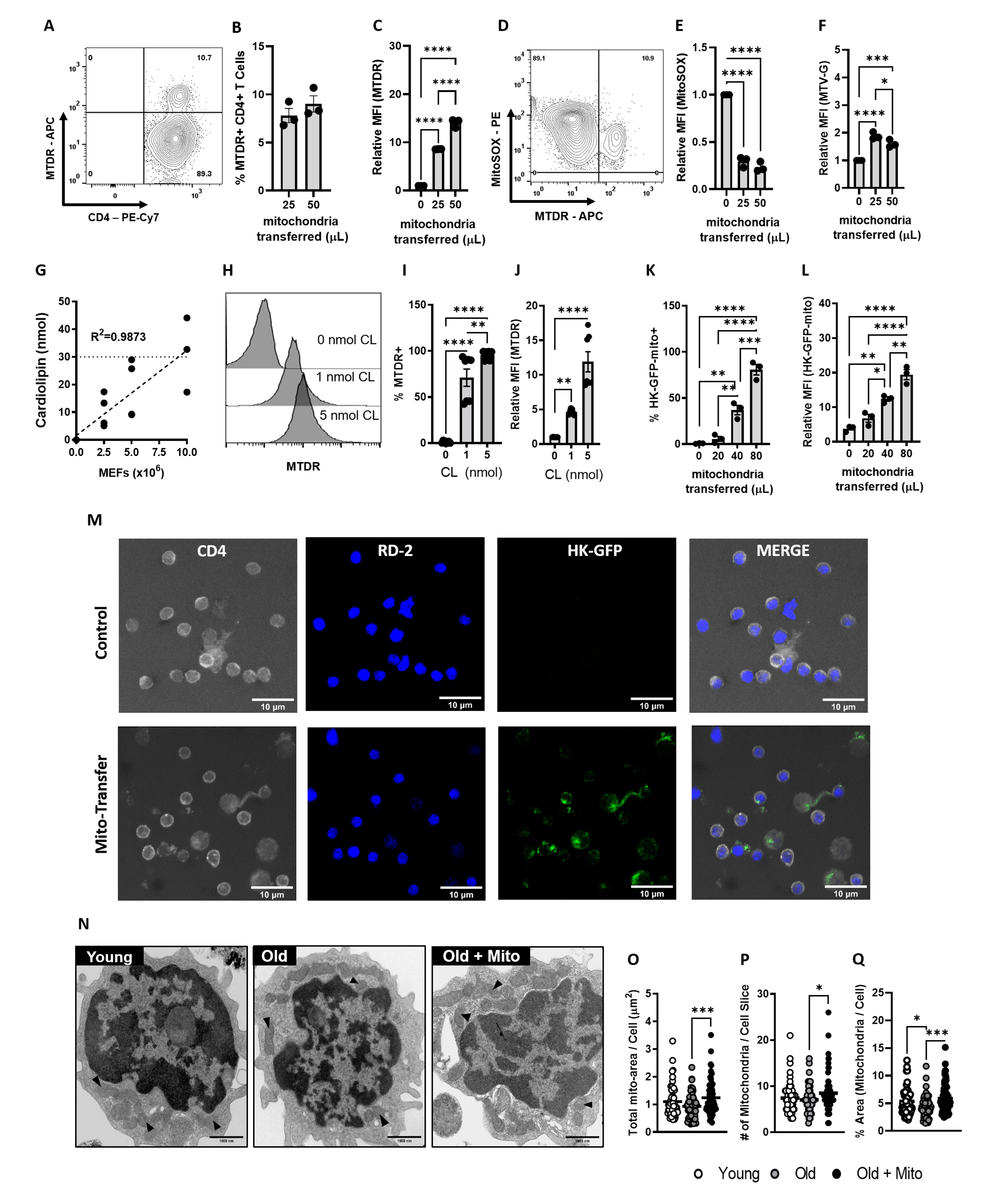
Mito-centrifugation efficiently delivers mitochondria to CD4^+^ T cells. (**A – F)** Donor mitochondria were prestained with Mito-tracker Deep Red (MTDR) and then transplanted into CD4^+^ T cells isolated from old mice via mito-centrifugation. After mito-transfer, cells were probed with MitoSOX to determine mitoROS production and MTV-G to determine relative mitochondrial mass. **A)** Representative Contour plot and scatter bar graphs of **B)** the percent MTDR^+^ CD4^+^ T cells and **C)** the relative fold change in MFI of MTDR^+^ CD4^+^ T cells after mito-transfer. **D)** Representative Contour plot and scatter bar graphs of **E)** relative MFI of MitoSOX Red and **F)** MTV-G in CD4^+^ T cells from old mice after mito-transfer. **G**) Cardiolipin concentration curve of isolated mitochondria from MEFs. Cardiolipin experiments were repeated twice with 3-4 mice/group. **H)** Flow cytometry histograms and scatter bar graphs of **I)** percent MTDR^+^ CD4^+^ T cells, and **J**) relative MFI of MTDR^+^ CD4^+^ T cells after mito-transfer, based on cardiolipin concentration. Flow cytometry scatter bar graphs of **K)** percent HK-GFP^+^ CD4^+^ T cells, and **L)** the relative MFI of HK-GFP^+^ CD4^+^ T cells after mito-transfer. **M)** Confocal images of CD4^+^ T cells with or without donor mitochondria (HK-GFP^+^). **N)** TEM images mitochondrial distributions (black arrows) in CD4^+^ T cells (young, old, and old+mito). Quantifications of **O)** total mitochondrial area, **P**) the number of mitochondria, and **Q)** percent area of mitochondria per cell area (in image slice) of CD4^+^ T cells isolated from young and old mice, and old mice ∼5-6 min after mito-transfer. TEM images were obtained from 2 mice/groups. All other experiments were done with at minimum 3 biological replicates (n=3). p ≤ 0.05 = *, p ≤ 0.01 = **, p ≤ 0.001 = *** and p ≤ 0.001 = ****, using unpaired Student’s *t*-test, or one-way-ANOVA where appropriate.

### Standardizing mito-transfer to CD4+ T cells from old mice

In eukaryotes, the phospholipid cardiolipin (CL) is exclusively localized in mitochondria [75-78], thus, for uniform cell cultures the abundance of CL is thought to accurately reflect the overall mitochondrial content [75, 78, 79]. Our results estimated that approximately 30 nmol of cardiolipin (equivalent to roughly 250 µg of extracted protein) is present in the mitochondrial fraction (*i.e.* supernatant) isolated from 1.0 × 10^7^ MEFs **(Fig. 1. G)**. We also assessed whether non-mitochondrial proteins were present in the mitochondrial fraction by detecting PCNA, a nuclear localized protein, PCNA, via western blot. PCNA was present in the mitochondrial fraction, but was comparatively lower than the cytoplasmic fraction that remained after mitochondria isolation **(Supp. Fig. 1)**. These data suggest that the mitochondrial fraction was enriched for mitochondria.

To further optimize mito-transfer efficiency, two volumes of MEF mitochondria based on CL concentrations (1 nmol and 5 nmol CL), were mito-centrifuged into CD4^+^ T cells. MEF were stained with MTDR before isolation and transfer. After mito-transfer, whole CD4^+^ T cell populations were analyzed by flow cytometry to quantify frequency of MTDR^+^ CD4^+^ T cells, and shifts in MTDR fluorescence (MFI), which are indicative of successful mito-transfer. Both concentrations tested significantly increased the frequencies of MTDR^+^ CD4^+^ T cells and shifted the MFI of MTDR (**Fig. 1. H-J**). The MFI of MTDR in CD4^+^ T cells increased by 5 fold with 1 nmol of cardiolipin, while 5 nmol increased the MFI by 10 fold (**Fig. 1. J**). However, maximum (100%) mito-transfer efficiency was achieved with 5 nmol CL (**Fig. 1. I**);which was used for subsequent experiments.

We further confirmed successful delivery of mitochondria to CD4^+^ T cells by quantification and visualization of fluorescent mitochondria (Turbo-GFP tagged hexokinase1; HK-GFP-mito), and transmission electron microscopy (TEM). Various volumes of HK-GFP-mito (0, 20, 40, 80 µL) were mito-centrifuged into CD4^+^ T cells. The percent of HK-GFP-mito^+^ CD4^+^ T cells increased with higher HK-GFP-mito volume **(Fig. 1.K-L)**, indicating increased delivery of functional mitochondria. These data paralleled our initial experiments with MTDR prestained mitochondria **(Fig 1.B, I)**. We further confirmed whether the GFP signal originated from within CD4^+^ T cells. A 3D volumetric reconstruction of confocal slices (*Z-stacks*) of CD4^+^ T cells counterstained with αCD4 surface marker and RedDot™2 (RD-2) nuclear stain, indicated that HK-GFP-mito signal was from within the cytosolyc compartment of recipient CD4^+^ T cells after mito-centrifugation **(Fig.1. M)**.

Immediately after mito-transfer (∼5 min), mito-transferred CD4^+^ T cells from old mice, and non-manipulated CD4^+^ T cells from young and old mice (controls) were fixed and processed for TEM. Black triangles in TEM micrographs are representative of individual mitochondria **(Fig. 1.N)**. Compared to CD4^+^ T cells from young mice, the total mitochondrial area **(Fig. 1.O)** and the number of mitochondria **(Fig. 1.P)** per CD4^+^ T cell micrograph from old mice trended towards a decrease but it was not significant. However, the % ratio of mitochondrial area to cell area **(Fig. 1.Q)** was significantly lower in CD4+ T cells from old mice compared to CD4 T cells from young mice (4.38% vs. 5.33%). After mito-transfer was performed on CD4^+^ T cells from old mice, there were significant increases in the total mitochondrial area (from 0.92 ± 0.06 to 1.24 ± 0.06 µm^2^), the number of mitochondria (from 7.03 ± 0.35 to 8.46 ± 0.46), and % ratio of mitochondria to cell area (from 4.38 ± 0.24 to 5.88 ± 0.28). Collectively, our flow cytometry, confocal microscopy and TEM data indicate that mitochondria were succeffully delivered into CD4^+^ T cells from old mice.

### Mito-transfer decreases mitochondrial oxidative stress in CD4^+^ T cells from old mice

Observations that mito-transfer of MEF mitochondria decreased mitoROS levels in CD4^+^ T cells from old mice (**Fig. 1.D-E**) warranted further examination of the oxidative stress status of CD4^+^ T cells from old mice prior to and after mito-transfer. The expression of global antioxidant proteins and proteins related to mitochondrial detoxification were detected by LC-MS/MS in non-manipulated and mito-transferred CD4^+^ T cells from old mice (i.e. **Fig.2. A, O1-O3**), at 4 h after mito-transfer. Compared to non-manipulated CD4^+^ T cells from old mice, a majority of cellular antioxidants and mitochondrial detoxification proteins were downregulated in mito-transferred CD4^+^ T cells from old mice (**Fig. 2.A**, **OM1-OM3**, and **Supp. Table 1**). Flow cytometry-based detection of total thiol levels, as indicators of sulfate-based oxidants, showed no significant changes in CD4^+^ T cells from old mice with or without mito-transfer, as well as CD4^+^ T cells from young mice **(Fig. 2.B).**

**Fig 2.**
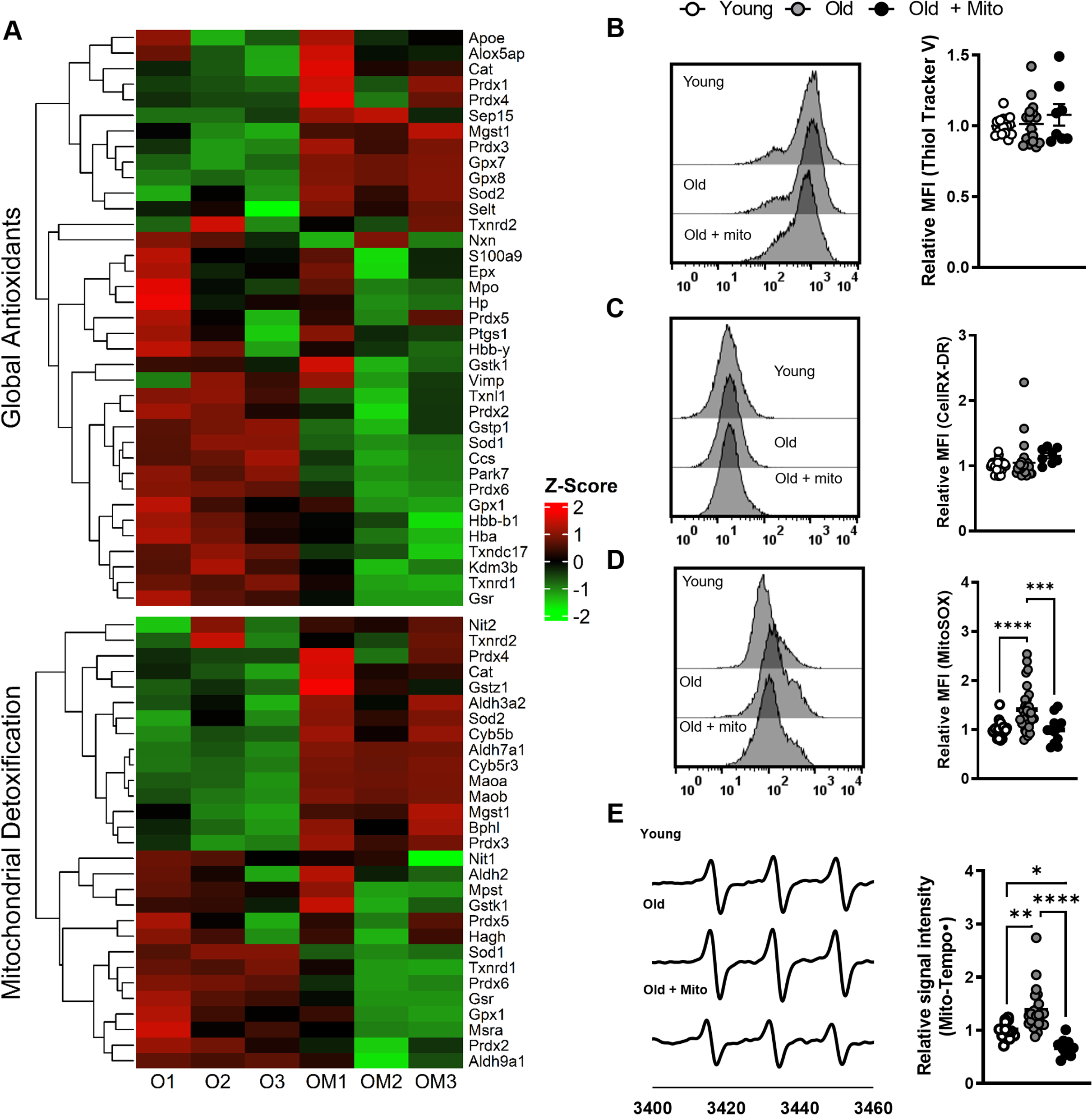
Mito-transfer decreases oxidative stress in CD4^+^ T cells from old mice. **A)** Heatmap of proteins related to cellular antioxidants and mitochondrial detoxification in CD4^+^ T cells from old mice with and without mito-transfer (4h). Flow cytometry histograms and scatter plots of the relative MFI of **B)** thiol-tracker violet, **C)** CellROX Deep Red, and **D)** MitoSOX Red stained CD4^+^ T cells from young, old mice and CD4^+^ T cells from old mice after mito-transfer. **E**) EPR spectrum and H) bar graphs of CD4+ T cells probed with Mito-Tempo-H. 4-5 mice per group, with experiments repeated at least once, p ≤ 0.05 = *, p ≤ 0.01 = **, p ≤ 0.001 = *** and p ≤ 0.001 = ****, using one-way-ANOVA where appropriate.

Basal levels of cytoplasmic (cytoROS) and mitochondrial (mitoROS) in old CD4^+^ T cells was also assessed at 4 h after mito-transfer and compared against non-manipulated CD4^+^ T cells from young and old mice. No significant difference was detected in cytoROS (relative MFI of CellROX DR) in CD4^+^ T cells **(Fig. 2.C)**. However, mitoROS levels (relative MFI of MitoSOX Red) in mito-transferred CD4^+^ T cells from old mice, and non-manipulated CD4^+^ T cells from young mice were similar, and both were significantly lower than mitoROS levels in non-manipulated CD4^+^ T cells from old mice **(Fig. 2.D)**. EPR-based detection of Mito-TEMPO• in CD4^+^ T cells also showed similar results, corroborating results from MitoSOX Red experiments **(Fig. 2.E)**. Overall, these data suggest that mito-transfer reduced the basal levels of mitoROS in CD4^+^ T cells from old mice to those measured in CD4^+^ T cells from young mice without affecting cytoROS levels.

### Mito-transfer alters mitochondrial gene expression and mitochondrial ultrastructure

Whether mito-transfer changed the mitochondrial proteome of CD4^+^ T cells from old mice was investigated through pathway analysis based on the cellular proteome detected by LC-MS/MS. The expression of the majority of proteins in the mitochondrial proteome increased in mito-transferred CD4^+^ T cells from old mice **(Fig. 3.A, Supp. Table 2.)**. Ingenuity Pathway Analysis (IPA) revealed that the most upregulated pathways were related to mitochondrial function **(Fig. 3.B).** Similar to IPA results, the most upregulated processes identified by Gene Ontology Term enrichment analysis were related to mitochondrial function and ultrastructure **(Fig. 3.C).** To corroborate these findings, and understanding that mitochondria function relies on the inner membrane, we compared the mitochondrial cristae ultrustructure of mito-transferred CD4^+^ T cells against non-manipulated CD4^+^ T cells from old mice, at 0.1 h and 4 h after mito-transfer **(Fig. 3.D)**. Compared to non-manipulated CD4^+^ T cells, mito-transferred CD4^+^ T cells from old mice had denser mitochondrial cristae evidenced by increased and more pronounced folds/wrinkles **(Fig. 3.D; black arrows)**, indicative of increased inner mitochondrial membrane surface area.

**Fig 3.**
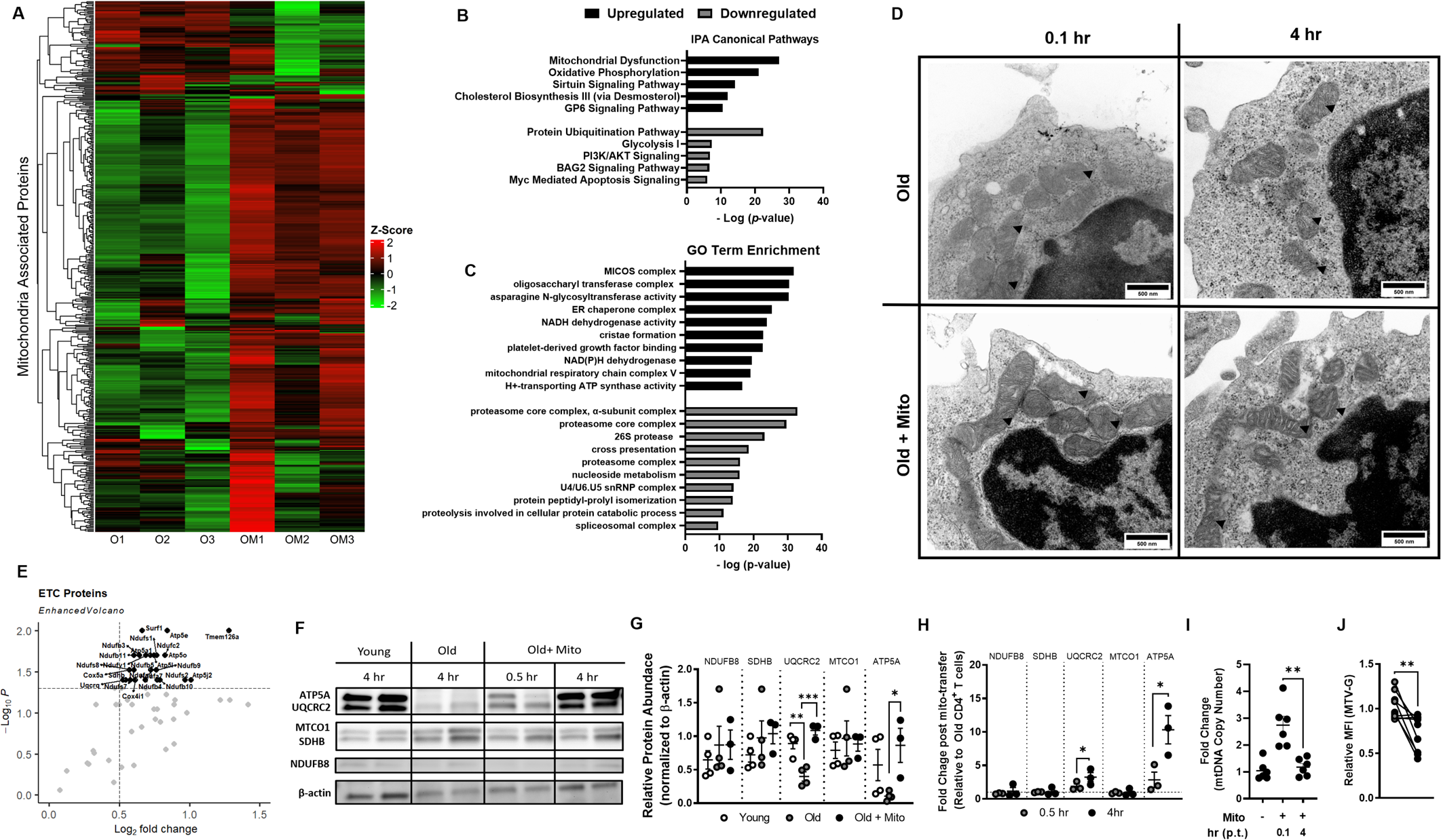
Mito-transfer alters mitochondrial gene expression and mitochondrial ultrastructure. **A)** Heatmap of the mitochondrial proteome in CD4^+^ T cells from old mice with and without mito-transfer. **B)** Top 5 upregulated and downregulated Canonical pathways (IPA) and the **C)** top 10 upregulated and downregulated GO Term enrichment of old CD4+ T cells after mito-transfer**. D)** TEM images of mitochondrial cristae morphology in CD4^+^ T cells from old mice with and without mito-transfer at ∼5 mins and 4 h after mito-transfer. **E)** Volcano plot of electron transport chain (ETC) proteome detected in CD4^+^ T cells from old mice with and without mito-transfer, 4 h after mito-transfer. **F)** Western blot images and, **G)** relative protein abundance in CD4^+^ T cells isolated from young and old mice, and CD4^+^ T cells from old mice 4 h after mito-transfer. **H)** Fold change in ETC proteins in CD4^+^ T cells from old mice after mito-transfer (at 30 min and 4 h). Proteomic experiments were performed once with 3 mice/group and internally normalized to β-actin. **I)** Fold change in mtDNA copy number in CD4^+^ from old mice after mito-transfer (at 5 min and 4 h). **J)** Relative MFI of MTV-G in CD4^+^ T cells from old mice (grey) and CD4^+^ T cells from old mice (4h) after mito-transfer. p ≤ 0.05 = * or p ≤ 0.01 = **, using paired and unpaired Students’ *t*-test where appropriate.

The subset of mitochondrial proteome related to the electron transport chain (ETC) of CD4^+^ T cells from old mice with and without mito-transfer was also analyzed. After 4 h of mito-transfer, CD4^+^ T cells from old mice showed significant upregulation (1.5-fold increase in protein expression in mitochondria-transferred old *vs.* old, with adjusted *p*-value <0.1) of proteins related to complexes of the ETC, such as NDUS2, SDHA, COX15, QCR7, ATPK **(Fig. 3.E, Supp. Table 3).** We further examined the expression of selected ETC proteins important at different steps of Ox-Phos (NDUFB8, SDHB, UQCRC2, MTCO1, and ATP5A) in old mouse CD4^+^ T cells at 30 min and 4 h after mito-transfer, and compared their relative expression against those of CD4^+^ T cells isolated from young and old mice **(Fig. 3.F-H)**. Compared to CD4^+^ T cells from young mice, significantly lower levels of UQCRC2, a subunit of CIII of the ETC, and lower levels of ATP5A, a protein subunit of the ATP synthase machinery, were detected in old mouse CD4^+^ T cells **(Fig. 3.G-H)**. However, after 4 h of mito-transfer, both ATP5A and UQCRC2 were significantly upregulated in CD4^+^ T cells from old mice **(Fig. 3.F-H)**. No other differences were noted between the detected ETC proteins of CD4^+^ T cells from all groups studied. These observed changes in mitochondrial proteome prompted us to consider the impact of mito-transfer on the mtDNA in CD4^+^ T cells from old mice after mito-transfer. indeed, immediately after mito-transfer, mtDNA copy number significantly increased in CD4^+^ T cells from old mice and returned to previous levels 4 h after **(Fig. 3.I)**.

To determine the impact on mitochondrial mass, T cells were fixed and then stained with MTV-G at 4 h post mito-transfer. Compared to non-manipulated CD4^+^ T cells from old mice, mito-transferred CD4^+^ T cells from old mice had significantly lower MFIs of MTV-G, indicating a decrease in overall mitochondrial mass **(Fig. 3.J)**. Collectively, these data suggest that mito-transfer altered the mitochondrial proteome, ultrastructure, and decreased overall mitochondrial abundance.

### Mito-transfer increases aerobic metabolism in CD4+ T cells from old mice

Extracellular flux analysis (mito-stress test) was used to assess changes in cellular oxygen kinetics (OCR) of non-manipulated and mito-transferred CD4^+^ T cells from old mice. The OCR of CD4^+^ T cells from young mice was also measured as a reference **(Fig. 4.A)**. Compared to CD4^+^ T cells from young mice, CD4^+^ T cells from old mice had increased maximal respiration, increased proton leak and decreased coupling efficiency **(Fig. 4.A-D)**. No other significant differences were noted in other OCR parameters measured in CD4^+^ T cells from young and old mice **(Fig. 4.A-D)**. However, mito-transferred CD4^+^ T cells from old mice had significantly increased basal, maximal, spare respiratory capacity **(Fig. 4.B)**, proton leak, non-mitochondrial oxygen consumption rates, and ATP production **(Fig. 4.C)**. Further, the coupling efficiency of mito-transferred CD4^+^ T cells from old mice decreased, indicating an increase in proton leak while maintaining ATP production **(Fig. 4.D)**. ECAR of CD4^+^ T cells from old mice was significantly higher than that of CD4^+^ T cells from young mice, and mito-transfer further enhanced ECAR of CD4^+^ T cells from old mice **(Supp. Fig. 2. A-B)**. Therefore, mito-transfer increased the rates of Ox-Phos in CD4^+^ T cells from old mice.

**Fig. 4.**
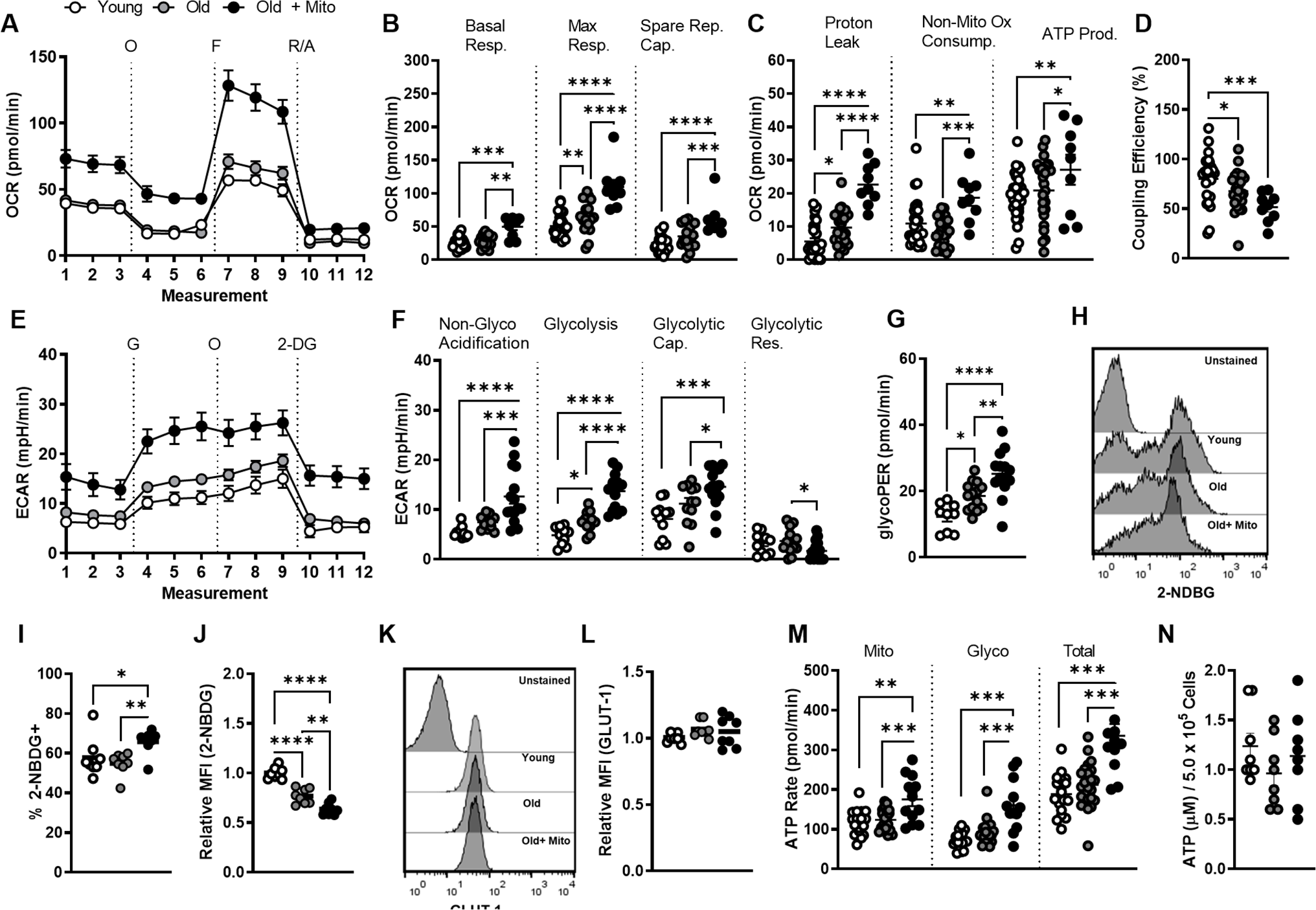
Mito-transfer increases aerobic metabolism in CD4^+^ T cells from old mice. CD4^+^ T cells (2.5 × 10^5^) were plated onto poly-D-lysine coated 96-well microplates, after which O_2_ consumption (OCR) and extracellular acidification rates (ECAR) were measured under basal conditions and in response to the mito-stress test. **A)** Representative mito stress test kinetic graph, **B)** quantitation of basal, maximal, and spare respiratory capacity, **C)** proton leak, non-mitochondrial oxygen consumption, and ATP production rate, and **D)** coupling efficiency. **E)** Representative glycolysis stress test kinetic graph, and quantitations of **F)** non-glycolytic acidification, glycolysis, glycolytic capacity, glycolytic reserve and **G)** GlycoPER of CD4^+^ T cells. **H)** Flow cytometry histograms and **I-J)** dot plots of 2-NBDG uptake in CD4^+^ T cells from young and old mice, and CD4^+^ T cells from old mice after mito-transfer. **K)** Flow cytometry histograms and **L)** dot plots of GLUT1 expression in CD4^+^ T cells from young and old mice, and CD4^+^ T cells from old mice after mito-transfer. **M)** Rates of glycolytic and mitochondrial-derived ATP, and total ATP production, and **N)** cellular levels of ATP in CD4^+^ T cells from young and old mice, and CD4^+^ T cells from old mice after mito-transfer; 3 to 4 mice were used per group and experiments were repeated at least once (n ≥ 2), p ≤ 0.05 = *, p ≤ 0.01 = **, p ≤ 0.001 = *** and p ≤ 0.001 = ****, using one-way-ANOVA.

### Increased glycolysis in CD4+ T cells from old mice after mito-transfer

Both glycolysis and mitochondrial respiration (β-oxidation, ATPase activity) can contribute to cellular ECAR [80]. We delineated the observed changes in ECAR by performing the glycolysis stress test **(Fig. 4.E)**. Basally, CD4^+^ T cells from old mice were more glycolytic than CD4^+^ T cells from young mice and had increased non-glycolytic acidification **(Fig. 4.F)**. Mito-transfer further significantly increased glycolysis, non-glycolytic acidification, and the glycolytic capacity of CD4^+^ T cells from old mice **(Fig. 4.F)**. No significant differences in the glycolytic reserve were observed between CD4^+^ T cells from young and old mice. However, after mito-transfer, CD4^+^ T cells from old mice had significantly lower glycolytic reserves **(Fig. 4.F)**.

The glycolytic rate assay and 2-NDBG glucose analog uptake were also performed to further corroborate results from the glycolysis stress test. The proton efflux rate (PER) obtained from the glycolytic rate assay is a measure of glycolytic flux (glucose →pyruvate). Compared to CD4^+^ T cells from young mice, CD4^+^ T cells from old mice had significantly higher glycoPER, which further increased in mito-transferred CD4^+^ T cells from old mice **(Fig. 4.G).** For 2-NDBG tracing, CD4^+^ T cells were incubated in DMEM containing 2-NDBG, afterwhich the percent and relative MFIs of 2-NDBG^+^ cells were quantified. CD4^+^ T cells from old and young mice had a similar percent of 2-NDBG^+^ populations **(Fig. 4.H-I)**. After mito-transfer, the percentage of 2-NDBG^+^ CD4^+^ T cells significantly increased **(Fig. 4.I)**. These data suggest that mito-transfer increased the glucose uptake of CD4^+^ T cells in old mice. Further, the relative MFI of 2-NDBG^+^ CD4^+^ T cells from young mice was significantly higher than that of CD4^+^ T cells from old mice **(Fig. 4.J)**. Mito-transfer further decreased the MFI of 2-NDBG in CD4^+^ T cells from old mice **(Fig. 4.J)**. This decrease in MFI suggests faster degradation of 2-NDBG, and increased glycolytic rate, which is in agreement with the glycoPER data [81, 82].

We additionally assessed the expression of GLUT-1 in CD4^+^ T cells after mito-transfer but observed no significant change **(Fig. 4.K-L)**. Collectively, data from seahorse and flow cytometry-based assays suggest that mito-transfer increased glycolysis in CD4^+^ T cells from old mice.

### Increased ATP production in CD4+ T cells from old mice after mito-transfer

Increased glycolysis and oxygen consumption hinted of an increase in the synthesis of ATP [83, 84]. Through the XF ATP rate assay, we determined whether mito-transfer increased ATP synthesis in CD4^+^ T cells from old mice, and what were the relative contributions of glycolysis and mitochondrial Ox-Phos. CD4^+^ T cells from young and old mice had comparable rates of mitochondria-linked ATP production (**Fig. 4.M**). Compared to young mice, CD4^+^ T cells from old mice showed higher rates of glycolysis-linked ATP production (**Fig. 4.M**). Mito-transfer enhanced both mitochondria and glycolysis-linked ATP production rates of CD4^+^ T cells from old mice, with a roughly 50% increase in total ATP production (**Fig. 4.M**). We additionally measured the intracellular levels of ATP but noted no significant differences among groups (**Fig. 4.N**).

### Mito-transfer improves T cell activation and cytokine production in CD4+ T cells from old mice

Because mitochondrial fitness impacts T cell activation, we compared TCR signalosome and phosphorylation kinetics of CD4^+^ T cells from old mice with and without mito-transfer. LC-MS/MS analysis showed that mito-transfer basally downregulated a majority of TCR signalosome proteins in CD4^+^ T cells from old mice **(Fig. 5.A-B, Supp. Table 4).** To examine the activation-induced T cell phosphorylation kinetics, CD4^+^ T cells were stimulated with αCD3/αCD28 for either 15, 30, or 60 min, fixed with 2% PFA and intracellularly stained for phosphorylated proteins (p-SLP76, p-PLCg1, p-ZAP70, and p-LCK). Compared to non-manipulated CD4^+^ T cells from old mice, the activation-induced phosphorylation of TCR signalosome proteins p-SLP76 and p-PLCg1 were significantly higher in mito-transferred CD4^+^ T cells from old mice **(Fig. 5.C-D)**, but no significant differences in the phosphorylation kinetics of p-ZAP70, and p-LCK were detected **(Fig. 5.E-F).**

**Fig 5.**
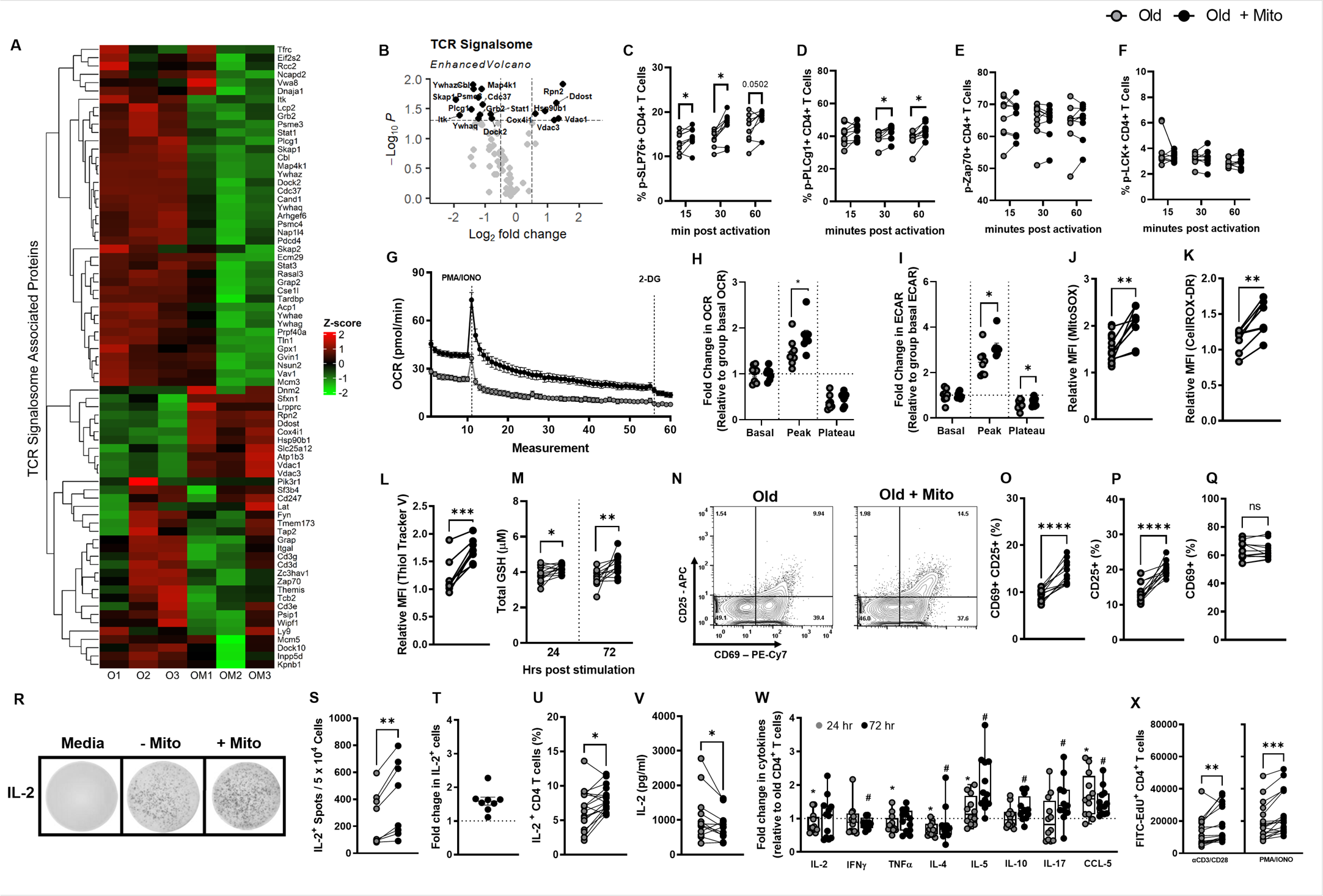
Mito-transfer improves T cell activation and cytokine production in CD4^+^ T cells from old mice. The CD4^+^ T cells from old mice with and without mito-transfer were evaluated for activation capabilities. **A)** Heatmap and **B)** volcano plot of TCR signalosome proteome detected in CD4^+^ T cells isolated from old mice with or without mito-transfer. The phosphorylation kinetics of TCR-related proteins **C)** SLP76, **D)** PLCg1, **E)** Zap70 and **F)** LCK in CD4^+^ T cells from old mice with or without mito-transfer, at 15, 30, and 60 min post activation with αCD3/αCD28 stimulation. **G)** OCR micrograph of CD4^+^ T cells from old mice with or without mito-transfer, after acute activation with PMA/IONO. The relative fold changes in **H)** OCR and **I)** ECAR of CD4^+^ T cells isolated from old mice with or without mito-transfer after acute activation with PMA/Ionomycin. The relative MFI of **J)** MitoSOX Red and **K)** CellROX Deep Red stained CD4^+^ T cells from old mice with or without mito-transfer, after stimulation PMA/ Ionomycin (4h). **L)** Thiol-tracker violet and the **M)** total GSH produced by CD4^+^ T cells from old mice with or without mito-transfer after stimulation with PMA/IONO.**N)** Contour plots of CD25 v CD69 expression on CD4^+^ T cells with and without mito-transfer, after PMA/ Ionomycin stimulation (24 h). The percent of **O)** CD25^+^ CD69^+^, **P)** CD25^+^ and, **Q)** CD69^+^ CD4^+^ T cells from old mice with or without mito-transfer, at stimulation with PMA/ Ionomycin (24h). **R)** Representative EliSpot images, the **S)** number IL-2^+^ producing T cells and the **T)** fold change in IL-2^+^ producing CD4^+^ T cells from old mice with or without mito-transfer, after stimulation with PMA/Ionomycin (24hr). The **U)** percent of IL-2^+^ CD4^+^ T cells via intracellular cytokine staining, in CD4^+^ T cells from old mice with or without mito-transfer, after stimulation with PMA/Ionomycin (4h). **V)** Soluble IL-2 detected in the supernatants of PMA/Ionomycin-stimulated (24 h) CD4^+^ T cells from old mice with or without mito-transfer. The relative fold change in cytokine production of PMA/Ionomycin-stimulated CD4^+^ T cells from old mice with or without mito-transfer, at 24 & 72 h after stimulation. **X)** Number of EdU^+^ cells in αCD3/αCD28 or PMA/Ionomycin-stimulated CD4^+^ T cell cultures from old mice with or without mito-transfer. All experiments were at least repeated (n=2), p ≤ 0.05 = *, p ≤ 0.01 = **, or p ≤ 0.001 = *** using paired Student’s *t*-test.

T cell activation-induced switch from Ox-Phos to aerobic glycolysis is concurrent with TCR signalosome phosphorylation events. Since mito-transfer increased basal Ox-Phos in CD4^+^ T cells from old mice, we questioned how mito-transfer impacted the ability of CD4^+^ T cells from old mice to undergo an activation-induced switch from Ox-Phos to aerobic glycolysis. The basal OCR of mito-transferred CD4^+^ T cells from old mice was significantly higher than that of non-manipulated CD4^+^ T cells from old mice **(Fig. 5.G)**. These data are in agreement with our OCR data (mito-stress test, **Fig. 4.A-B**). After acute injection of PMA/IONO, the peak OCR values of mito-transferred CD4^+^ T cells were significantly higher than those of non-manipulated CD4^+^ T cells from old mice **(Fig. 5.H)**. Similar trends were noted for ECAR measurements of mito-transferred CD4^+^ T cells from old mice **(Fig. 5.I)**. Collectively, these data suggest that mito-transferred CD4^+^ T cells more rapidly switched from Ox-Phos to aerobic glycolysis.

TCR-induced activation also increases ROS production in CD4^+^ T cells. Compared to non-manipulated CD4^+^ T cells from old mice, mito-transferred CD4^+^ T cells had significantly higher levels of mitoROs (mitoSOX fluorescence, **Fig. 5.J)**, and levels of cytoROS (CellROX DR fluorescence, **Fig. 5.K)** after activation. These data suggest that mito-transfer improved TCR signaling during T cell activation. Cellular antioxidant levels are important to counterbalance ROS production during T cell activation. Thus, the relative levels of intracellular thiols and total GSH in the supernatant of activated CD4^+^ T cells from old mice with or without mito-transfer were compared. The relative MFI of thiol-tracker violet increased by roughly 50% in mito-transferred CD4^+^ T cells from old mice **(Fig. 5.L).** The total GSH produced by T cells was determined at 24 h and 72 h post-activation. Compared to CD4^+^ T cells without mito-transfer, at both timepoints, mito-transferred CD4^+^ T cells from old mice produced significantly higher amounts of GSH **(Fig. 5.M).**

To further examine whether mito-transfer would improve T cell activation, the expression of CD69 and CD25, common markers of T cell activation, were determined 24 h after mitogenic stimulation. Indeed, mito-transfer increased the number of CD25^+^CD69^+^ activated cells by approximately 50% **(Fig. 5.N-O**). Overall, mito-transfer increased the amount of CD4^+^ T cells from old mice that expressed CD25^+^ (**Fig. 5.P**), but did not change the proportion of total CD69^+^ CD4^+^ T cells after activation (**Fig. 5.Q**).

Previous studies highlight the involvement of mitoROS, specifically H_2_O_2_, in modulating IL-2 expression in T cells [15]. The *ex vivo* stimulation of mito-transferred CD4^+^ T cells from old mice led to increased expression of CD25, which is also the high affility IL-2 receptor (IL-2Rα) [85, 86]. Mito-transfer increased the amount of IL-2^+^ spots (counted as individual spots) by approximately 50% (**Fig. 5.R-T**). Intracellular IL-2 cytokine staining and flow cytometry analysis further corroborated our EliSPOT results **(Fig. 5.U).**

To determine whether the impact of mito-transfer extended beyond IL-2 production or T helper cell lineage, we additionally measured levels of soluble IL-2 and other cytokines (IFNχ, TNFα, IL-4, IL-5, IL-10, IL-17) and the chemokine CCL-5/RANTES, in culture media of activated non-manipulated and mito-transferred CD4^+^ T cells at 24 h and 72 h post-activation **(Fig. 5.V-W)**. The levels of IL-2 in the media of mito-transferred CD4^+^ T cells from old mice were significantly lower than that of non-manipulated CD4^+^ T cells from old mice at 24 h post-activation **(Fig. 5. V-W).** The media of mito-transferred CD4^+^ T cells from old mice also contained significantly lower levels of cytokines TNFα and IL-4, significantly higher levels of IL-5, and CCL-5/RANTES, and comparable levels of IFNγ, IL-10, and IL-17 at 24 h post activation. At 72 h post-infection, these cytokine/ chemokine levels shifted with significantly lower IFNγ and IL-4, significantly higher IL-5, IL-10, IL-17a, and CCL-5/RANTES, and comparable IL-2 and TNFα levels (**Fig. 5.V-W**). These data suggest that mito-transfer in old CD4^+^ T cells impacts broad cytokine production and extends beyond 24 h of activation.

Finally, whether the proliferative capacity of CD4^+^ T cells from old mice improved after mito-transfer was also examined. CD4^+^ T cells from old mice were stimulated for 72 h in complete media supplemented with 50 µM EdU, which is incorporated into newly synthesized DNA during cell proliferation. Compared to non-manipulated CD4^+^ T cells from old mice, CD4^+^ T cells after mito-transfer had a two-fold increase in the number of Edu^+^ cells **(Fig. 5.X).**

### Mito-transfer in naïve CD4 T cells from old mice improves the control of infection

Previous studies suggest that compared to other T cell subpopulations, naïve CD4^+^ T cells are most impacted by aging-associated dysregulation and immunosenescence [7, 9, 84, 85], and their dysfunction drives susceptibility to several microbial infections in the elderly. To further assess this, the *in vivo* function of mito-transferred naïve CD4^+^ T cells was evaluated in separate Rag1-KO adoptive transfer models of influenza A virus (IAV, acute) and *Mycobacterium tuberculosis* (*M.tb*, chronic) infections **(Fig. 6.A)**.

**Fig 6.**
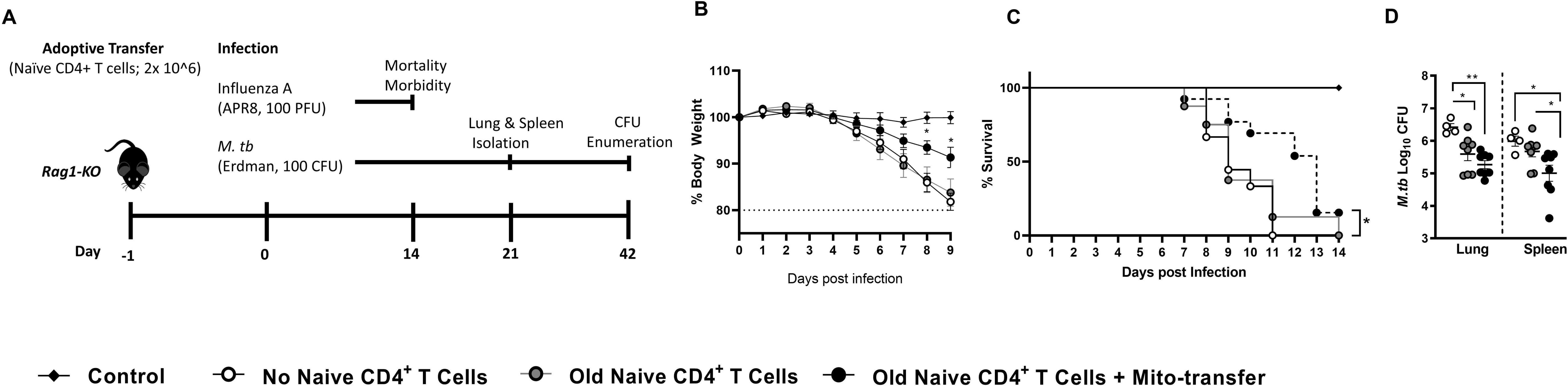
Mito-transfer in naïve CD4 T cells from old mice protects mice against pathogens & promotes protective T cell phenotypes. **A)** 2.0 × 10^6^ naïve CD4^+^ T cells from old mice with or without mito-transfer, or PBS were tail vein injected in Rag1-KO mice that were subsequently infected with either IAV or *M.tb*. Rag1-KO mice infected with IAV were monitored for **B)** disease body weight (morbidity) and **C)** survival (mortality). After 21 days of infection, **D)** lung and spleen of Rag1-KO mice infected with *M.tb* were processed to determine M.tb burden.

Compared to Rag1-KO mice that received either PBS or non-manipulated naïve CD4^+^ T cells from old mice, Rag1-KO mice that received mito-transferred naïve CD4^+^ T cells from old mice had a significant delay in IAV-induced morbidity (weight loss) by day 8 post-infection **(Fig. 6.B)**. Rag1-KO mice that received mito-transferred naïve CD4^+^ T from old mice also had significantly reduced mortality (13 days median survival), compared to mice that received either PBS or non-manipulated naïve old mouse CD4^+^ T cells [9 days median survival **(Fig. 6. C)**]. Overall, these data suggest that mito-transferred naïve CD4^+^ T cells from old mice improved control of infection (morbidity and mortality) over non-manipulated naïve CD4^+^ T cells from old mice during IAV infection.

In an *M.tb* model of infection, fewer *M.tb* colony forming units (CFUs) were observed in the lungs of Rag1-KO mice that received mito-transferred naïve CD4^+^ T cells (*p* = 0.004) or non-manipulated naïve CD4^+^ T cells from old mice (*p* = 0.01), compared to control infected Rag1-KO mice **(Fig. 6.D)**. There were, however, no significant differences between the *M.tb* burden in the lung of Rag1-KO mice that received mito-transferred naïve CD4^+^ T cells and non-manipulated naïve CD4^+^ T cells from old mice **(Fig. 6.D)**. The *M.tb* burden in the spleen of Rag1-KO mice that received mito-transferred naïve CD4^+^ T cells was significantly lower than that of Rag1-KO mice that received non-manipulated naïve CD4^+^ T cells from old mice (*p* = 0.016) or PBS (control, *p* = 0.019) **(Fig. 6.D)**. There were no significant differences in *M.tb* burden in the spleen of Rag1-KO mice that received non-manipulated naïve CD4^+^ T cells or PBS **(Fig. 6.D)**. *M.tb* studies were repeated using half the amount of adoptively transferred cells (1.0 × 10^6^) and while similar trends were observed, only the lungs of mice that received mito-transferred naïve CD4^+^ T cells from old mice had a statistically lower *M.tb* burden (p = 0.002) than control mice that received PBS or were infected with *M.tb*. **(Supp. Fig. 5)**. These data suggest that mito-transferred naïve T cells from old mice are capable of developing effector responses and responding to *M.tb* infection but on a Rag1-KO background, absent other essential immune cells, these responses had minimal effect.

## 4. Discussion

Mitochondria are indispensable for proper T cell homeostasis [13, 15, 17, 20, 27], and aging-associated decline in adaptive immunity is linked to mitochondrial dysfunction and redox imbalance [6, 17, 87-89]. The restoration of redox balance through the reconstitution of mitochondrial fitness (mito-transfer) in lymphocytes from old mice and the elderly humans has not been thoroughly examined. Our results support our hypothesis that restoration via mito-transfer would improve immunological CD4^+^ T cell functions.

In our experiments using CD4^+^ T cells from old mice, mito-transfer altered the basal levels of mitoROS and a majority of antioxidant proteins, shifting cells towards lower basal levels of intracellular ROS. Proteomic analysis (IPA and GO term enrichment) of mito-transferred CD4^+^ T cells from old mice showed that the most upregulated cellular pathways and processes were related to mitochondrial remodeling and Ox-Phos. Visually, this was corroborated by qualitative EM images, where the cristae ultrastructure of mito-transferred CD4 T cells was much denser than non-manipulated CD4^+^ T cells. This is in accordance with studies showing that increased mitochondrial cristae density does not always reflect an overall increased in mitochondrial mass, but increased cristae density correlates with elevated oxygen consumption [90].

Moreover, extracellular flux analysis of mito-transferred CD4^+^ T cells showed enhanced aerobic respiration, which is supported by our proteomic data. Our data on mitochondrial mass and mtDNA copy number hint that mito-transfer perhaps transiently increased mitochondrial mass before or during the cellular reprogramming process. At a minimum, the increased mitochondrial activity can be attributed to the increased expression of proteins related to ETC. This also is likely the reason behind the concurrent rise in glycolysis in mito-transferred CD4^+^ T cells, as the amount of acetyl-CoA needed for the TCA cycle significantly increased. Acetyl-CoA is primarily generated either through β-oxidation of intracellular fatty acids or glycolysis [91]. We did not assess the preferential utilization of lipids in this study, and the medium preparation used to perform the metabolic analysis did not contain a significant amount of lipids. In line with Le Chatelier’s principles of substrate equilibrium, it can be argued that after mito-transfer, CD4^+^ T cells from old mice upregulated glycolysis to compensate for lagging intracellular levels of acetyl-CoA. Additionally, the ATP production rate of CD4^+^ T cells isolated from old mice rose by 40% in comparison to non-manipulated CD4^+^ T cells. Primary information regarding T cell metabolism in the context of aging is still sparse [92]. As of now, it seems the impact of mito-transfer on the metabolism of CD4^+^ T cells differs from the Warburg phenomenon, as there was no bias towards glycolysis, and mito-transfer decreased mitoROS in the CD4^+^ T cells from old mice. Moreover, these data collectively suggest that the mitochondrial cristae of CD4^+^ T cells restructured after mito-transfer, favoring increased cristae density per mitochondria rather than an increase in mitochondrial mass.

Increased cristae density would also explain why the level of mitoROS was significantly higher than that of non-manipulated CD4^+^ T cells after mito-transferred CD4^+^ from old mice were activated with PMA/IONO. We also noted that the switch from Ox-Phos to aerobic glycolysis in mito-transferred CD4^+^ T cells was much faster than that of non-manipulated CD4^+^ T cells from old mice. Beyond the observed impact on ROS and metabolism, there were other improved T cell functions. Indeed, we identified significant improvements in TCR-signaling kinetics, expression of a T cell activation marker (CD25), cytokine production, and an increase in activation-induced proliferation. Compared to non-manipulated CD4^+^ T cells, the proportion of activated T cells expressing CD25^+^ and CD69^+^ doubled after mito-transfer and 24 h of activating stimulus. We also noted a concomitant doubling of CD4^+^ T cells producing IL-2, which correlated with increased cell proliferation and contributed to the alteration of cytokine profiles.

The IL-2 signaling pathway is immunomodulatory [93-95], and while CD4^+^ T cells, CD8^+^ T cells, NK cells, NK T cells, and dendritic cells are all capable of secreting IL-2, activated CD4^+^ T cells are its largest producer [85, 93, 96]. Previous research has shown that CD4^+^ T cells from elderly humans and mice produce insufficient amounts of IL-2, as well as insufficiently express the high-affinity IL-2 receptor IL-2Rα (CD25) necessary for signal transduction [97, 98], and the current consensus is that aging-associated defects in IL-2 signaling can dynamically shift the architecture of adaptive immune responses or even promote T cell anergy [5, 97, 99]. The underlying mechanics regarding blunted IL-2 production and related signaling in the elderly are still unclear [5, 99, 100]. Intracellular IL-2 signaling transduction occurs through the Janus kinase/signal transducer and activation of transcription factor (JAK/STAT), phosphoinositide-3-kinase (PI3K), and mitogen-activated kinase (MAPK) pathways, all of which are sensitive to oxidative modifications [101, 102]. Previous studies have also characterized the role of redox equilibrium in maintaining proper lymphocyte function, and mitoROS, specifically H_2_O_2_, modulates IL-2 expression in T cells [15, 103]. Supplementation with the antioxidant vitamin E improved the ability of T cells from the elderly to produce IL-2 and proliferate [104]. However, maintaining the intracellular redox balance is delicate [23, 37, 38], and excess antioxidants can hinder proper T cell activation and cytokine production as others have described [105, 106].

Our *in vivo* IAV and *M.tb* infection outcomes highlight the potential of mito-transfer as an immune-booster in the elderly. We demonstrated that mito-transfer can enhance control of viral and non-viral pathogen infections *in vivo,* and the enhanced protection is likely related to the preferential generation of antigen-specific effector memory CD4^+^ T cells. As little as 2.0 × 10^6^ naïve mito-transferred CD4^+^ T cells from old mice were capable of differentiating into either IAV or *M.tb* specific effector CD4^+^ T cells that had a physiological impact on IAV-induced morbidity and mortality and on *M.tb* burden. This study could not fully delineate whether antigen-specific effector CD4^+^ T cell quality (antigen-specific) and or quantity (proliferative capacity) was preferentially enhanced by mito-transfer. Based on our collective data, we hypothesize that both parameters (i.e. antigen specificity and proliferative capacity) might improve *in vivo*, and consider that improved disease outcomes should be achievable with an increase in the quantity of mito-transferred naïve T cells injected into mice.

The capacity to upscale the process of mito-transfer from *in vitro/ex vivo* towards *in vivo* models is likely limited by available techniques to perform MRT and the number of viable mitochondria or mtDNA needed to revert a specific mitochondrial metabolic abnormality in an organism [107, 108]. Mitochondrial encapsulation or injections can facilitate *in vivo* studies [109, 110] and Chang *et al.* showed that cell-penetrating peptide (pep-1) mediated delivery of healthy mitochondria to MCF-7 breast cancer cells reversed Warburg metabolism, decreased oxidative stress, and increased cancer cell susceptibility to chemotherapeutics [109].

Nevertheless, mito-transfer is an especially promising avenue for the immunological landscape, partly because it is plausibly easy to isolate and re-insert autologous and allogeneic lymphocytes/leukocytes in a clinic or lab setting. This could have transformative impact on aging-associated immune dysregulations and on immune based dysregulation that generally involve mitochondrial dysfunction. For example, work by Vaena *et al*. showed that anti-tumor T cells from aged individuals succumb to ceramide accumulation-induced mitochondrial dysfunction and this consequently inhibits anti-tumor T cell responses [111]. It would be interesting to assess whether mito-transferred anti-tumor CD8^+^ T cells from aged mice and elderly can mitigate ceramide induce mito-dysfunction and promote anti-tumor activity. Guo *et al*. and team also demonstrated that mitochondrial reprograming through upregulation of mitochondrial pyruvate carrier was essential to restoring the functionality of exhausted CD8^+^ T cells, and this improved their anti-tumor capacities [112]. Undoubtedly, future studies will need to investigate the full potential of mito-transfer on anti-tumor immunity.

In conclusion, under our experimental conditions, mito-transfer decreased mitochondrial dysfunction, improved activation, cytokine production, and proliferation of CD4^+^ T cells from old mice. Importantly, mito-transfer had a physiological impact on IAV-induced morbidity and mortality, and on *M.tb* burden *in vivo*. In addition to mitoROS mediated redox signaling [113-117], calcium mobilization and buffering in coordination with the endoplasmic reticulum (ER) [13, 23, 45, 118], cytochrome c mediated apoptosis [119-123], NAD^+^ mediated epigenetic regulation (sirtuins) [124, 125], as well as protein transport and degradation [126, 127], are all interdependent on proper mitochondrial form and function. We suspect that the impact of mito-transfer on other aspects of T cell function were also improved. More in-depth mechanistic studies outside the scope of this current study are warranted to elucidate the specific means by which mito-transfer altered the pathways in CD4^+^ T cells in old mice and the elderly.

## Acknowledgments

We would like to acknowledge the thoughtful discussions with Drs. Susan Weintraub, Larry S. Schlesinger, Douglas Green & James Stambulli that have strengthened this manuscript. We would also like to acknowledge Drs. Gourav Choudhury and Marcel Daadi for sharing their MEF isolation protocol. Finally we would like to acknowledge Ms. Kimberley Olsen for her technical and administrative assistance throughout the duration of this project.

## Author Contributions

This project was conceived by CAH & JT; CAH designed the experiments; CAH, AM, AV, SG, VD, AW, AA, JD, AH, RA, KC, HZ, and HC performed experiments; CAH and JG performed the statistical analysis, CAH wrote the manuscript; CAH, PS, LMS, YW, JBT and JT edited the manuscript.

## Data Availability

The authors declare that the data supporting the findings of this study are available within the paper and its supplementary information files. Reagents are available upon request.

## Supplementary figures

**Supp. Fig. 1.**
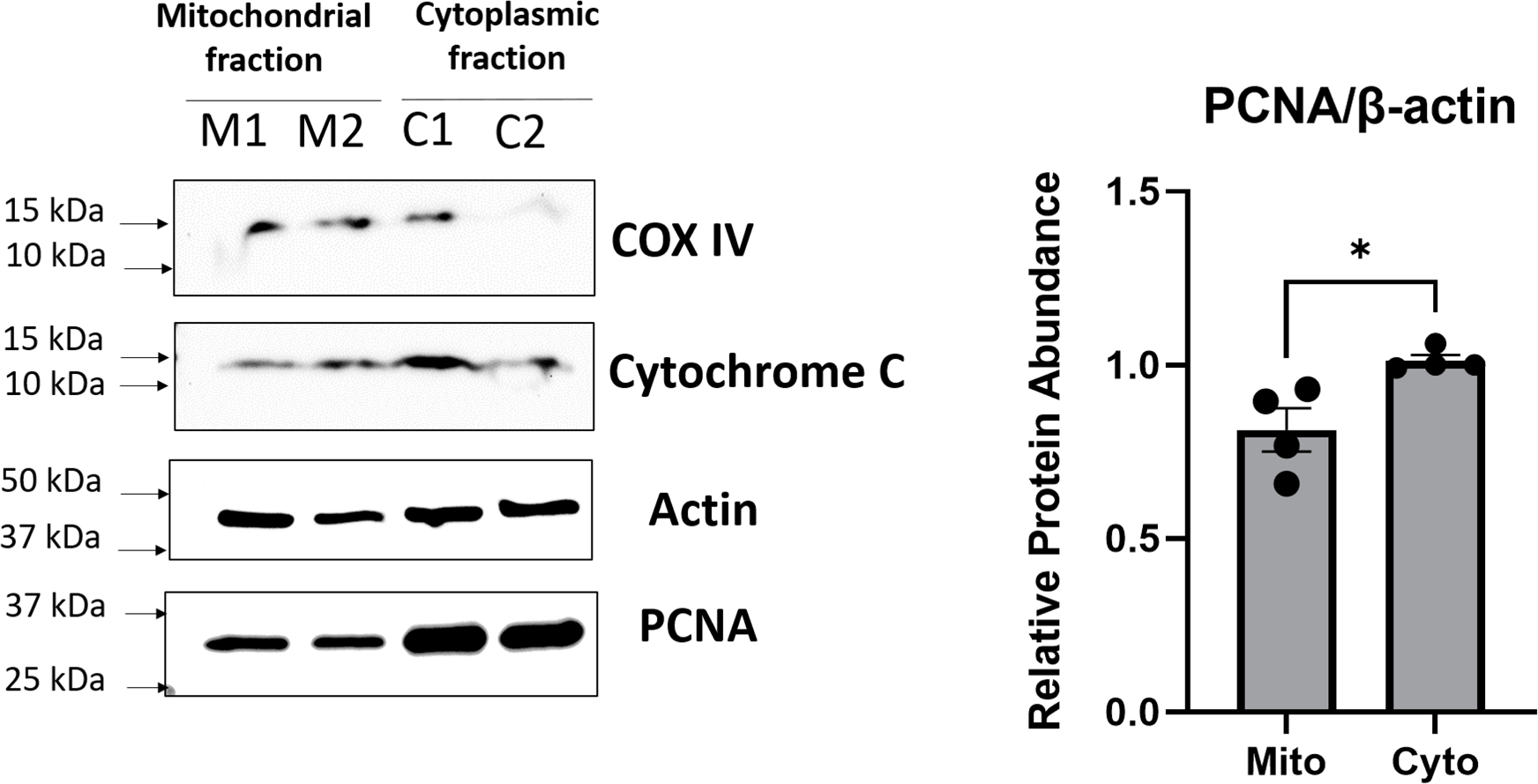
Purity of mitochondrial isolation. **A)** Representative wester blot images of probed mitochondrial and nuclear proteins in the mitochondrial fraction and cell fragments/unlysed cells after mitochondrial isolation. **B)** Relative amount of PCNA normalized to B-actin. (n=4), p<0.05=significant (*) using unpaired Student’s t-test.

**Supp. Fig. 2.**
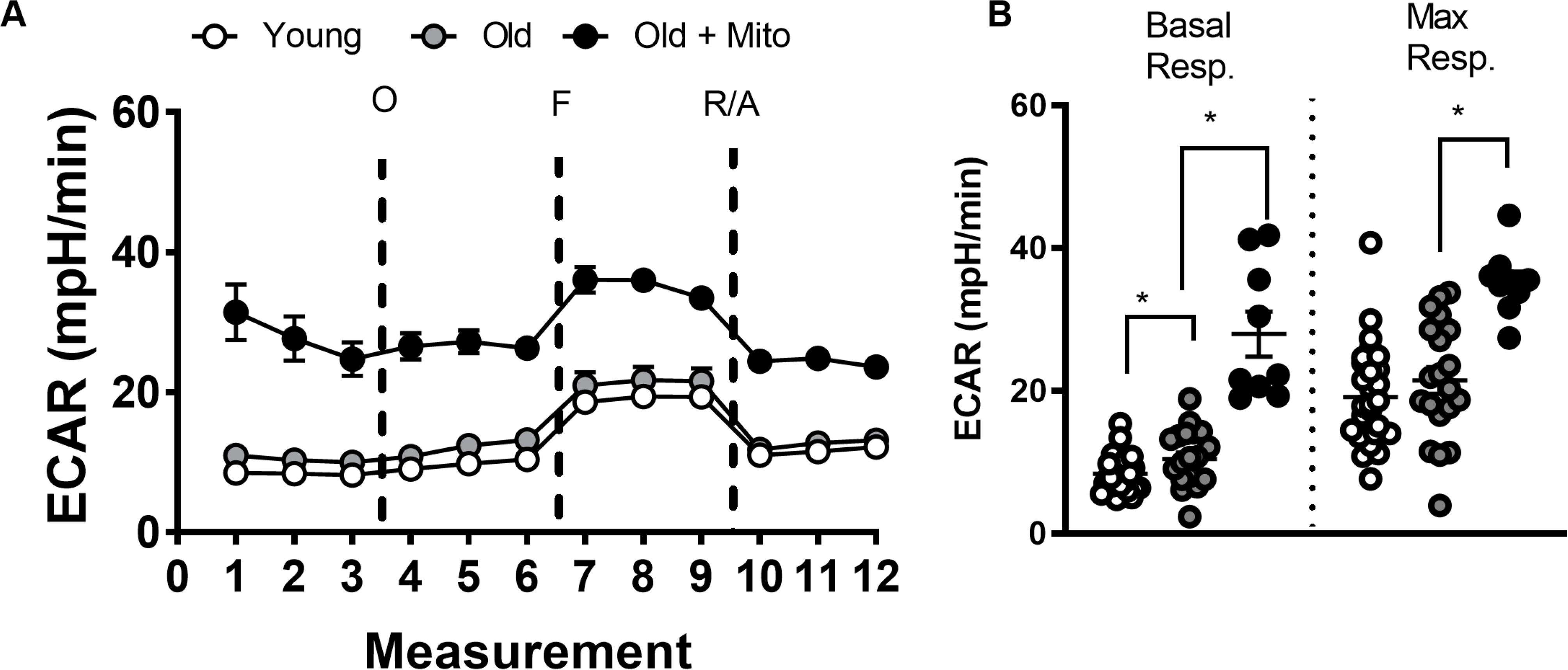
Mito-transfer increases ECAR of CD4^+^ T cells from old mice. **A)** Kinetic curve of ECAR collected from mito-stress test assay, **B)** ECAR corresponding to basal and maximal respiration of CD4^+^ T cells. 3-4 mice were used per group and experiments were replicated twice (n=3), p<0.05=significant (*) using unpaired Student’s t-test.

**Supp. Fig. 3.**
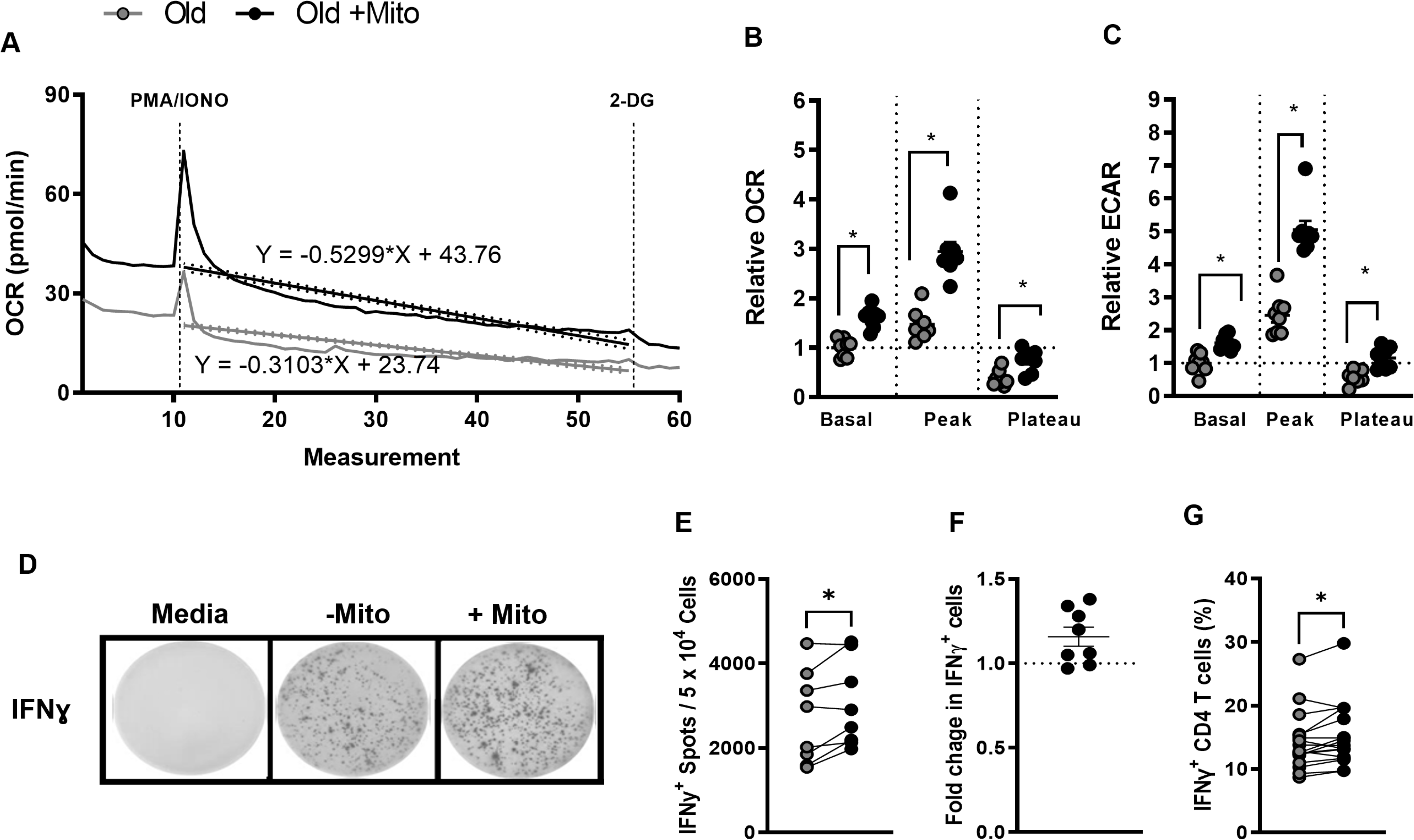
Mito-transfer enhaced T cell activation induced metabolic switch and increased IFNγ^+^ CD4^+^ T cells. **A)** The slope of OCR depression after acute injection of PMA/IONO in mito-transferred and non-manipulated CD4+ T cells from old mice. The relative change in **B)** OCR and **C)** ECAR of activated T cells (non-normalized to group). **D-G)** CD4^+^ T cells from old mice with or without mito-transfer were stimulated with PMA/Ionomycin, after which various assays were used to quantify IFNγ production. **D)** Representative EliSpot images, **E)** the number IFNγ^+^ producing T cells, and **F)** the fold change in IFNγ^+^ producing CD4^+^ T cells from old mice with or without mito-transfer, after stimulation with PMA/Ionomycin (24hr). The **G)** percent of IFNɣ^+^ CD4^+^ T cells via intracellular cytokine staining, in CD4^+^ T cells from old mice with or without mito-transfer, after stimulation with PMA/Ionomycin (4h). 3-4 mice per group, repeated at least once (n ≥ 2). p< 0.05 = significant (*) using paired Student’s *t*-test.

**Supp. Fig. 4.**
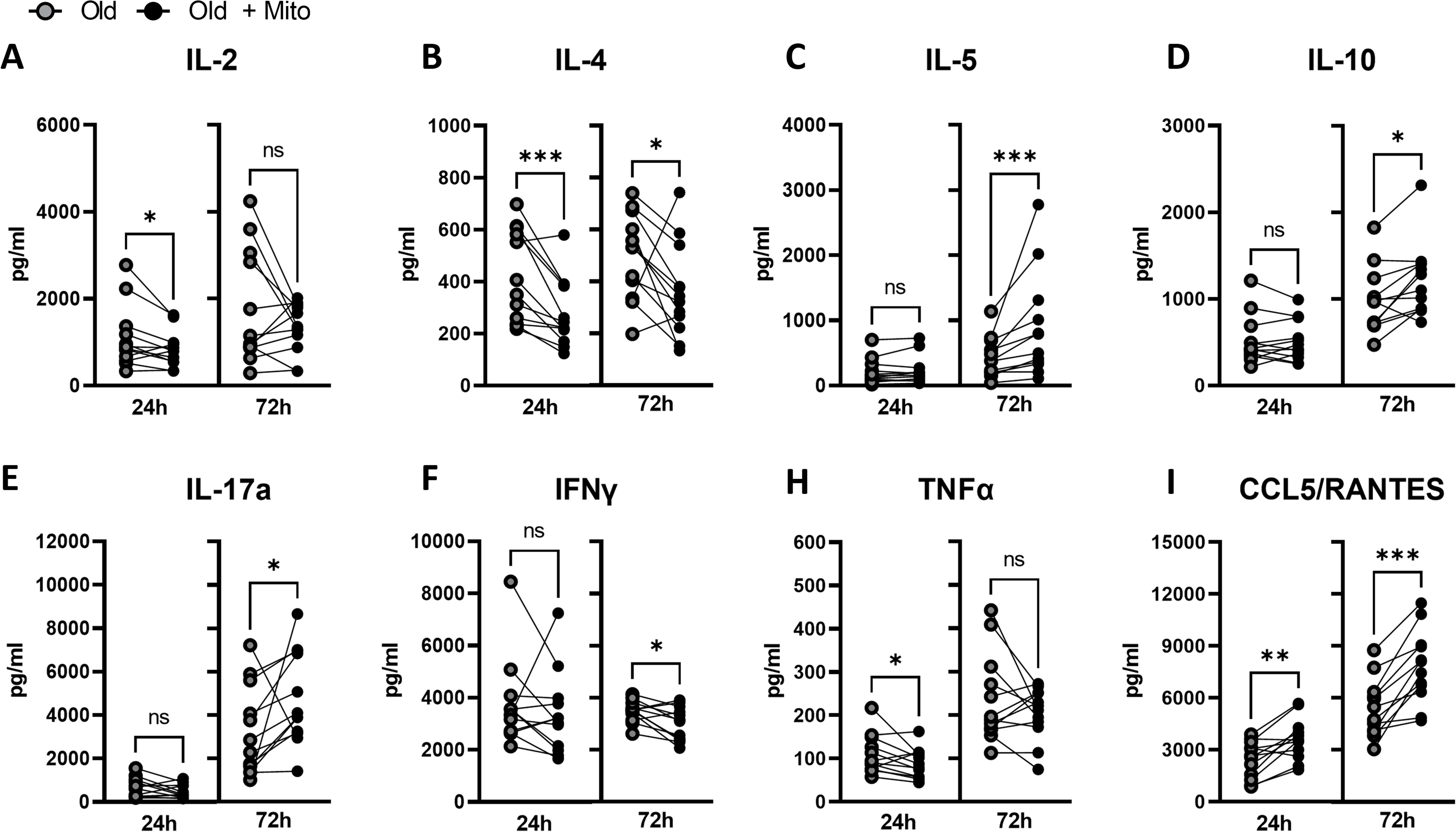
Mito-transfer alters T cell activation & cytokine production of CD4^+^ T cells in old mice. CD4^+^ T cells from old mice with or without mito-transfer were stimulated with PMA/Ionomycin. After 24 and 72 h of stimulation with PMA/Ionomycin, the supernatants were examined by Luminex array for cytokines produced. p ≤ 0.05 = *, p ≤ 0.01 = **, or p ≤ 0.001 = *** using paired Student’s *t*-test.

**Supp. Fig. 5.**
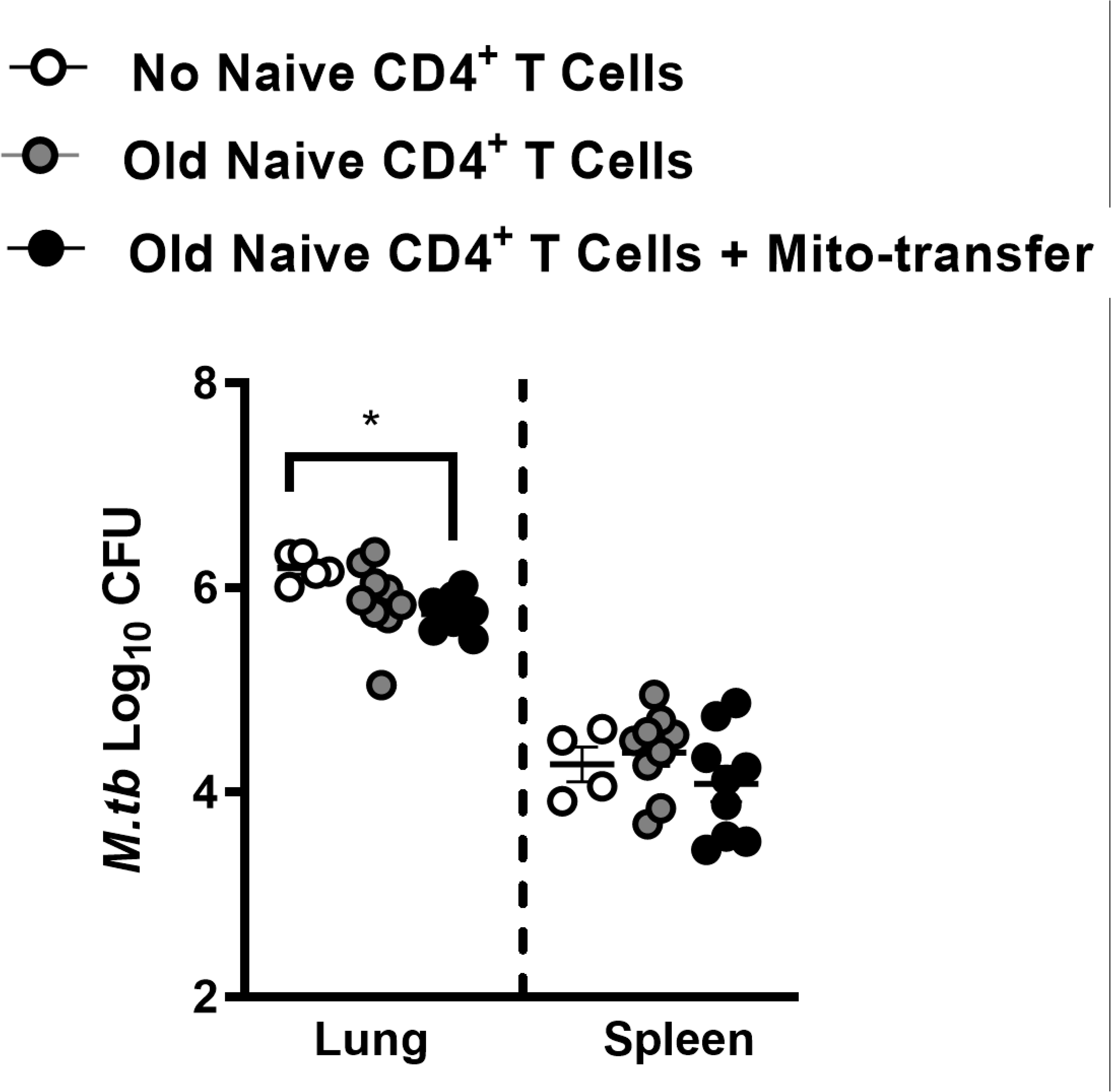
Mito-transfer in naïve CD4 T cells from old mice protected mice against pathogens. Naïve CD4^+^ T cells from old mice with or without mito-transfer, or PBS were tail vein injected in Rag1-KO mice that were subsequently infected with *M.tb*. Lung and spleen CFU burden after 21 days of *M.tb* infection in Rag1-KO mice adoptively treated with 1x 10^6^ naïve CD4^+^ T cells from old mice with or without mito-transfer. 4-8 mice per group, with p ≤ 0.05 = *, using unpaired Student’s *t*-test.

**Supp. Table 1.**
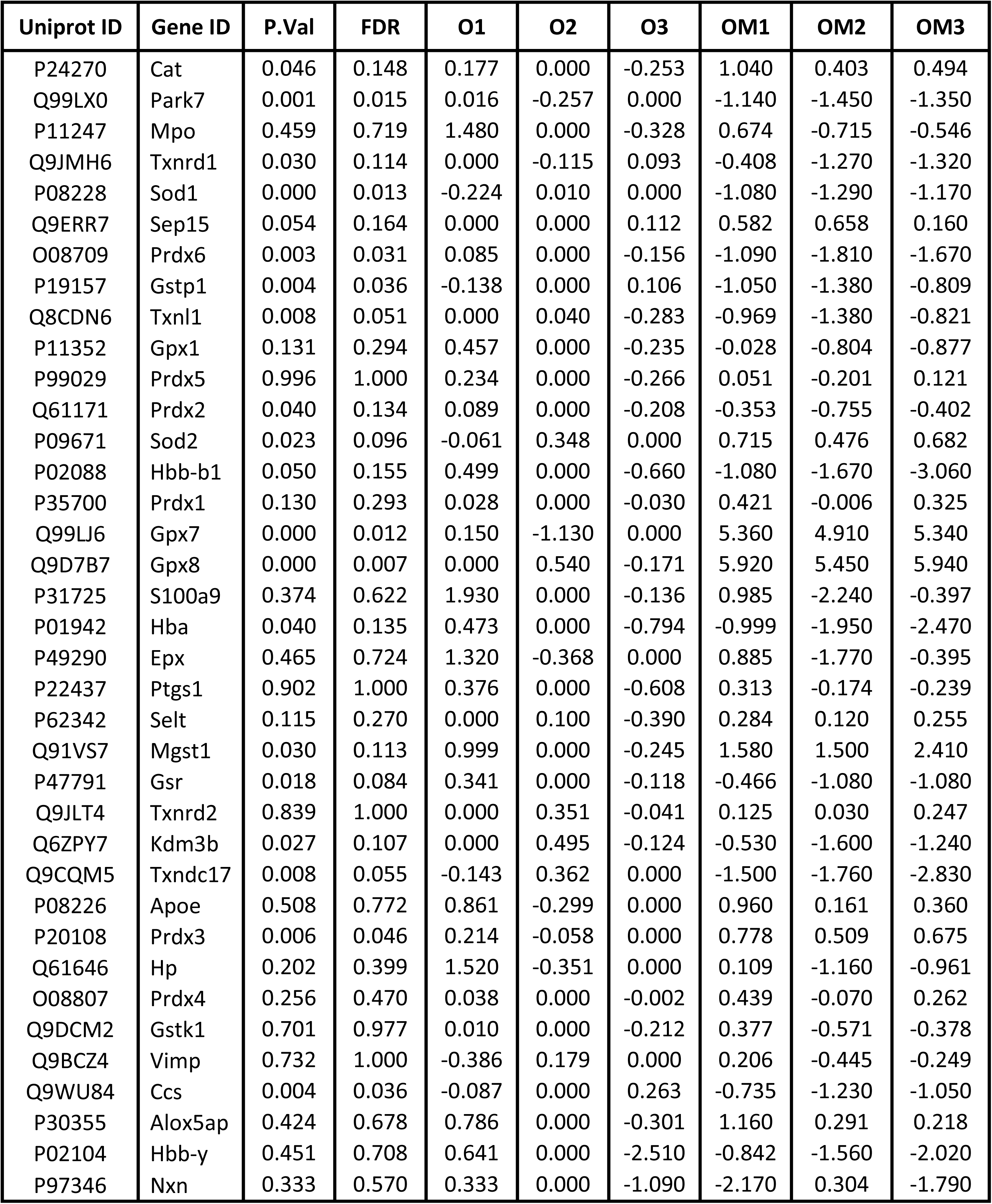
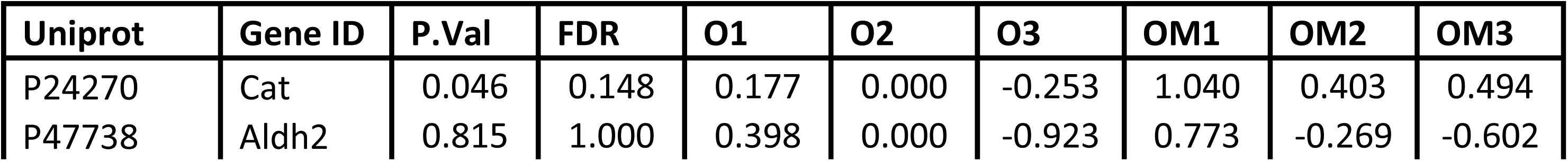
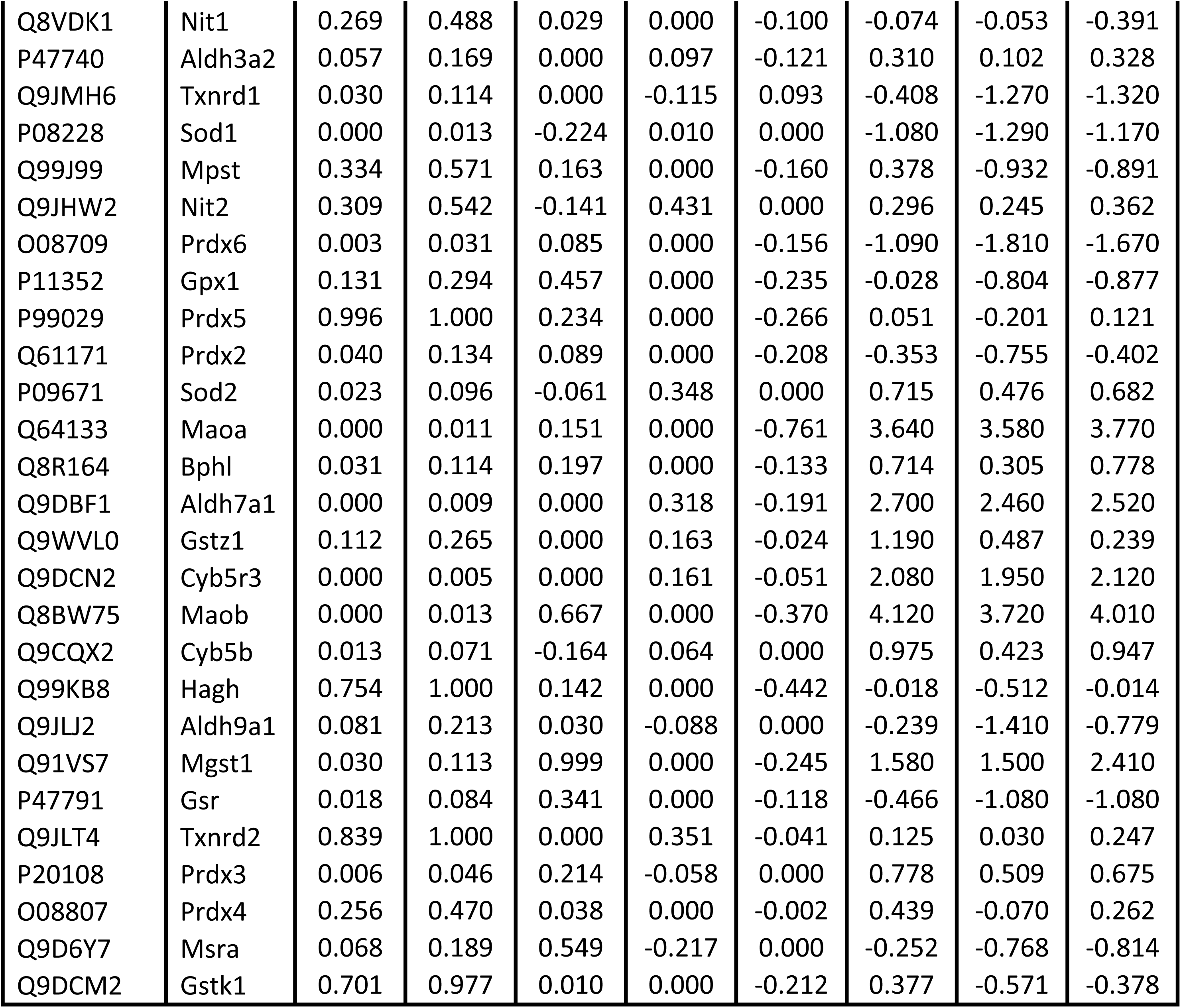
Antioxidant & mitochondrial detoxification proteins were detected in CD4^+^ T cells isolated from old mice after mito-transfer and in non-manipulated CD4^+^ T cells from old mice. CD4^+^ T cells from young and old mice, and from old mice after mito-transfer were cultured for 4 h before processing for mass spectrometry analysis. Data expressed as median protein Log2 fold change of CD4^+^ T cells from old mice, from 3 individual old mice (paired experiment)

**Supp. Table 2.**
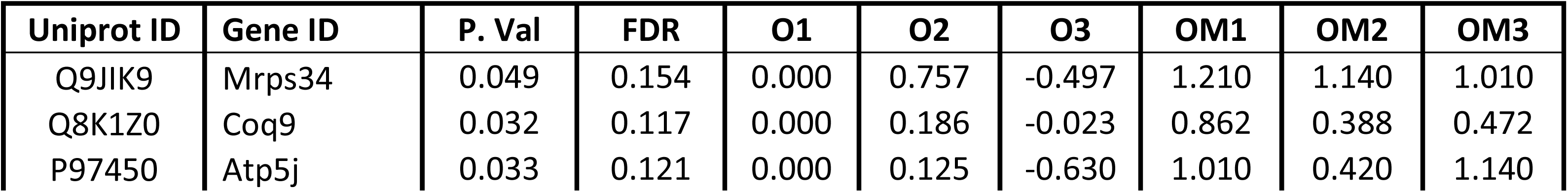
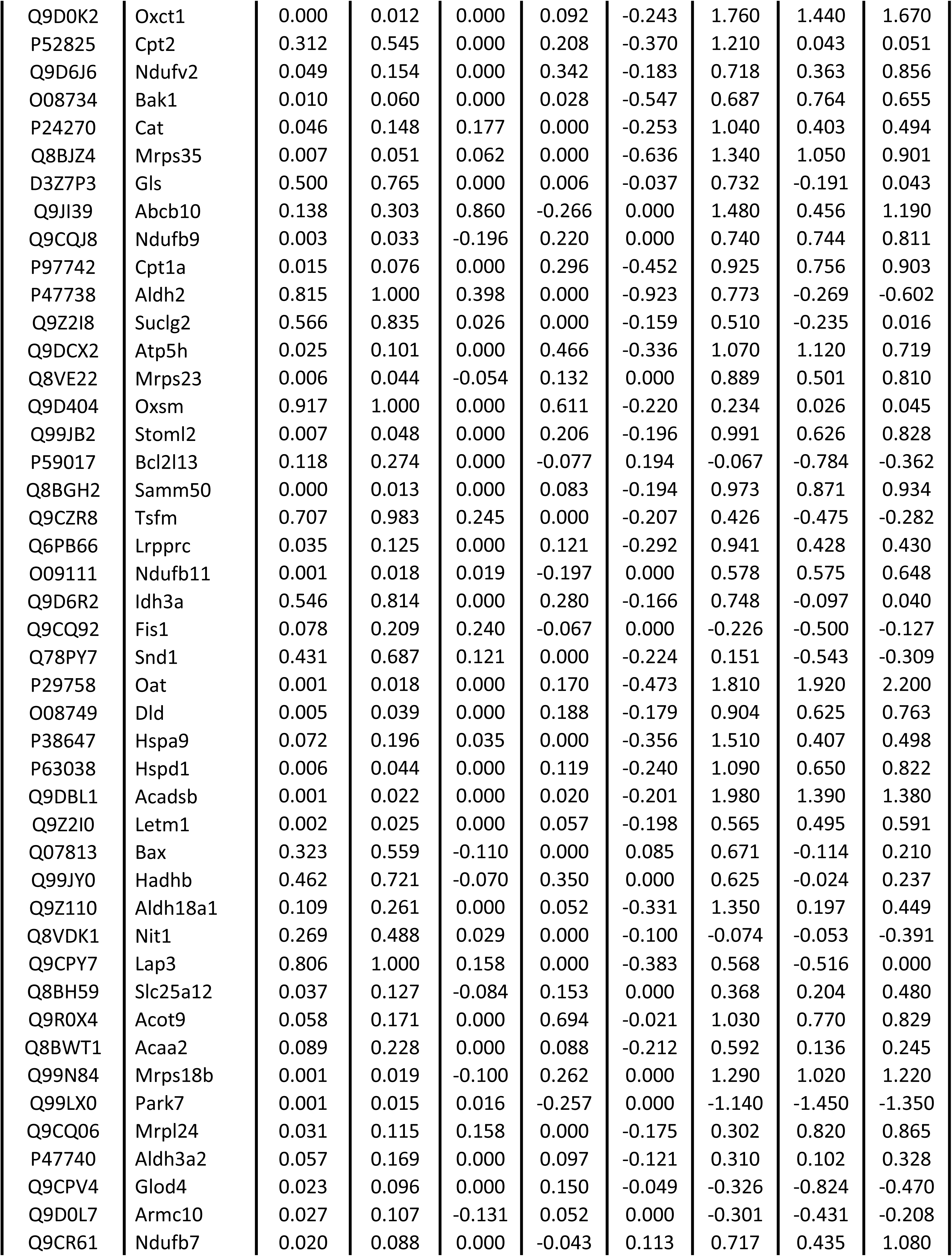
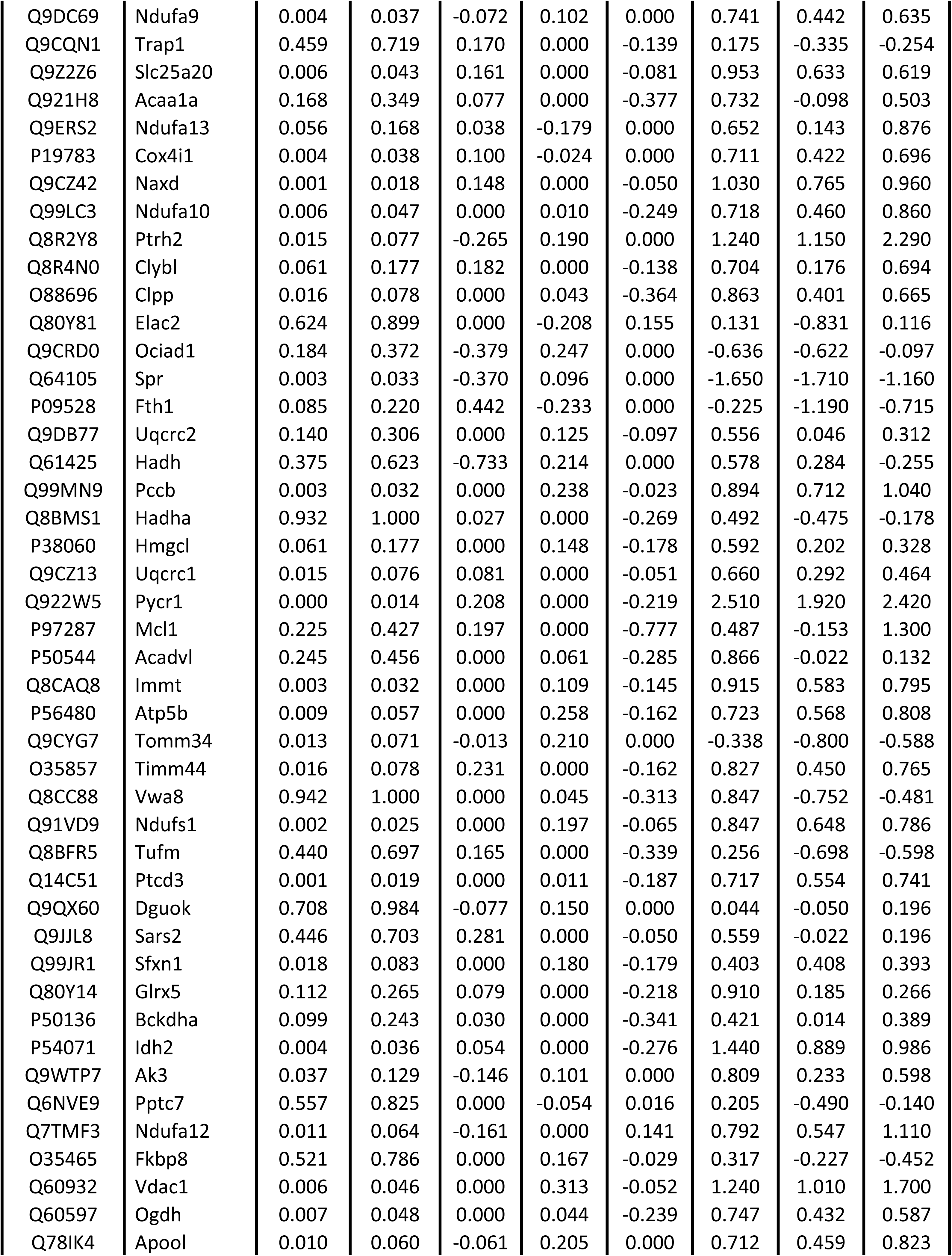
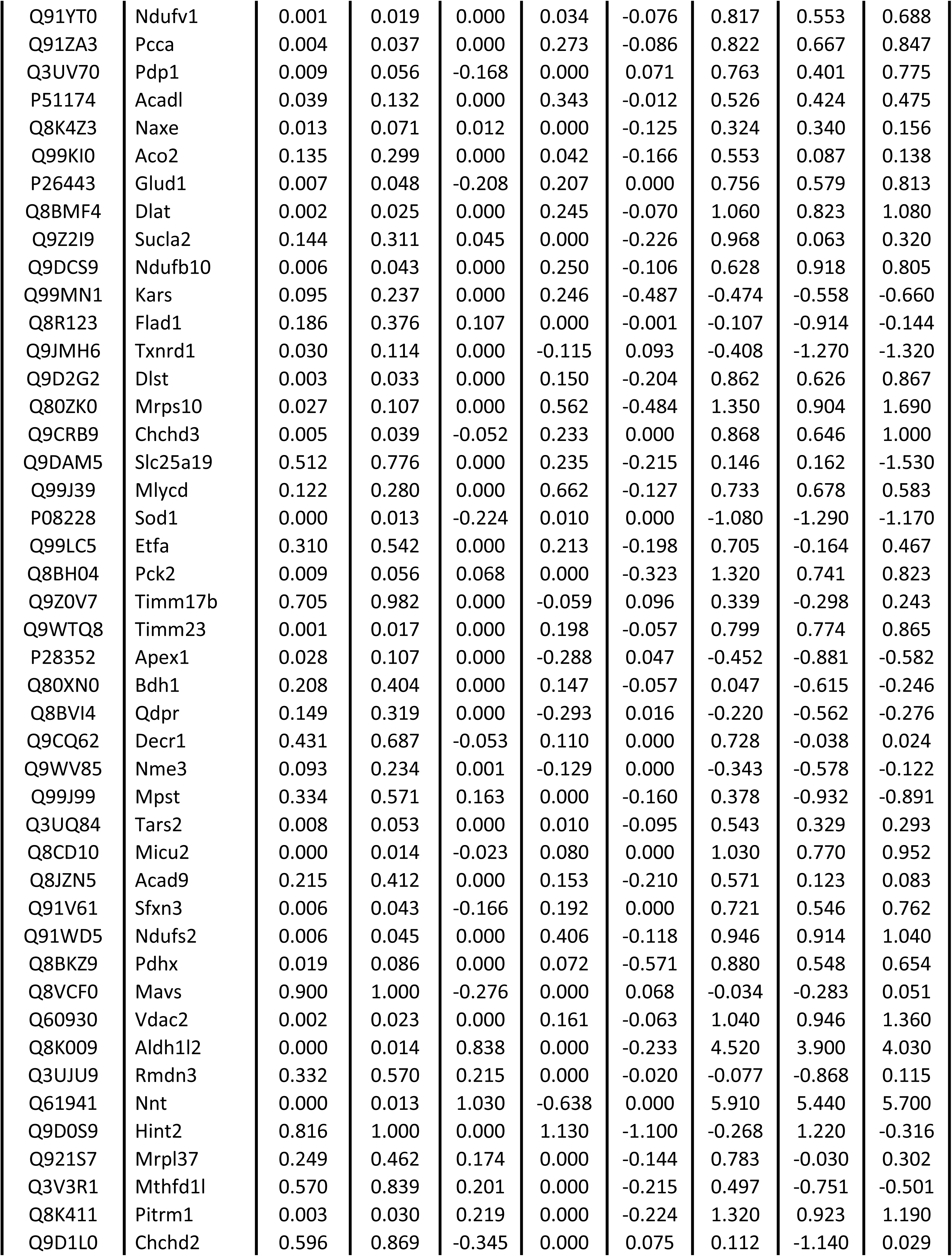
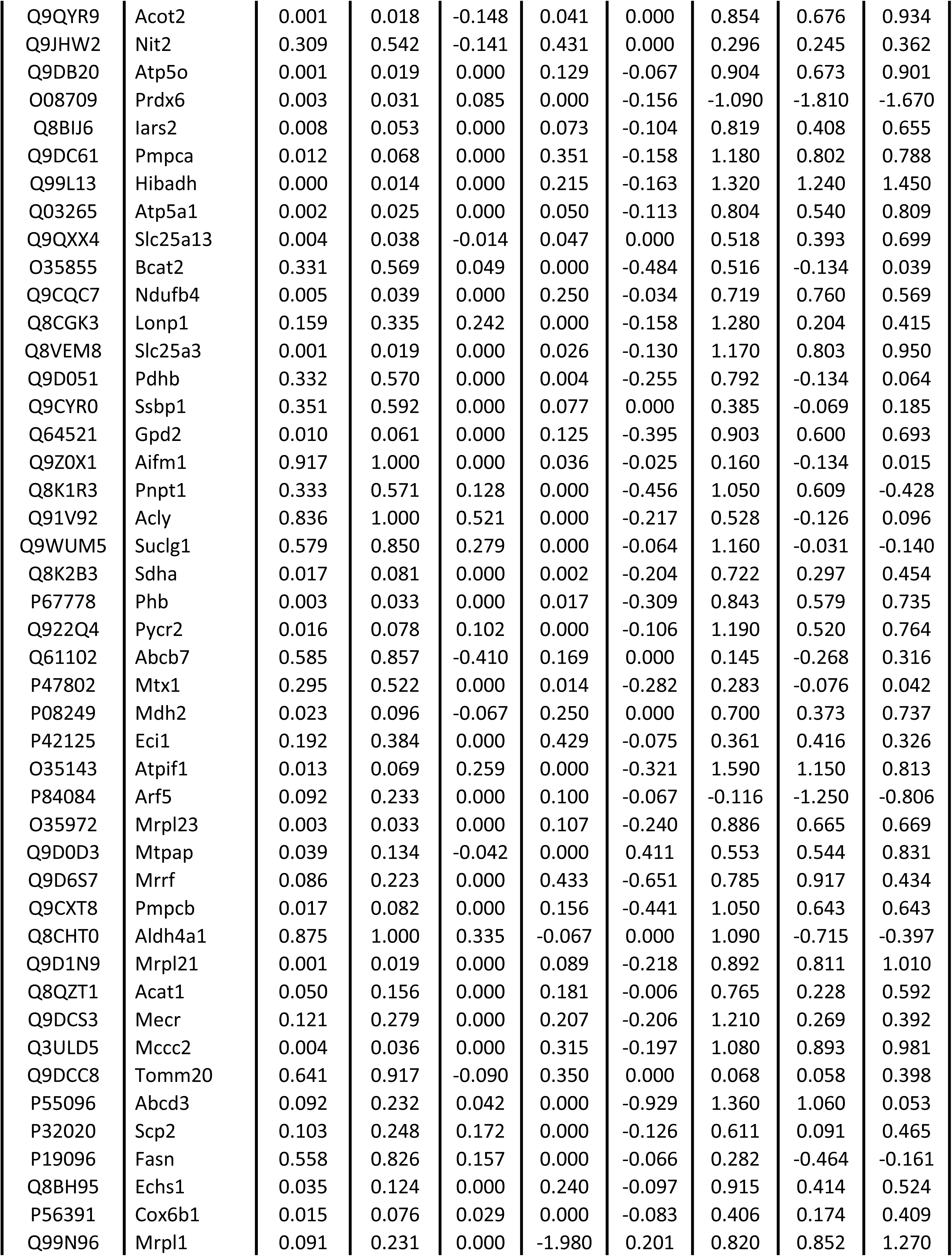
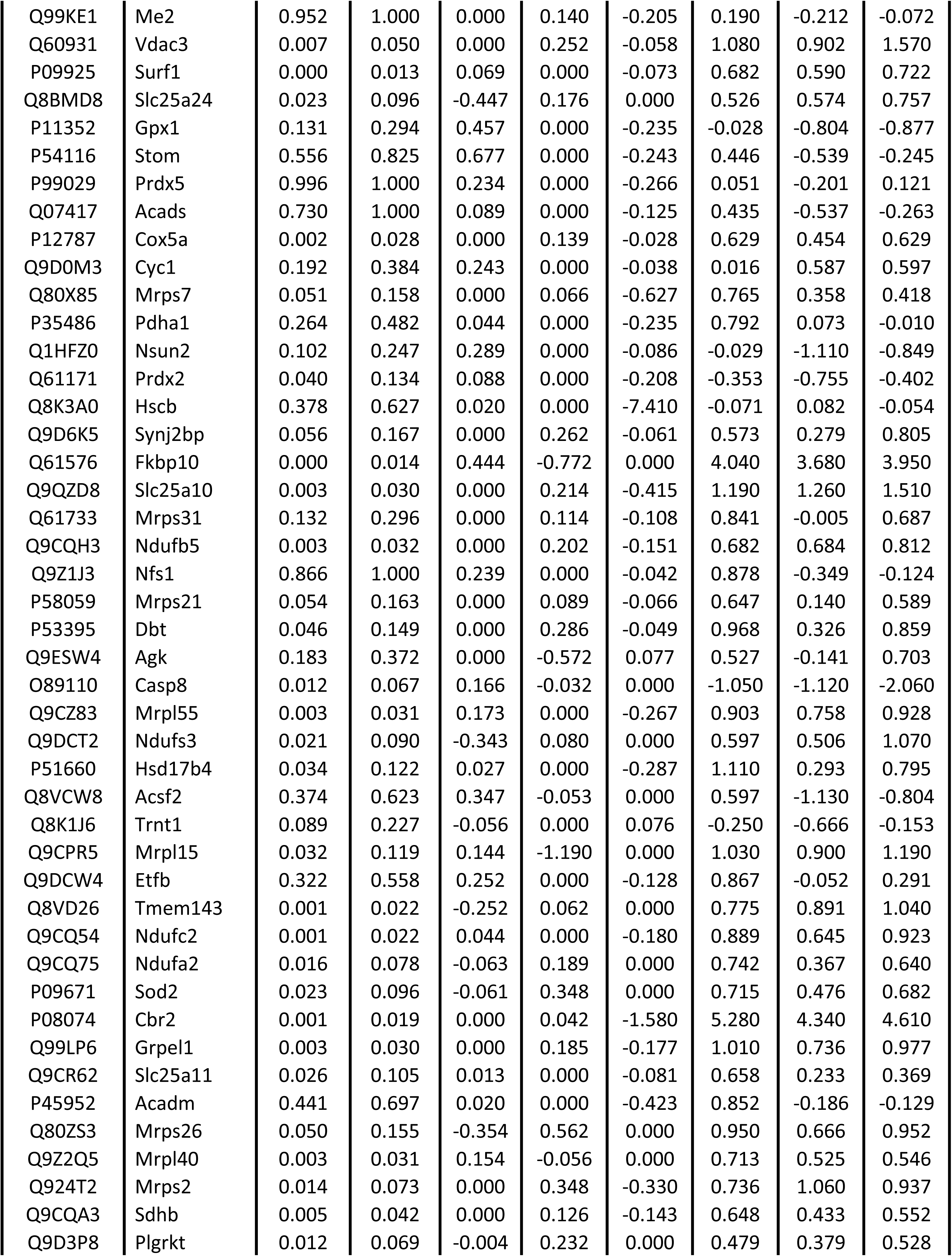
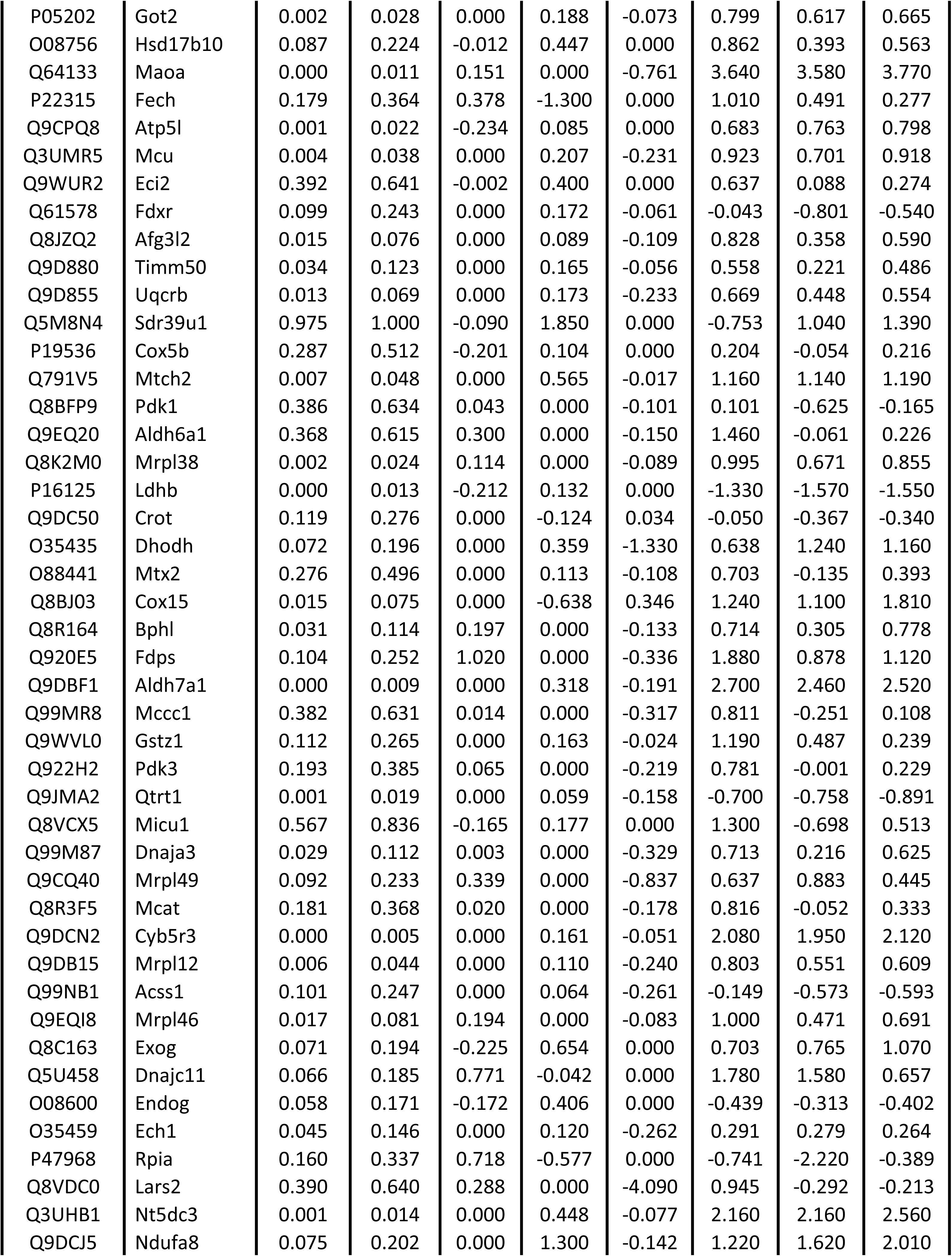
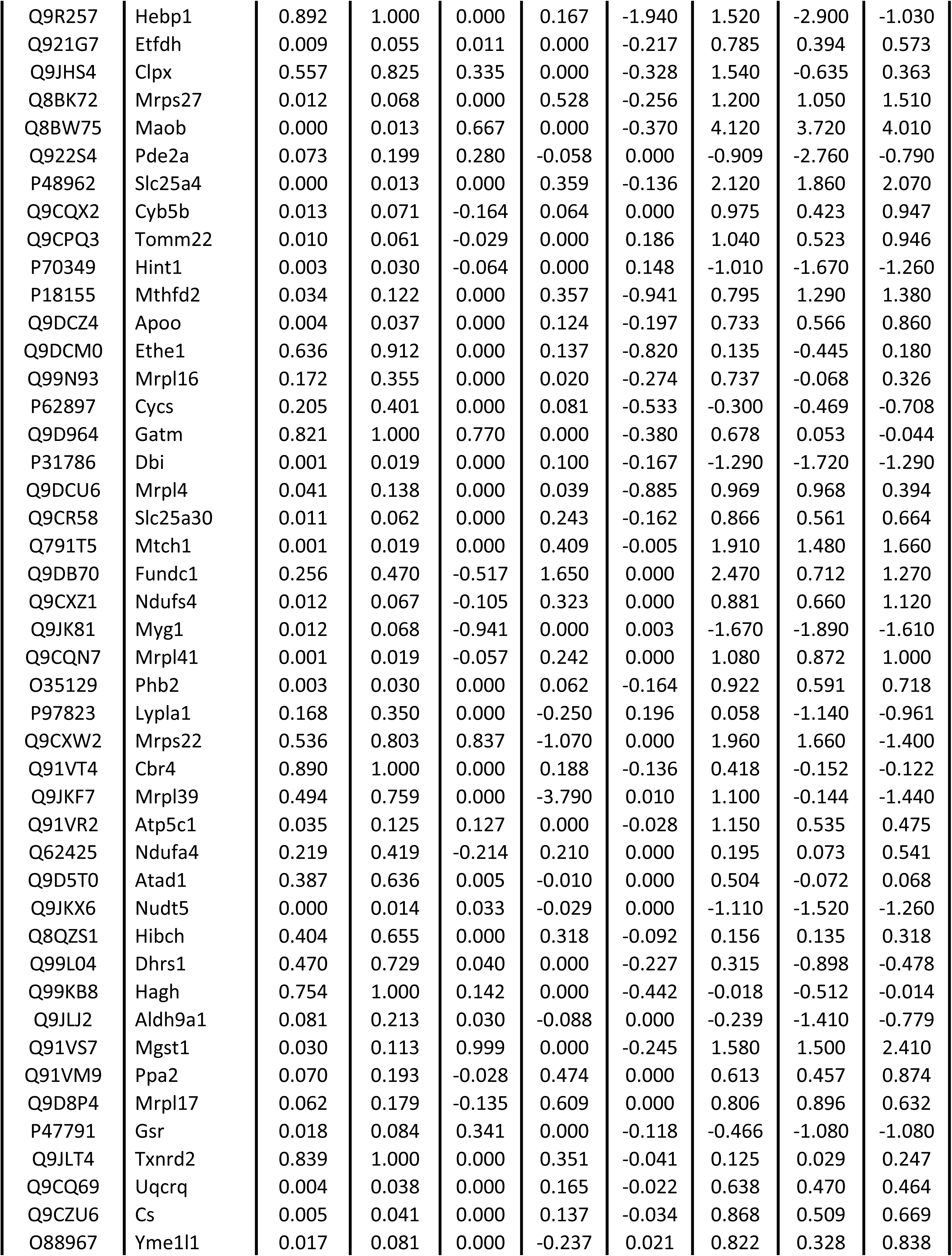
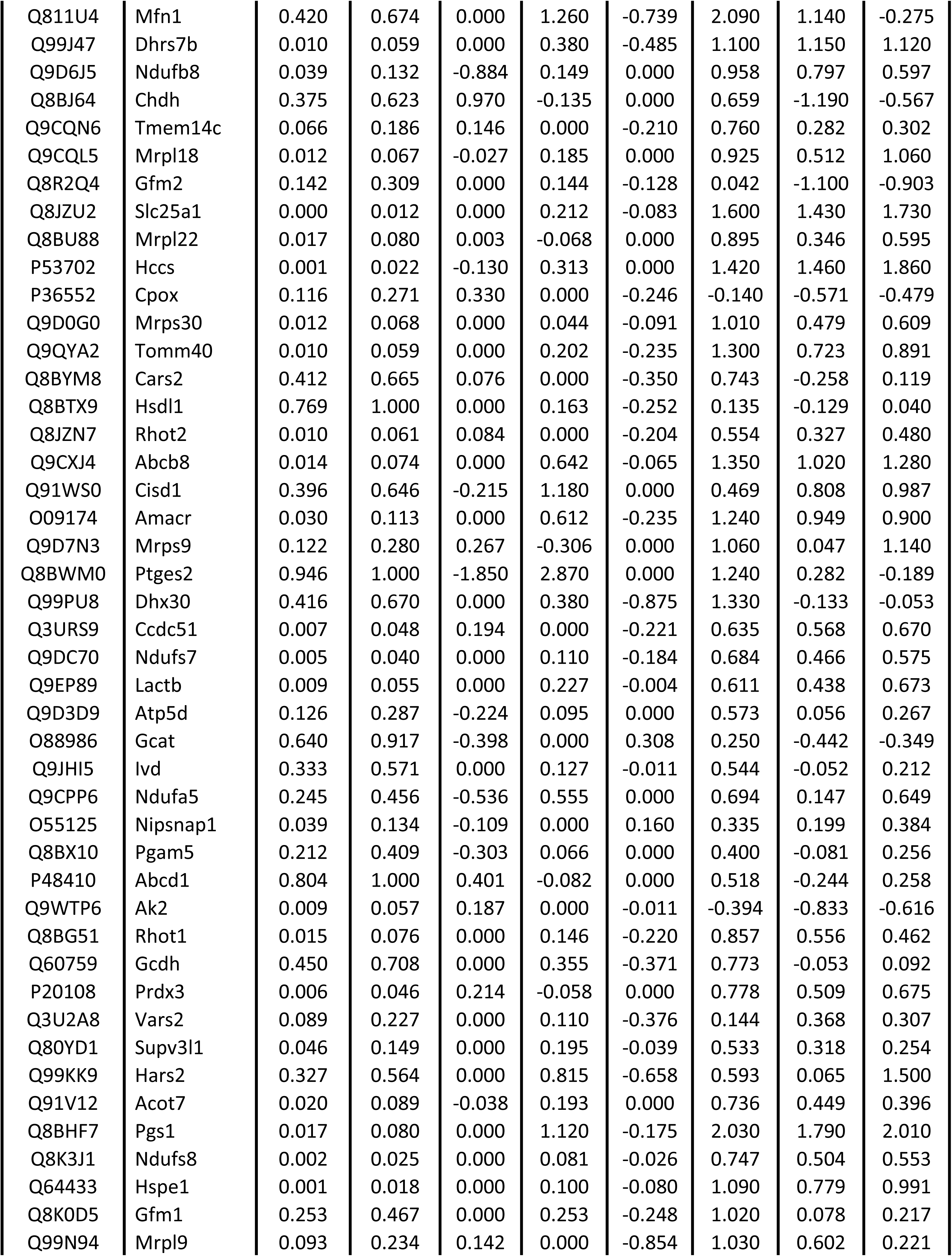
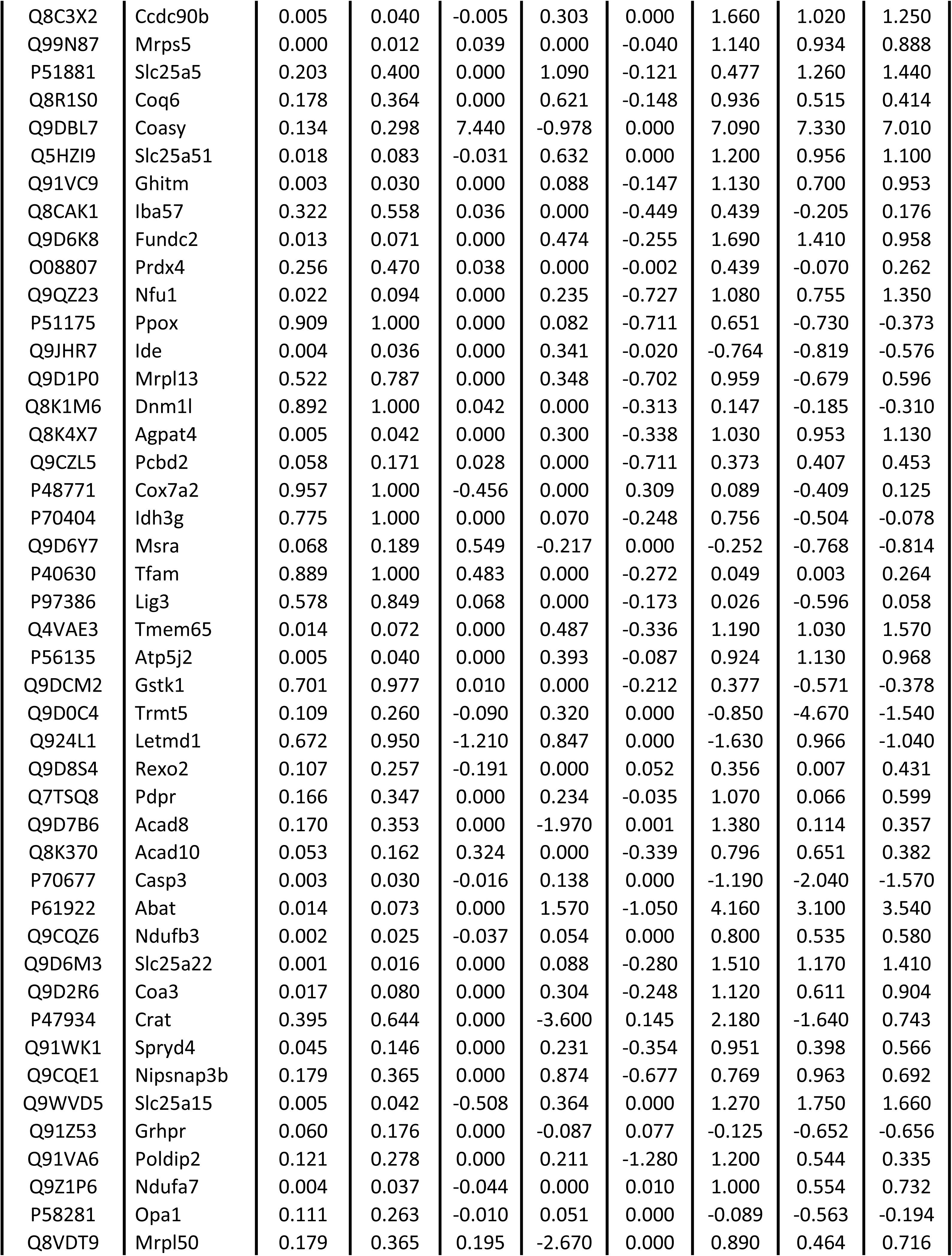
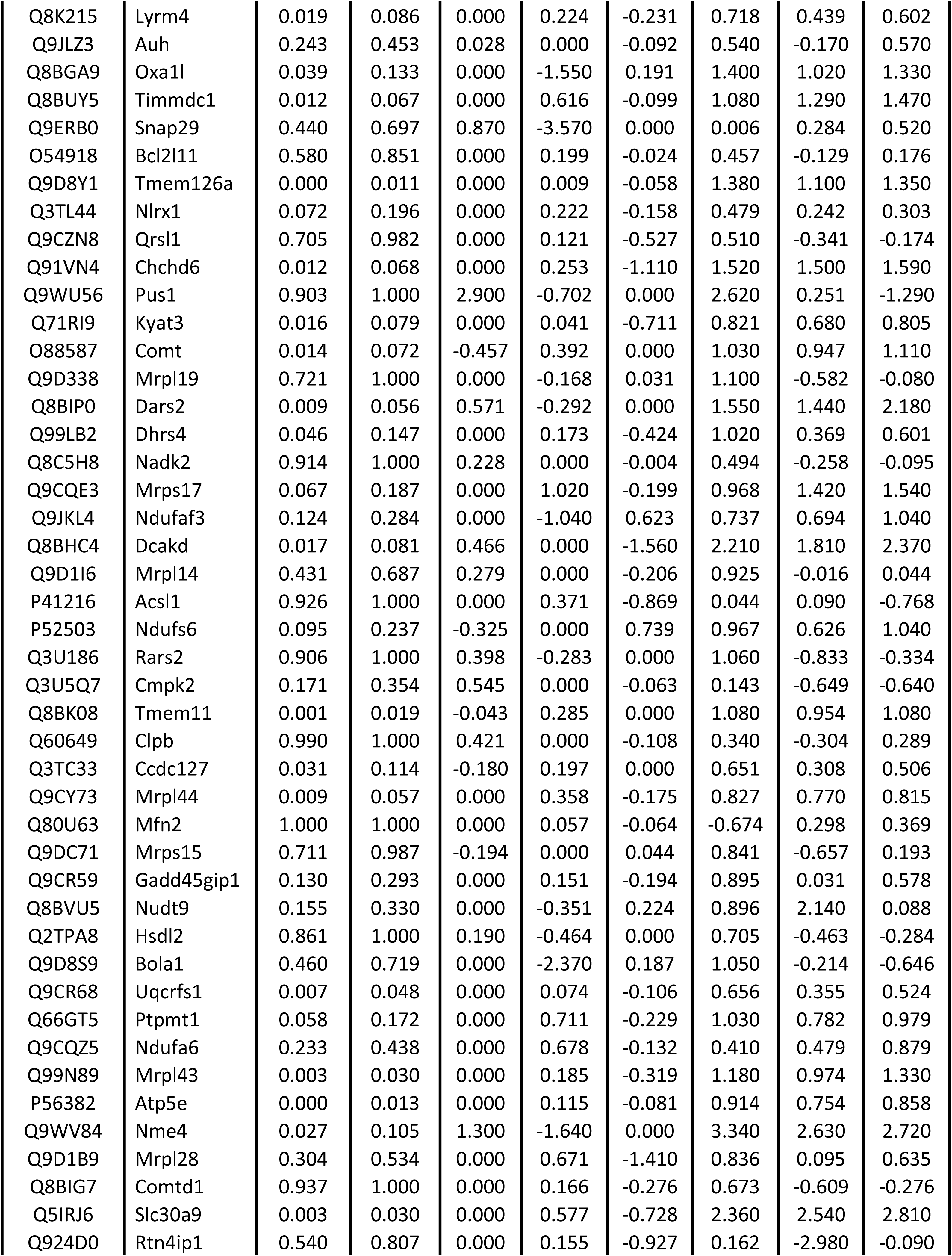
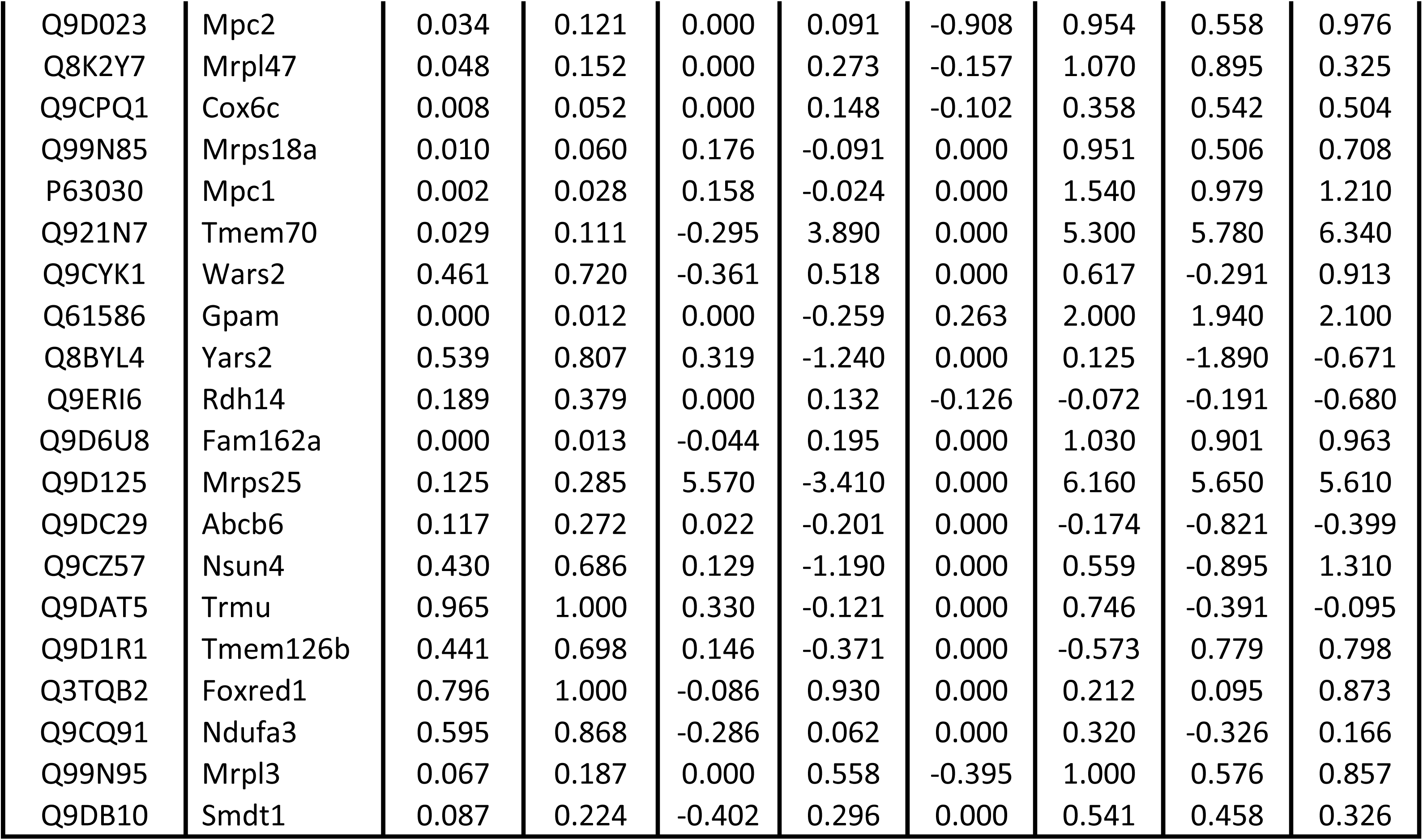
Mitochondrial proteome was detected in CD4^+^ T cells isolated from old mice after mito-transfer and in non-manipulated CD4^+^ T cells from old mice. CD4^+^ T cells from young and old mice, and from old mice after mito-transfer were cultured for 4 h before processing for mass spectrometry analysis. Data expressed as median protein Log2 fold change of CD4^+^ T cells from old mice, from 3 individual old mice (paired experiment)

**Supp. Table 3.**
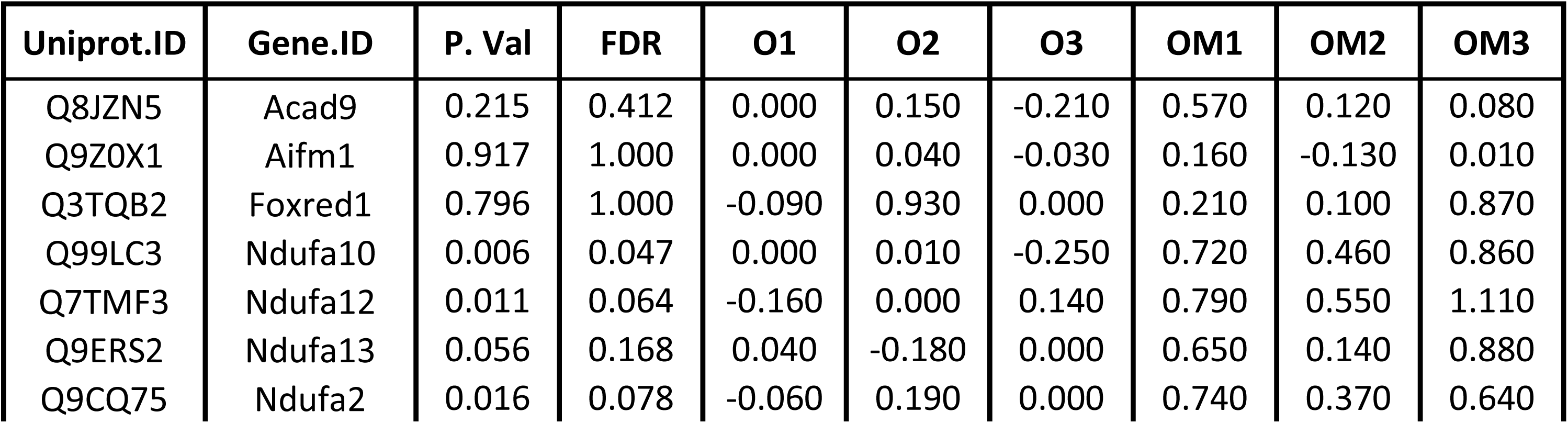
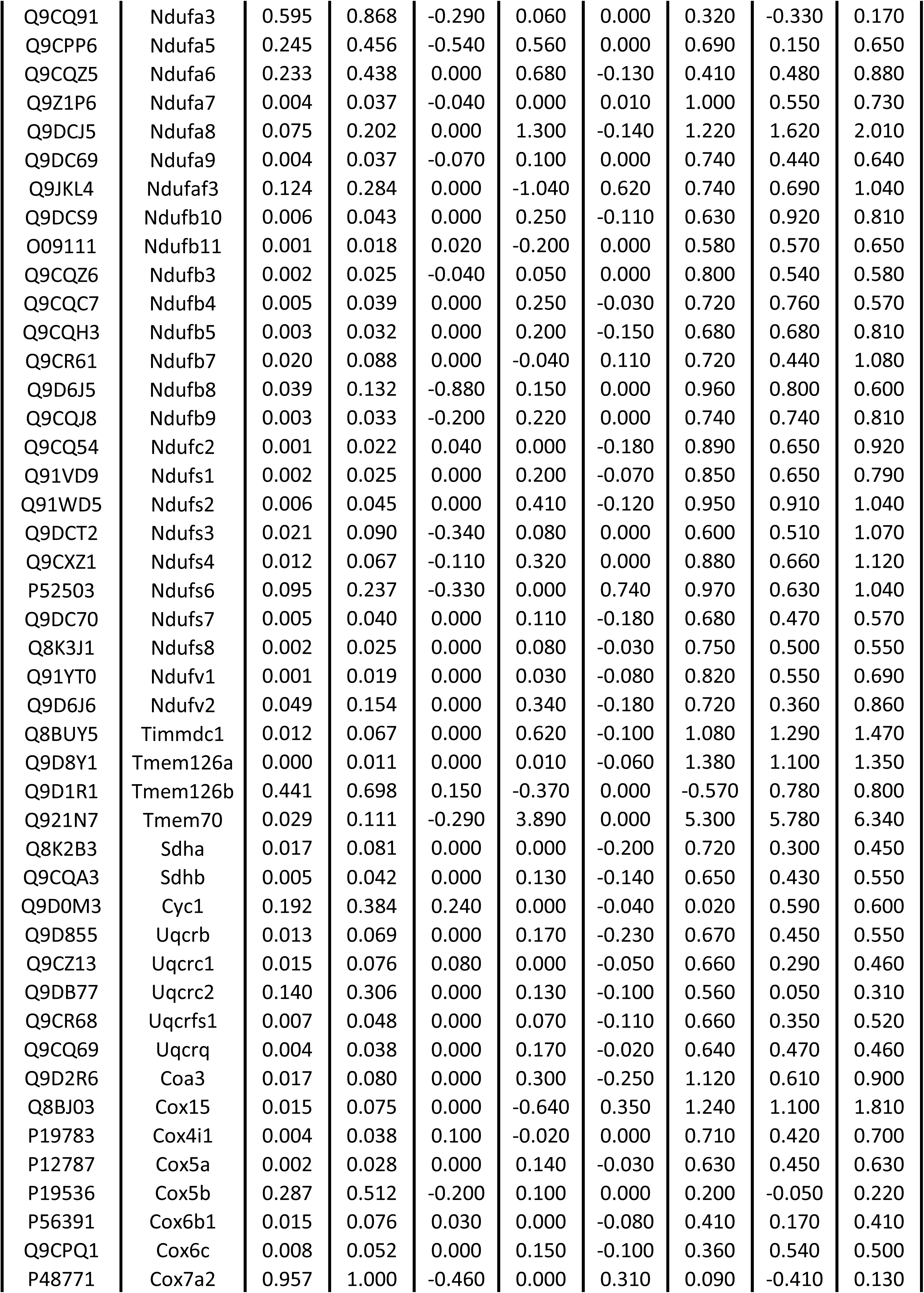
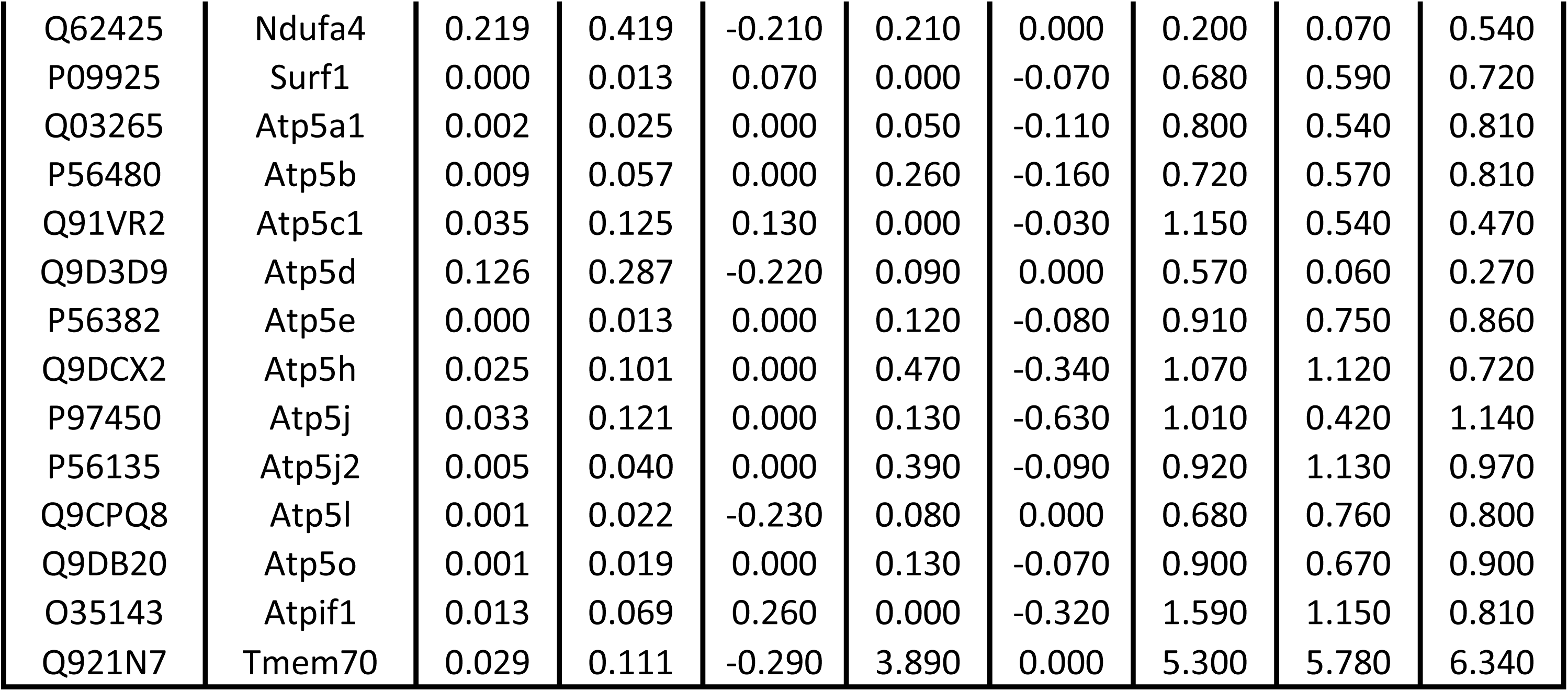
Proteins related to ETC were detected in CD4^+^ T cells isolated from old mice after mito-transfer and in non-manipulated CD4^+^ T cells from old mice. CD4^+^ T cells from young and old mice, and from old mice after mito-transfer were cultured for 4 h before processing for mass spectrometry analysis. Data expressed as median protein Log2 fold change of CD4^+^ T cells from old mice, from 3 individual old mice (paired experiment)

**Supp. Table 4.**
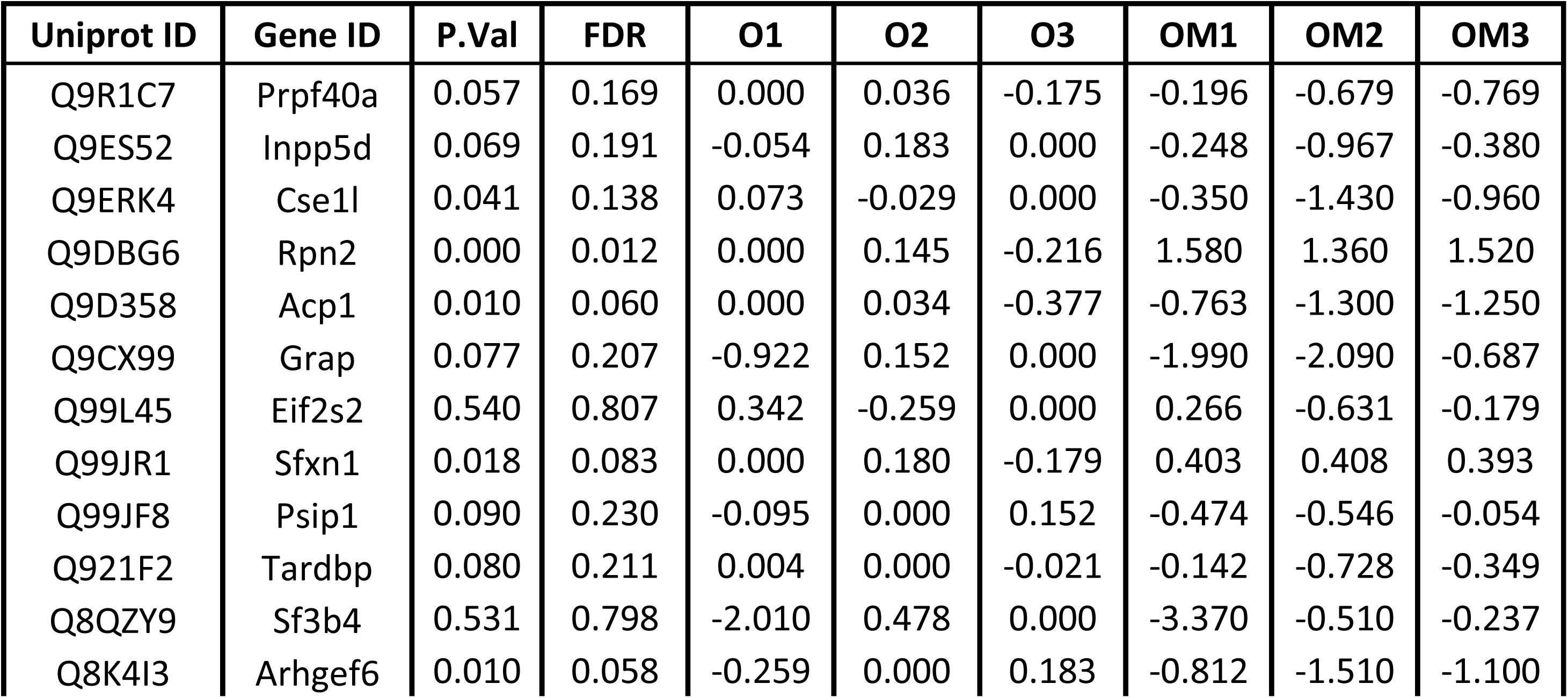
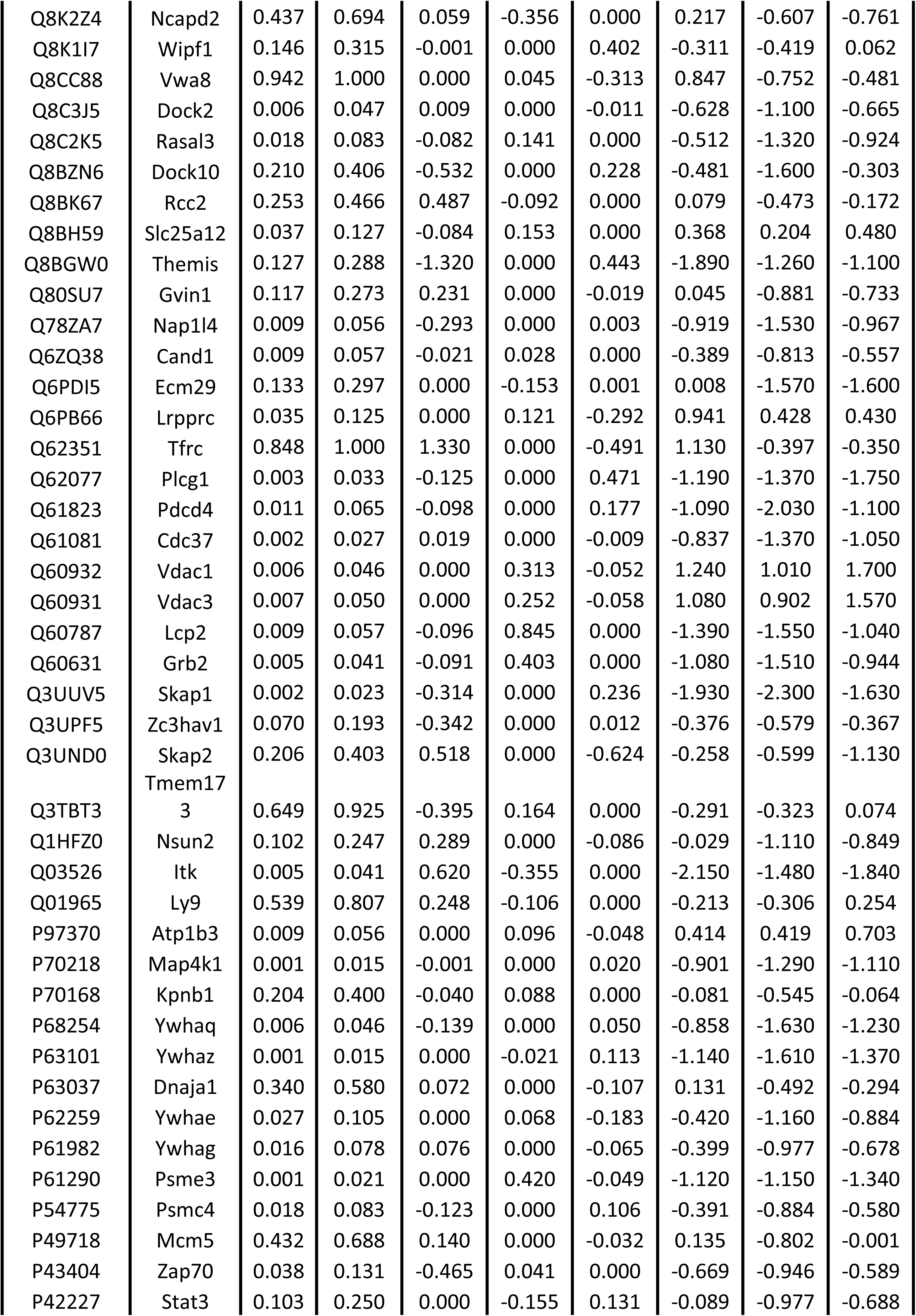
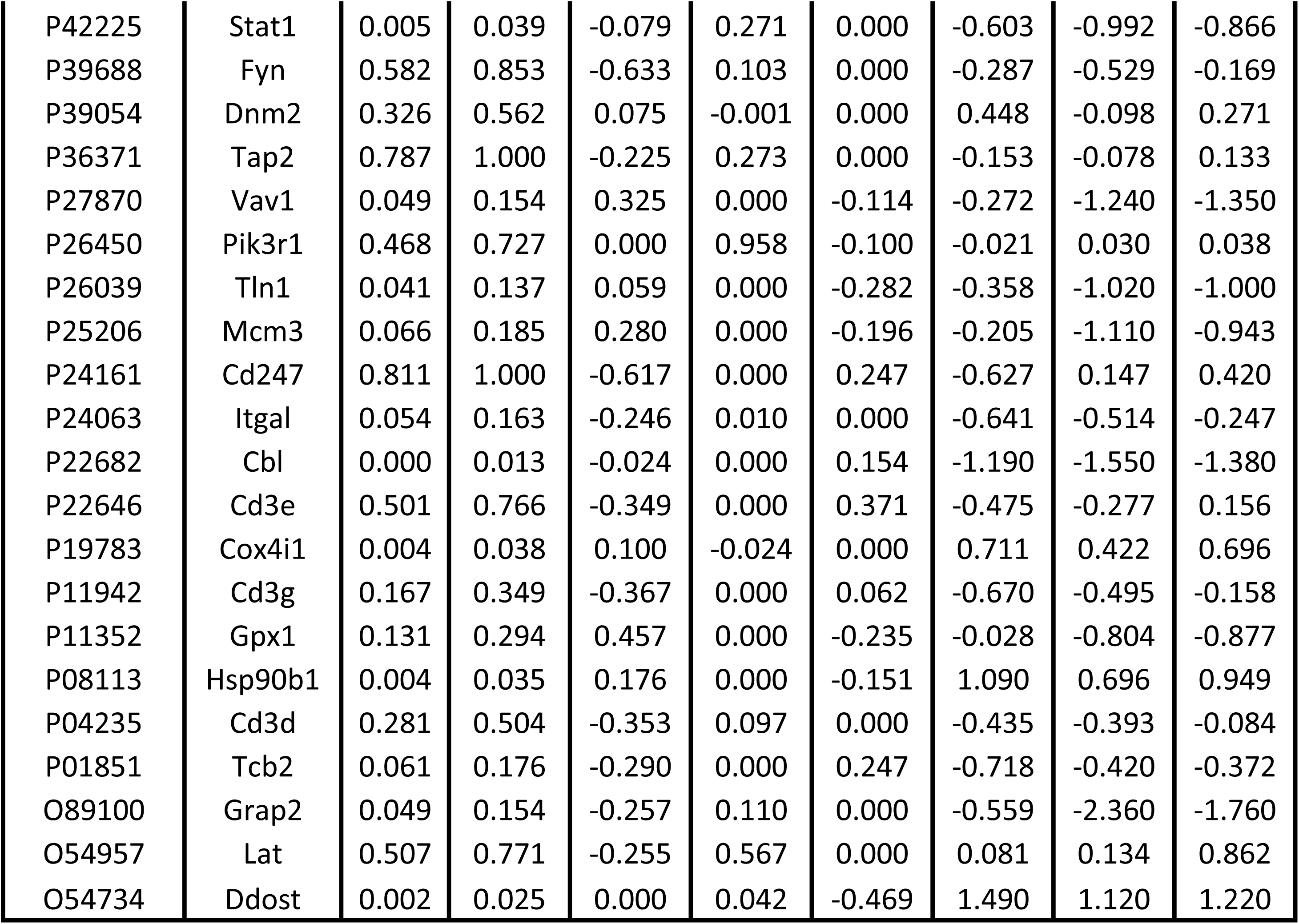
Proteins related to TCR signalosome were detected in CD4^+^ T cells isolated from old mice after mito-transfer and in non-manipulated CD4^+^ T cells from old mice. CD4^+^ T cells from young and old mice, and from old mice after mito-transfer were cultured for 4 h before processing for mass spectrometry analysis. Data expressed as median protein Log2 fold change of CD4^+^ T cells from old mice, from 3 individual old mice (paired experiment)

