## Supplemental Figures for "Extracellular Delivery of Functional Mitochondria Rescues the Dysfunction of CD4^+^ T Cells in Aging"

### Supplementary figures.

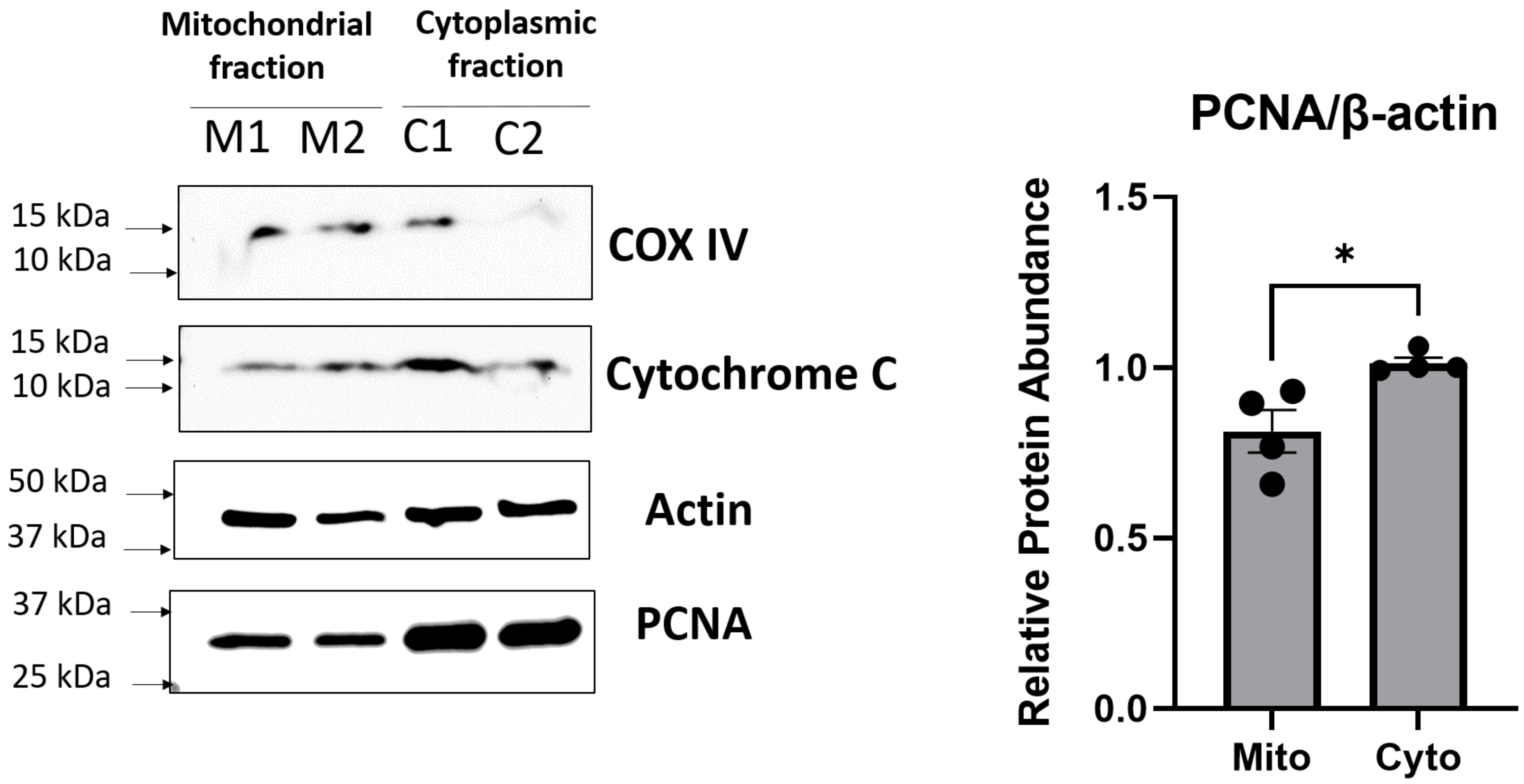
Mitochondrial fraction
Cytoplasmic fraction
M1
M2
C1
C2
15 kDa
10 kDa
COX IV
15 kDa
10 kDa
Cytochrome C
50 kDa
37 kDa
Actin
37 kDa
25 kDa
PCNA
PCNA/β-actin
Relative Protein Abundance
1.5
1.0
0.5
0.0
*
Mito
Cyto

**Supp. Fig. 1. Purity of mitochondrial isolation. A)** Representative wester blot images of probed mitochondrial and nuclear proteins in the mitochondrial fraction and cell fragments/unlysed cells after mitochondrial isolation. **B)** Relative amount of PCNA normalized to B-actin. (n=4), $p<0.05$=significant (*) using unpaired Student's t-test.

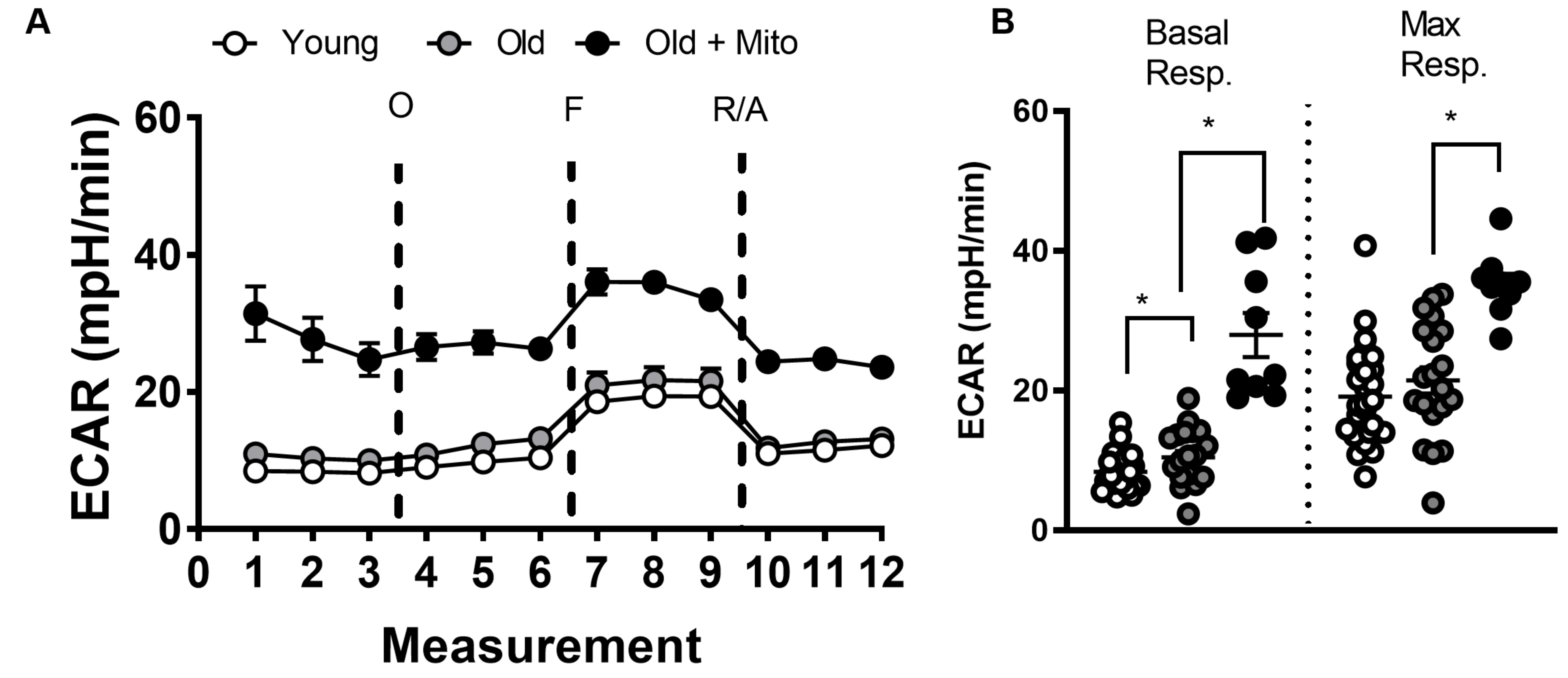

A
Young
Old
Old + Mito
ECAR (mpH/min)
60
40
20
0
O
F
R/A
0
1
2
3
4
5
6
7
8
9
10
11
12
Measurement
B
Basal Resp.
Max Resp.
ECAR (mpH/min)
60
40
20
0
*
*
*

**Supp. Fig. 2. Mito-transfer increases ECAR of $CD4^+$ T cells from old mice. A)** Kinetic curve of ECAR collected from mito-stress test assay, **B)** ECAR corresponding to basal and maximal respiration of $CD4^+$ T cells. 3-4 mice were used per group and experiments were replicated twice (n=3), $p<0.05$=significant (*) using unpaired Student's t-test.

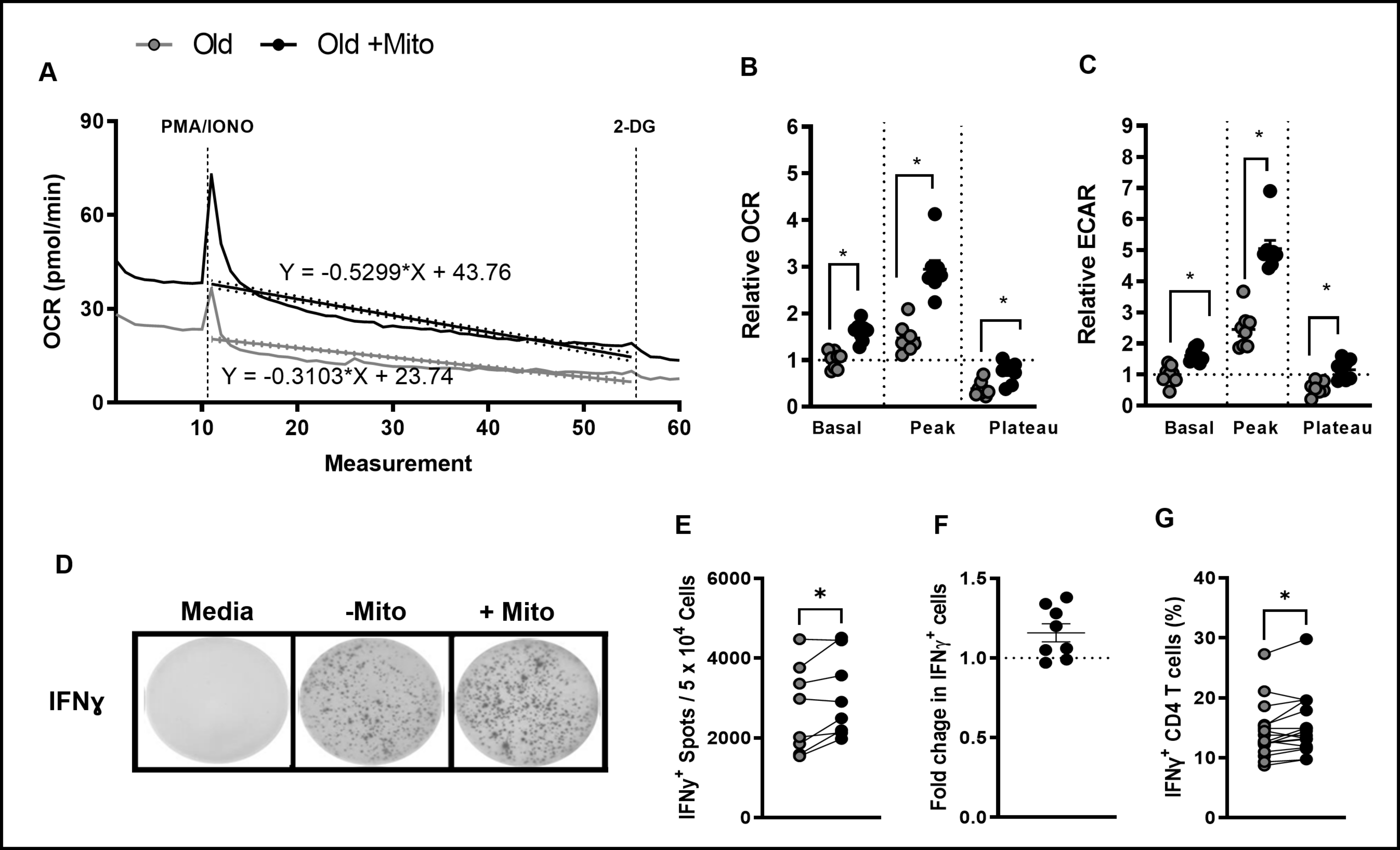

Old
Old +Mito
A
B
C
PMA/IONO
2-DG
OCR (pmol/min)
Y = -0.5299*X + 43.76
Y = -0.3103*X + 23.74
Measurement
Relative OCR
Relative ECAR
Basal
Peak
Plateau
D
Media
-Mito
+ Mito
IFNγ
E
F
G
IFNγ+ Spots / 5 x 10^4 Cells
Fold chage in IFNγ+ cells
IFNγ+ CD4 T cells (%)

**Supp. Fig. 3. Mito-transfer enhaced T cell activation induced metabolic switch and increased IFNγ$^+$ CD4$^+$ T cells. A)** The slope of OCR depression after acute injection of PMA/IONO in mito-transferred and non-manipulated CD4+ T cells from old mice. The relative change in **B)** OCR and **C)** ECAR of activated T cells (non-normalized to group). **D-G)** CD4$^+$ T cells from old mice with or without mito-transfer were stimulated with PMA/Ionomycin, after which various assays were used to quantify IFNγ production. **D)** Representative EliSpot images, **E)** the number IFNγ$^+$ producing T cells, and **F)** the fold change in IFNγ$^+$ producing CD4$^+$ T cells from old mice with or without mito-transfer, after stimulation with PMA/Ionomycin (24hr). The **G)** percent of IFNɣ$^+$ CD4$^+$ T cells via intracellular cytokine staining, in CD4$^+$ T cells from old mice with or without mito-transfer, after stimulation with PMA/Ionomycin (4h). 3-4 mice per group, repeated at least once ($n \geq 2$). $p < 0.05$ = significant (*) using paired Student's *t*-test.

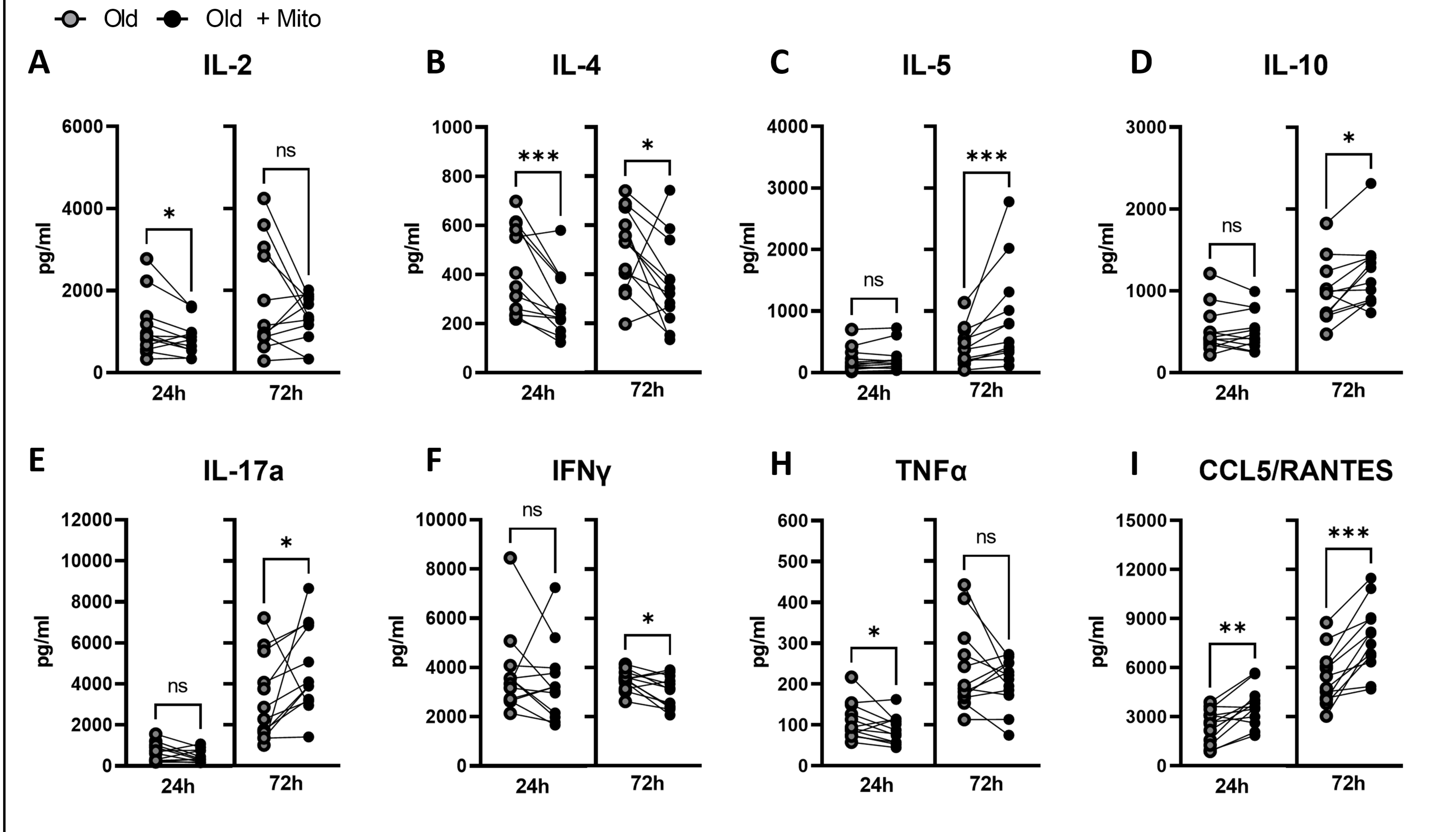

Old
Old + Mito
A
IL-2
B
IL-4
C
IL-5
D
IL-10
E
IL-17a
F
IFNγ
H
TNFα
I
CCL5/RANTES
pg/ml
24h
72h
ns
*
**
***

**Supp. Fig. 4. Mito-transfer alters T cell activation & cytokine production of $CD4^+$ T cells in old mice.** $CD4^+$ T cells from old mice with or without mito-transfer were stimulated with PMA/Ionomycin. After 24 and 72 h of stimulation with PMA/Ionomycin, the supernatants were examined by Luminex array for cytokines produced. $p \leq 0.05$ = *, $p \leq 0.01$ = **, or $p \leq 0.001$ = *** using paired Student's *t*-test.

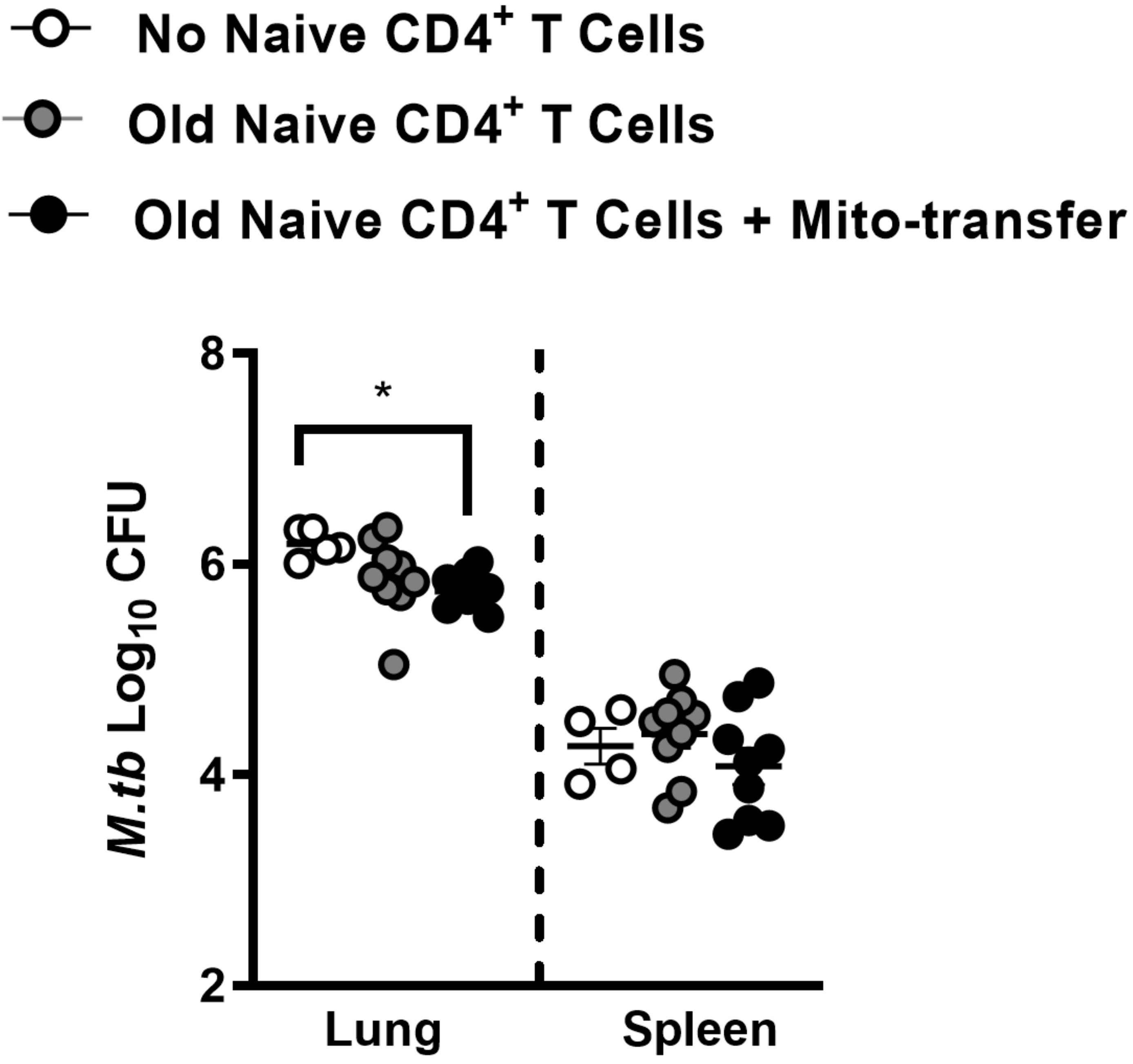

No Naive CD4+ T Cells
Old Naive CD4+ T Cells
Old Naive CD4+ T Cells + Mito-transfer
M.tb Log10 CFU
8
6
4
2
*
Lung
Spleen

**Supp. Fig. 5. Mito-transfer in naïve CD4 T cells from old mice protected mice against pathogens.** Naïve $CD4^+$ T cells from old mice with or without mito-transfer, or PBS were tail vein injected in Rag1-KO mice that were subsequently infected with *M.tb*. Lung and spleen CFU burden after 21 days of *M.tb* infection in Rag1-KO mice adoptively treated with 1x $10^6$ naïve $CD4^+$ T cells from old mice with or without mito-transfer. 4-8 mice per group, with $p \leq 0.05$ = *, using unpaired Student's *t*-test.

| Uniprot ID | Gene ID | P.Val | FDR | O1 | O2 | O3 | OM1 | OM2 | OM3 |
|---|---|---|---|---|---|---|---|---|---|
| P24270 | Cat | 0.046 | 0.148 | 0.177 | 0.000 | -0.253 | 1.040 | 0.403 | 0.494 |
| Q99LX0 | Park7 | 0.001 | 0.015 | 0.016 | -0.257 | 0.000 | -1.140 | -1.450 | -1.350 |
| P11247 | Mpo | 0.459 | 0.719 | 1.480 | 0.000 | -0.328 | 0.674 | -0.715 | -0.546 |
| Q9JMH6 | Txnrd1 | 0.030 | 0.114 | 0.000 | -0.115 | 0.093 | -0.408 | -1.270 | -1.320 |
| P08228 | Sod1 | 0.000 | 0.013 | -0.224 | 0.010 | 0.000 | -1.080 | -1.290 | -1.170 |
| Q9ERR7 | Sep15 | 0.054 | 0.164 | 0.000 | 0.000 | 0.112 | 0.582 | 0.658 | 0.160 |
| O08709 | Prdx6 | 0.003 | 0.031 | 0.085 | 0.000 | -0.156 | -1.090 | -1.810 | -1.670 |
| P19157 | Gstp1 | 0.004 | 0.036 | -0.138 | 0.000 | 0.106 | -1.050 | -1.380 | -0.809 |
| Q8CDN6 | Txnl1 | 0.008 | 0.051 | 0.000 | 0.040 | -0.283 | -0.969 | -1.380 | -0.821 |
| P11352 | Gpx1 | 0.131 | 0.294 | 0.457 | 0.000 | -0.235 | -0.028 | -0.804 | -0.877 |
| P99029 | Prdx5 | 0.996 | 1.000 | 0.234 | 0.000 | -0.266 | 0.051 | -0.201 | 0.121 |
| Q61171 | Prdx2 | 0.040 | 0.134 | 0.089 | 0.000 | -0.208 | -0.353 | -0.755 | -0.402 |
| P09671 | Sod2 | 0.023 | 0.096 | -0.061 | 0.348 | 0.000 | 0.715 | 0.476 | 0.682 |
| P02088 | Hbb-b1 | 0.050 | 0.155 | 0.499 | 0.000 | -0.660 | -1.080 | -1.670 | -3.060 |
| P35700 | Prdx1 | 0.130 | 0.293 | 0.028 | 0.000 | -0.030 | 0.421 | -0.006 | 0.325 |
| Q99LJ6 | Gpx7 | 0.000 | 0.012 | 0.150 | -1.130 | 0.000 | 5.360 | 4.910 | 5.340 |
| Q9D7B7 | Gpx8 | 0.000 | 0.007 | 0.000 | 0.540 | -0.171 | 5.920 | 5.450 | 5.940 |
| P31725 | S100a9 | 0.374 | 0.622 | 1.930 | 0.000 | -0.136 | 0.985 | -2.240 | -0.397 |
| P01942 | Hba | 0.040 | 0.135 | 0.473 | 0.000 | -0.794 | -0.999 | -1.950 | -2.470 |
| P49290 | Epx | 0.465 | 0.724 | 1.320 | -0.368 | 0.000 | 0.885 | -1.770 | -0.395 |
| P22437 | Ptgs1 | 0.902 | 1.000 | 0.376 | 0.000 | -0.608 | 0.313 | -0.174 | -0.239 |
| P62342 | Selt | 0.115 | 0.270 | 0.000 | 0.100 | -0.390 | 0.284 | 0.120 | 0.255 |
| Q91VS7 | Mgst1 | 0.030 | 0.113 | 0.999 | 0.000 | -0.245 | 1.580 | 1.500 | 2.410 |
| P47791 | Gsr | 0.018 | 0.084 | 0.341 | 0.000 | -0.118 | -0.466 | -1.080 | -1.080 |
| Q9JLT4 | Txnrd2 | 0.839 | 1.000 | 0.000 | 0.351 | -0.041 | 0.125 | 0.030 | 0.247 |
| Q6ZPY7 | Kdm3b | 0.027 | 0.107 | 0.000 | 0.495 | -0.124 | -0.530 | -1.600 | -1.240 |
| Q9CQM5 | Txndc17 | 0.008 | 0.055 | -0.143 | 0.362 | 0.000 | -1.500 | -1.760 | -2.830 |
| P08226 | Apoe | 0.508 | 0.772 | 0.861 | -0.299 | 0.000 | 0.960 | 0.161 | 0.360 |
| P20108 | Prdx3 | 0.006 | 0.046 | 0.214 | -0.058 | 0.000 | 0.778 | 0.509 | 0.675 |
| Q61646 | Hp | 0.202 | 0.399 | 1.520 | -0.351 | 0.000 | 0.109 | -1.160 | -0.961 |
| O08807 | Prdx4 | 0.256 | 0.470 | 0.038 | 0.000 | -0.002 | 0.439 | -0.070 | 0.262 |
| Q9DCM2 | Gstk1 | 0.701 | 0.977 | 0.010 | 0.000 | -0.212 | 0.377 | -0.571 | -0.378 |
| Q9BCZ4 | Vimp | 0.732 | 1.000 | -0.386 | 0.179 | 0.000 | 0.206 | -0.445 | -0.249 |
| Q9WU84 | Ccs | 0.004 | 0.036 | -0.087 | 0.000 | 0.263 | -0.735 | -1.230 | -1.050 |
| P30355 | Alox5ap | 0.424 | 0.678 | 0.786 | 0.000 | -0.301 | 1.160 | 0.291 | 0.218 |
| P02104 | Hbb-y | 0.451 | 0.708 | 0.641 | 0.000 | -2.510 | -0.842 | -1.560 | -2.020 |
| P97346 | Nxn | 0.333 | 0.570 | 0.333 | 0.000 | -1.090 | -2.170 | 0.304 | -1.790 |

| Uniprot | Gene ID | P.Val | FDR | O1 | O2 | O3 | OM1 | OM2 | OM3 |
|---|---|---|---|---|---|---|---|---|---|
| P24270 | Cat | 0.046 | 0.148 | 0.177 | 0.000 | -0.253 | 1.040 | 0.403 | 0.494 |
| P47738 | Aldh2 | 0.815 | 1.000 | 0.398 | 0.000 | -0.923 | 0.773 | -0.269 | -0.602 |

| | | | | | | | | | |
|---|---|---|---|---|---|---|---|---|---|
| Q8VDK1 | Nit1 | 0.269 | 0.488 | 0.029 | 0.000 | -0.100 | -0.074 | -0.053 | -0.391 |
| P47740 | Aldh3a2 | 0.057 | 0.169 | 0.000 | 0.097 | -0.121 | 0.310 | 0.102 | 0.328 |
| Q9JMH6 | Txnrd1 | 0.030 | 0.114 | 0.000 | -0.115 | 0.093 | -0.408 | -1.270 | -1.320 |
| P08228 | Sod1 | 0.000 | 0.013 | -0.224 | 0.010 | 0.000 | -1.080 | -1.290 | -1.170 |
| Q99J99 | Mpst | 0.334 | 0.571 | 0.163 | 0.000 | -0.160 | 0.378 | -0.932 | -0.891 |
| Q9JHW2 | Nit2 | 0.309 | 0.542 | -0.141 | 0.431 | 0.000 | 0.296 | 0.245 | 0.362 |
| O08709 | Prdx6 | 0.003 | 0.031 | 0.085 | 0.000 | -0.156 | -1.090 | -1.810 | -1.670 |
| P11352 | Gpx1 | 0.131 | 0.294 | 0.457 | 0.000 | -0.235 | -0.028 | -0.804 | -0.877 |
| P99029 | Prdx5 | 0.996 | 1.000 | 0.234 | 0.000 | -0.266 | 0.051 | -0.201 | 0.121 |
| Q61171 | Prdx2 | 0.040 | 0.134 | 0.089 | 0.000 | -0.208 | -0.353 | -0.755 | -0.402 |
| P09671 | Sod2 | 0.023 | 0.096 | -0.061 | 0.348 | 0.000 | 0.715 | 0.476 | 0.682 |
| Q64133 | Maoa | 0.000 | 0.011 | 0.151 | 0.000 | -0.761 | 3.640 | 3.580 | 3.770 |
| Q8R164 | Bphl | 0.031 | 0.114 | 0.197 | 0.000 | -0.133 | 0.714 | 0.305 | 0.778 |
| Q9DBF1 | Aldh7a1 | 0.000 | 0.009 | 0.000 | 0.318 | -0.191 | 2.700 | 2.460 | 2.520 |
| Q9WVL0 | Gstz1 | 0.112 | 0.265 | 0.000 | 0.163 | -0.024 | 1.190 | 0.487 | 0.239 |
| Q9DCN2 | Cyb5r3 | 0.000 | 0.005 | 0.000 | 0.161 | -0.051 | 2.080 | 1.950 | 2.120 |
| Q8BW75 | Maob | 0.000 | 0.013 | 0.667 | 0.000 | -0.370 | 4.120 | 3.720 | 4.010 |
| Q9CQX2 | Cyb5b | 0.013 | 0.071 | -0.164 | 0.064 | 0.000 | 0.975 | 0.423 | 0.947 |
| Q99KB8 | Hagh | 0.754 | 1.000 | 0.142 | 0.000 | -0.442 | -0.018 | -0.512 | -0.014 |
| Q9JLJ2 | Aldh9a1 | 0.081 | 0.213 | 0.030 | -0.088 | 0.000 | -0.239 | -1.410 | -0.779 |
| Q91VS7 | Mgst1 | 0.030 | 0.113 | 0.999 | 0.000 | -0.245 | 1.580 | 1.500 | 2.410 |
| P47791 | Gsr | 0.018 | 0.084 | 0.341 | 0.000 | -0.118 | -0.466 | -1.080 | -1.080 |
| Q9JLT4 | Txnrd2 | 0.839 | 1.000 | 0.000 | 0.351 | -0.041 | 0.125 | 0.030 | 0.247 |
| P20108 | Prdx3 | 0.006 | 0.046 | 0.214 | -0.058 | 0.000 | 0.778 | 0.509 | 0.675 |
| O08807 | Prdx4 | 0.256 | 0.470 | 0.038 | 0.000 | -0.002 | 0.439 | -0.070 | 0.262 |
| Q9D6Y7 | Msra | 0.068 | 0.189 | 0.549 | -0.217 | 0.000 | -0.252 | -0.768 | -0.814 |
| Q9DCM2 | Gstk1 | 0.701 | 0.977 | 0.010 | 0.000 | -0.212 | 0.377 | -0.571 | -0.378 |

**Supp. Table 1**. Antioxidant & mitochondrial detoxification proteins were detected in $CD4^+$ T cells isolated from old mice after mito-transfer and in non-manipulated $CD4^+$ T cells from old mice. $CD4^+$ T cells from young and old mice, and from old mice after mito-transfer were cultured for 4 h before processing for mass spectrometry analysis. Data expressed as median protein Log2 fold change of $CD4^+$ T cells from old mice, from 3 individual old mice (paired experiment)

| Uniprot ID | Gene ID | P. Val | FDR | O1 | O2 | O3 | OM1 | OM2 | OM3 |
|---|---|---|---|---|---|---|---|---|---|
| Q9JIK9 | Mrps34 | 0.049 | 0.154 | 0.000 | 0.757 | -0.497 | 1.210 | 1.140 | 1.010 |
| Q8K1Z0 | Coq9 | 0.032 | 0.117 | 0.000 | 0.186 | -0.023 | 0.862 | 0.388 | 0.472 |
| P97450 | Atp5j | 0.033 | 0.121 | 0.000 | 0.125 | -0.630 | 1.010 | 0.420 | 1.140 |

| | | | | | | | | | |
|---|---|---|---|---|---|---|---|---|---|
| Q9D0K2 | Oxct1 | 0.000 | 0.012 | 0.000 | 0.092 | -0.243 | 1.760 | 1.440 | 1.670 |
| P52825 | Cpt2 | 0.312 | 0.545 | 0.000 | 0.208 | -0.370 | 1.210 | 0.043 | 0.051 |
| Q9D6J6 | Ndufv2 | 0.049 | 0.154 | 0.000 | 0.342 | -0.183 | 0.718 | 0.363 | 0.856 |
| O08734 | Bak1 | 0.010 | 0.060 | 0.000 | 0.028 | -0.547 | 0.687 | 0.764 | 0.655 |
| P24270 | Cat | 0.046 | 0.148 | 0.177 | 0.000 | -0.253 | 1.040 | 0.403 | 0.494 |
| Q8BJZ4 | Mrps35 | 0.007 | 0.051 | 0.062 | 0.000 | -0.636 | 1.340 | 1.050 | 0.901 |
| D3Z7P3 | Gls | 0.500 | 0.765 | 0.000 | 0.006 | -0.037 | 0.732 | -0.191 | 0.043 |
| Q9JI39 | Abcb10 | 0.138 | 0.303 | 0.860 | -0.266 | 0.000 | 1.480 | 0.456 | 1.190 |
| Q9CQJ8 | Ndufb9 | 0.003 | 0.033 | -0.196 | 0.220 | 0.000 | 0.740 | 0.744 | 0.811 |
| P97742 | Cpt1a | 0.015 | 0.076 | 0.000 | 0.296 | -0.452 | 0.925 | 0.756 | 0.903 |
| P47738 | Aldh2 | 0.815 | 1.000 | 0.398 | 0.000 | -0.923 | 0.773 | -0.269 | -0.602 |
| Q9Z2I8 | Suclg2 | 0.566 | 0.835 | 0.026 | 0.000 | -0.159 | 0.510 | -0.235 | 0.016 |
| Q9DCX2 | Atp5h | 0.025 | 0.101 | 0.000 | 0.466 | -0.336 | 1.070 | 1.120 | 0.719 |
| Q8VE22 | Mrps23 | 0.006 | 0.044 | -0.054 | 0.132 | 0.000 | 0.889 | 0.501 | 0.810 |
| Q9D404 | Oxsm | 0.917 | 1.000 | 0.000 | 0.611 | -0.220 | 0.234 | 0.026 | 0.045 |
| Q99JB2 | Stoml2 | 0.007 | 0.048 | 0.000 | 0.206 | -0.196 | 0.991 | 0.626 | 0.828 |
| P59017 | Bcl2l13 | 0.118 | 0.274 | 0.000 | -0.077 | 0.194 | -0.067 | -0.784 | -0.362 |
| Q8BGH2 | Samm50 | 0.000 | 0.013 | 0.000 | 0.083 | -0.194 | 0.973 | 0.871 | 0.934 |
| Q9CZR8 | Tsfm | 0.707 | 0.983 | 0.245 | 0.000 | -0.207 | 0.426 | -0.475 | -0.282 |
| Q6PB66 | Lrpprc | 0.035 | 0.125 | 0.000 | 0.121 | -0.292 | 0.941 | 0.428 | 0.430 |
| O09111 | Ndufb11 | 0.001 | 0.018 | 0.019 | -0.197 | 0.000 | 0.578 | 0.575 | 0.648 |
| Q9D6R2 | Idh3a | 0.546 | 0.814 | 0.000 | 0.280 | -0.166 | 0.748 | -0.097 | 0.040 |
| Q9CQ92 | Fis1 | 0.078 | 0.209 | 0.240 | -0.067 | 0.000 | -0.226 | -0.500 | -0.127 |
| Q78PY7 | Snd1 | 0.431 | 0.687 | 0.121 | 0.000 | -0.224 | 0.151 | -0.543 | -0.309 |
| P29758 | Oat | 0.001 | 0.018 | 0.000 | 0.170 | -0.473 | 1.810 | 1.920 | 2.200 |
| O08749 | Dld | 0.005 | 0.039 | 0.000 | 0.188 | -0.179 | 0.904 | 0.625 | 0.763 |
| P38647 | Hspa9 | 0.072 | 0.196 | 0.035 | 0.000 | -0.356 | 1.510 | 0.407 | 0.498 |
| P63038 | Hspd1 | 0.006 | 0.044 | 0.000 | 0.119 | -0.240 | 1.090 | 0.650 | 0.822 |
| Q9DBL1 | Acadsb | 0.001 | 0.022 | 0.000 | 0.020 | -0.201 | 1.980 | 1.390 | 1.380 |
| Q9Z2I0 | Letm1 | 0.002 | 0.025 | 0.000 | 0.057 | -0.198 | 0.565 | 0.495 | 0.591 |
| Q07813 | Bax | 0.323 | 0.559 | -0.110 | 0.000 | 0.085 | 0.671 | -0.114 | 0.210 |
| Q99JY0 | Hadhb | 0.462 | 0.721 | -0.070 | 0.350 | 0.000 | 0.625 | -0.024 | 0.237 |
| Q9Z110 | Aldh18a1 | 0.109 | 0.261 | 0.000 | 0.052 | -0.331 | 1.350 | 0.197 | 0.449 |
| Q8VDK1 | Nit1 | 0.269 | 0.488 | 0.029 | 0.000 | -0.100 | -0.074 | -0.053 | -0.391 |
| Q9CPY7 | Lap3 | 0.806 | 1.000 | 0.158 | 0.000 | -0.383 | 0.568 | -0.516 | 0.000 |
| Q8BH59 | Slc25a12 | 0.037 | 0.127 | -0.084 | 0.153 | 0.000 | 0.368 | 0.204 | 0.480 |
| Q9R0X4 | Acot9 | 0.058 | 0.171 | 0.000 | 0.694 | -0.021 | 1.030 | 0.770 | 0.829 |
| Q8BWT1 | Acaa2 | 0.089 | 0.228 | 0.000 | 0.088 | -0.212 | 0.592 | 0.136 | 0.245 |
| Q99N84 | Mrps18b | 0.001 | 0.019 | -0.100 | 0.262 | 0.000 | 1.290 | 1.020 | 1.220 |
| Q99LX0 | Park7 | 0.001 | 0.015 | 0.016 | -0.257 | 0.000 | -1.140 | -1.450 | -1.350 |
| Q9CQ06 | Mrpl24 | 0.031 | 0.115 | 0.158 | 0.000 | -0.175 | 0.302 | 0.820 | 0.865 |
| P47740 | Aldh3a2 | 0.057 | 0.169 | 0.000 | 0.097 | -0.121 | 0.310 | 0.102 | 0.328 |
| Q9CPV4 | Glod4 | 0.023 | 0.096 | 0.000 | 0.150 | -0.049 | -0.326 | -0.824 | -0.470 |
| Q9D0L7 | Armc10 | 0.027 | 0.107 | -0.131 | 0.052 | 0.000 | -0.301 | -0.431 | -0.208 |
| Q9CR61 | Ndufb7 | 0.020 | 0.088 | 0.000 | -0.043 | 0.113 | 0.717 | 0.435 | 1.080 |

| | | | | | | | | | |
|---|---|---|---|---|---|---|---|---|---|
| Q9DC69 | Ndufa9 | 0.004 | 0.037 | -0.072 | 0.102 | 0.000 | 0.741 | 0.442 | 0.635 |
| Q9CQN1 | Trap1 | 0.459 | 0.719 | 0.170 | 0.000 | -0.139 | 0.175 | -0.335 | -0.254 |
| Q9Z2Z6 | Slc25a20 | 0.006 | 0.043 | 0.161 | 0.000 | -0.081 | 0.953 | 0.633 | 0.619 |
| Q921H8 | Acaa1a | 0.168 | 0.349 | 0.077 | 0.000 | -0.377 | 0.732 | -0.098 | 0.503 |
| Q9ERS2 | Ndufa13 | 0.056 | 0.168 | 0.038 | -0.179 | 0.000 | 0.652 | 0.143 | 0.876 |
| P19783 | Cox4i1 | 0.004 | 0.038 | 0.100 | -0.024 | 0.000 | 0.711 | 0.422 | 0.696 |
| Q9CZ42 | Naxd | 0.001 | 0.018 | 0.148 | 0.000 | -0.050 | 1.030 | 0.765 | 0.960 |
| Q99LC3 | Ndufa10 | 0.006 | 0.047 | 0.000 | 0.010 | -0.249 | 0.718 | 0.460 | 0.860 |
| Q8R2Y8 | Ptrh2 | 0.015 | 0.077 | -0.265 | 0.190 | 0.000 | 1.240 | 1.150 | 2.290 |
| Q8R4N0 | Clybl | 0.061 | 0.177 | 0.182 | 0.000 | -0.138 | 0.704 | 0.176 | 0.694 |
| O88696 | Clpp | 0.016 | 0.078 | 0.000 | 0.043 | -0.364 | 0.863 | 0.401 | 0.665 |
| Q80Y81 | Elac2 | 0.624 | 0.899 | 0.000 | -0.208 | 0.155 | 0.131 | -0.831 | 0.116 |
| Q9CRD0 | Ociad1 | 0.184 | 0.372 | -0.379 | 0.247 | 0.000 | -0.636 | -0.622 | -0.097 |
| Q64105 | Spr | 0.003 | 0.033 | -0.370 | 0.096 | 0.000 | -1.650 | -1.710 | -1.160 |
| P09528 | Fth1 | 0.085 | 0.220 | 0.442 | -0.233 | 0.000 | -0.225 | -1.190 | -0.715 |
| Q9DB77 | Uqcrc2 | 0.140 | 0.306 | 0.000 | 0.125 | -0.097 | 0.556 | 0.046 | 0.312 |
| Q61425 | Hadh | 0.375 | 0.623 | -0.733 | 0.214 | 0.000 | 0.578 | 0.284 | -0.255 |
| Q99MN9 | Pccb | 0.003 | 0.032 | 0.000 | 0.238 | -0.023 | 0.894 | 0.712 | 1.040 |
| Q8BMS1 | Hadha | 0.932 | 1.000 | 0.027 | 0.000 | -0.269 | 0.492 | -0.475 | -0.178 |
| P38060 | Hmgcl | 0.061 | 0.177 | 0.000 | 0.148 | -0.178 | 0.592 | 0.202 | 0.328 |
| Q9CZ13 | Uqcrc1 | 0.015 | 0.076 | 0.081 | 0.000 | -0.051 | 0.660 | 0.292 | 0.464 |
| Q922W5 | Pycr1 | 0.000 | 0.014 | 0.208 | 0.000 | -0.219 | 2.510 | 1.920 | 2.420 |
| P97287 | Mcl1 | 0.225 | 0.427 | 0.197 | 0.000 | -0.777 | 0.487 | -0.153 | 1.300 |
| P50544 | Acadvl | 0.245 | 0.456 | 0.000 | 0.061 | -0.285 | 0.866 | -0.022 | 0.132 |
| Q8CAQ8 | Immt | 0.003 | 0.032 | 0.000 | 0.109 | -0.145 | 0.915 | 0.583 | 0.795 |
| P56480 | Atp5b | 0.009 | 0.057 | 0.000 | 0.258 | -0.162 | 0.723 | 0.568 | 0.808 |
| Q9CYG7 | Tomm34 | 0.013 | 0.071 | -0.013 | 0.210 | 0.000 | -0.338 | -0.800 | -0.588 |
| O35857 | Timm44 | 0.016 | 0.078 | 0.231 | 0.000 | -0.162 | 0.827 | 0.450 | 0.765 |
| Q8CC88 | Vwa8 | 0.942 | 1.000 | 0.000 | 0.045 | -0.313 | 0.847 | -0.752 | -0.481 |
| Q91VD9 | Ndufs1 | 0.002 | 0.025 | 0.000 | 0.197 | -0.065 | 0.847 | 0.648 | 0.786 |
| Q8BFR5 | Tufm | 0.440 | 0.697 | 0.165 | 0.000 | -0.339 | 0.256 | -0.698 | -0.598 |
| Q14C51 | Ptcd3 | 0.001 | 0.019 | 0.000 | 0.011 | -0.187 | 0.717 | 0.554 | 0.741 |
| Q9QX60 | Dguok | 0.708 | 0.984 | -0.077 | 0.150 | 0.000 | 0.044 | -0.050 | 0.196 |
| Q9JJL8 | Sars2 | 0.446 | 0.703 | 0.281 | 0.000 | -0.050 | 0.559 | -0.022 | 0.196 |
| Q99JR1 | Sfxn1 | 0.018 | 0.083 | 0.000 | 0.180 | -0.179 | 0.403 | 0.408 | 0.393 |
| Q80Y14 | Glrx5 | 0.112 | 0.265 | 0.079 | 0.000 | -0.218 | 0.910 | 0.185 | 0.266 |
| P50136 | Bckdha | 0.099 | 0.243 | 0.030 | 0.000 | -0.341 | 0.421 | 0.014 | 0.389 |
| P54071 | Idh2 | 0.004 | 0.036 | 0.054 | 0.000 | -0.276 | 1.440 | 0.889 | 0.986 |
| Q9WTP7 | Ak3 | 0.037 | 0.129 | -0.146 | 0.101 | 0.000 | 0.809 | 0.233 | 0.598 |
| Q6NVE9 | Pptc7 | 0.557 | 0.825 | 0.000 | -0.054 | 0.016 | 0.205 | -0.490 | -0.140 |
| Q7TMF3 | Ndufa12 | 0.011 | 0.064 | -0.161 | 0.000 | 0.141 | 0.792 | 0.547 | 1.110 |
| O35465 | Fkbp8 | 0.521 | 0.786 | 0.000 | 0.167 | -0.029 | 0.317 | -0.227 | -0.452 |
| Q60932 | Vdac1 | 0.006 | 0.046 | 0.000 | 0.313 | -0.052 | 1.240 | 1.010 | 1.700 |
| Q60597 | Ogdh | 0.007 | 0.048 | 0.000 | 0.044 | -0.239 | 0.747 | 0.432 | 0.587 |
| Q78IK4 | Apool | 0.010 | 0.060 | -0.061 | 0.205 | 0.000 | 0.712 | 0.459 | 0.823 |

| | | | | | | | | | |
|---|---|---|---|---|---|---|---|---|---|
| Q91YT0 | Ndufv1 | 0.001 | 0.019 | 0.000 | 0.034 | -0.076 | 0.817 | 0.553 | 0.688 |
| Q91ZA3 | Pcca | 0.004 | 0.037 | 0.000 | 0.273 | -0.086 | 0.822 | 0.667 | 0.847 |
| Q3UV70 | Pdp1 | 0.009 | 0.056 | -0.168 | 0.000 | 0.071 | 0.763 | 0.401 | 0.775 |
| P51174 | Acadl | 0.039 | 0.132 | 0.000 | 0.343 | -0.012 | 0.526 | 0.424 | 0.475 |
| Q8K4Z3 | Naxe | 0.013 | 0.071 | 0.012 | 0.000 | -0.125 | 0.324 | 0.340 | 0.156 |
| Q99KI0 | Aco2 | 0.135 | 0.299 | 0.000 | 0.042 | -0.166 | 0.553 | 0.087 | 0.138 |
| P26443 | Glud1 | 0.007 | 0.048 | -0.208 | 0.207 | 0.000 | 0.756 | 0.579 | 0.813 |
| Q8BMF4 | Dlat | 0.002 | 0.025 | 0.000 | 0.245 | -0.070 | 1.060 | 0.823 | 1.080 |
| Q9Z2I9 | Sucla2 | 0.144 | 0.311 | 0.045 | 0.000 | -0.226 | 0.968 | 0.063 | 0.320 |
| Q9DCS9 | Ndufb10 | 0.006 | 0.043 | 0.000 | 0.250 | -0.106 | 0.628 | 0.918 | 0.805 |
| Q99MN1 | Kars | 0.095 | 0.237 | 0.000 | 0.246 | -0.487 | -0.474 | -0.558 | -0.660 |
| Q8R123 | Flad1 | 0.186 | 0.376 | 0.107 | 0.000 | -0.001 | -0.107 | -0.914 | -0.144 |
| Q9JMH6 | Txnrd1 | 0.030 | 0.114 | 0.000 | -0.115 | 0.093 | -0.408 | -1.270 | -1.320 |
| Q9D2G2 | Dlst | 0.003 | 0.033 | 0.000 | 0.150 | -0.204 | 0.862 | 0.626 | 0.867 |
| Q80ZK0 | Mrps10 | 0.027 | 0.107 | 0.000 | 0.562 | -0.484 | 1.350 | 0.904 | 1.690 |
| Q9CRB9 | Chchd3 | 0.005 | 0.039 | -0.052 | 0.233 | 0.000 | 0.868 | 0.646 | 1.000 |
| Q9DAM5 | Slc25a19 | 0.512 | 0.776 | 0.000 | 0.235 | -0.215 | 0.146 | 0.162 | -1.530 |
| Q99J39 | Mlycd | 0.122 | 0.280 | 0.000 | 0.662 | -0.127 | 0.733 | 0.678 | 0.583 |
| P08228 | Sod1 | 0.000 | 0.013 | -0.224 | 0.010 | 0.000 | -1.080 | -1.290 | -1.170 |
| Q99LC5 | Etfa | 0.310 | 0.542 | 0.000 | 0.213 | -0.198 | 0.705 | -0.164 | 0.467 |
| Q8BH04 | Pck2 | 0.009 | 0.056 | 0.068 | 0.000 | -0.323 | 1.320 | 0.741 | 0.823 |
| Q9Z0V7 | Timm17b | 0.705 | 0.982 | 0.000 | -0.059 | 0.096 | 0.339 | -0.298 | 0.243 |
| Q9WTQ8 | Timm23 | 0.001 | 0.017 | 0.000 | 0.198 | -0.057 | 0.799 | 0.774 | 0.865 |
| P28352 | Apex1 | 0.028 | 0.107 | 0.000 | -0.288 | 0.047 | -0.452 | -0.881 | -0.582 |
| Q80XN0 | Bdh1 | 0.208 | 0.404 | 0.000 | 0.147 | -0.057 | 0.047 | -0.615 | -0.246 |
| Q8BVI4 | Qdpr | 0.149 | 0.319 | 0.000 | -0.293 | 0.016 | -0.220 | -0.562 | -0.276 |
| Q9CQ62 | Decr1 | 0.431 | 0.687 | -0.053 | 0.110 | 0.000 | 0.728 | -0.038 | 0.024 |
| Q9WV85 | Nme3 | 0.093 | 0.234 | 0.001 | -0.129 | 0.000 | -0.343 | -0.578 | -0.122 |
| Q99J99 | Mpst | 0.334 | 0.571 | 0.163 | 0.000 | -0.160 | 0.378 | -0.932 | -0.891 |
| Q3UQ84 | Tars2 | 0.008 | 0.053 | 0.000 | 0.010 | -0.095 | 0.543 | 0.329 | 0.293 |
| Q8CD10 | Micu2 | 0.000 | 0.014 | -0.023 | 0.080 | 0.000 | 1.030 | 0.770 | 0.952 |
| Q8JZN5 | Acad9 | 0.215 | 0.412 | 0.000 | 0.153 | -0.210 | 0.571 | 0.123 | 0.083 |
| Q91V61 | Sfxn3 | 0.006 | 0.043 | -0.166 | 0.192 | 0.000 | 0.721 | 0.546 | 0.762 |
| Q91WD5 | Ndufs2 | 0.006 | 0.045 | 0.000 | 0.406 | -0.118 | 0.946 | 0.914 | 1.040 |
| Q8BKZ9 | Pdhx | 0.019 | 0.086 | 0.000 | 0.072 | -0.571 | 0.880 | 0.548 | 0.654 |
| Q8VCF0 | Mavs | 0.900 | 1.000 | -0.276 | 0.000 | 0.068 | -0.034 | -0.283 | 0.051 |
| Q60930 | Vdac2 | 0.002 | 0.023 | 0.000 | 0.161 | -0.063 | 1.040 | 0.946 | 1.360 |
| Q8K009 | Aldh1l2 | 0.000 | 0.014 | 0.838 | 0.000 | -0.233 | 4.520 | 3.900 | 4.030 |
| Q3UJU9 | Rmdn3 | 0.332 | 0.570 | 0.215 | 0.000 | -0.020 | -0.077 | -0.868 | 0.115 |
| Q61941 | Nnt | 0.000 | 0.013 | 1.030 | -0.638 | 0.000 | 5.910 | 5.440 | 5.700 |
| Q9D0S9 | Hint2 | 0.816 | 1.000 | 0.000 | 1.130 | -1.100 | -0.268 | 1.220 | -0.316 |
| Q921S7 | Mrpl37 | 0.249 | 0.462 | 0.174 | 0.000 | -0.144 | 0.783 | -0.030 | 0.302 |
| Q3V3R1 | Mthfd1l | 0.570 | 0.839 | 0.201 | 0.000 | -0.215 | 0.497 | -0.751 | -0.501 |
| Q8K411 | Pitrm1 | 0.003 | 0.030 | 0.219 | 0.000 | -0.224 | 1.320 | 0.923 | 1.190 |
| Q9D1L0 | Chchd2 | 0.596 | 0.869 | -0.345 | 0.000 | 0.075 | 0.112 | -1.140 | 0.029 |

| | | | | | | | | | |
|---|---|---|---|---|---|---|---|---|---|
| Q9QYR9 | Acot2 | 0.001 | 0.018 | -0.148 | 0.041 | 0.000 | 0.854 | 0.676 | 0.934 |
| Q9JHW2 | Nit2 | 0.309 | 0.542 | -0.141 | 0.431 | 0.000 | 0.296 | 0.245 | 0.362 |
| Q9DB20 | Atp5o | 0.001 | 0.019 | 0.000 | 0.129 | -0.067 | 0.904 | 0.673 | 0.901 |
| O08709 | Prdx6 | 0.003 | 0.031 | 0.085 | 0.000 | -0.156 | -1.090 | -1.810 | -1.670 |
| Q8BIJ6 | Iars2 | 0.008 | 0.053 | 0.000 | 0.073 | -0.104 | 0.819 | 0.408 | 0.655 |
| Q9DC61 | Pmpca | 0.012 | 0.068 | 0.000 | 0.351 | -0.158 | 1.180 | 0.802 | 0.788 |
| Q99L13 | Hibadh | 0.000 | 0.014 | 0.000 | 0.215 | -0.163 | 1.320 | 1.240 | 1.450 |
| Q03265 | Atp5a1 | 0.002 | 0.025 | 0.000 | 0.050 | -0.113 | 0.804 | 0.540 | 0.809 |
| Q9QXX4 | Slc25a13 | 0.004 | 0.038 | -0.014 | 0.047 | 0.000 | 0.518 | 0.393 | 0.699 |
| O35855 | Bcat2 | 0.331 | 0.569 | 0.049 | 0.000 | -0.484 | 0.516 | -0.134 | 0.039 |
| Q9CQC7 | Ndufb4 | 0.005 | 0.039 | 0.000 | 0.250 | -0.034 | 0.719 | 0.760 | 0.569 |
| Q8CGK3 | Lonp1 | 0.159 | 0.335 | 0.242 | 0.000 | -0.158 | 1.280 | 0.204 | 0.415 |
| Q8VEM8 | Slc25a3 | 0.001 | 0.019 | 0.000 | 0.026 | -0.130 | 1.170 | 0.803 | 0.950 |
| Q9D051 | Pdhb | 0.332 | 0.570 | 0.000 | 0.004 | -0.255 | 0.792 | -0.134 | 0.064 |
| Q9CYR0 | Ssbp1 | 0.351 | 0.592 | 0.000 | 0.077 | 0.000 | 0.385 | -0.069 | 0.185 |
| Q64521 | Gpd2 | 0.010 | 0.061 | 0.000 | 0.125 | -0.395 | 0.903 | 0.600 | 0.693 |
| Q9Z0X1 | Aifm1 | 0.917 | 1.000 | 0.000 | 0.036 | -0.025 | 0.160 | -0.134 | 0.015 |
| Q8K1R3 | Pnpt1 | 0.333 | 0.571 | 0.128 | 0.000 | -0.456 | 1.050 | 0.609 | -0.428 |
| Q91V92 | Acly | 0.836 | 1.000 | 0.521 | 0.000 | -0.217 | 0.528 | -0.126 | 0.096 |
| Q9WUM5 | Suclg1 | 0.579 | 0.850 | 0.279 | 0.000 | -0.064 | 1.160 | -0.031 | -0.140 |
| Q8K2B3 | Sdha | 0.017 | 0.081 | 0.000 | 0.002 | -0.204 | 0.722 | 0.297 | 0.454 |
| P67778 | Phb | 0.003 | 0.033 | 0.000 | 0.017 | -0.309 | 0.843 | 0.579 | 0.735 |
| Q922Q4 | Pycr2 | 0.016 | 0.078 | 0.102 | 0.000 | -0.106 | 1.190 | 0.520 | 0.764 |
| Q61102 | Abcb7 | 0.585 | 0.857 | -0.410 | 0.169 | 0.000 | 0.145 | -0.268 | 0.316 |
| P47802 | Mtx1 | 0.295 | 0.522 | 0.000 | 0.014 | -0.282 | 0.283 | -0.076 | 0.042 |
| P08249 | Mdh2 | 0.023 | 0.096 | -0.067 | 0.250 | 0.000 | 0.700 | 0.373 | 0.737 |
| P42125 | Eci1 | 0.192 | 0.384 | 0.000 | 0.429 | -0.075 | 0.361 | 0.416 | 0.326 |
| O35143 | Atpif1 | 0.013 | 0.069 | 0.259 | 0.000 | -0.321 | 1.590 | 1.150 | 0.813 |
| P84084 | Arf5 | 0.092 | 0.233 | 0.000 | 0.100 | -0.067 | -0.116 | -1.250 | -0.806 |
| O35972 | Mrpl23 | 0.003 | 0.033 | 0.000 | 0.107 | -0.240 | 0.886 | 0.665 | 0.669 |
| Q9D0D3 | Mtpap | 0.039 | 0.134 | -0.042 | 0.000 | 0.411 | 0.553 | 0.544 | 0.831 |
| Q9D6S7 | Mrrf | 0.086 | 0.223 | 0.000 | 0.433 | -0.651 | 0.785 | 0.917 | 0.434 |
| Q9CXT8 | Pmpcb | 0.017 | 0.082 | 0.000 | 0.156 | -0.441 | 1.050 | 0.643 | 0.643 |
| Q8CHT0 | Aldh4a1 | 0.875 | 1.000 | 0.335 | -0.067 | 0.000 | 1.090 | -0.715 | -0.397 |
| Q9D1N9 | Mrpl21 | 0.001 | 0.019 | 0.000 | 0.089 | -0.218 | 0.892 | 0.811 | 1.010 |
| Q8QZT1 | Acat1 | 0.050 | 0.156 | 0.000 | 0.181 | -0.006 | 0.765 | 0.228 | 0.592 |
| Q9DCS3 | Mecr | 0.121 | 0.279 | 0.000 | 0.207 | -0.206 | 1.210 | 0.269 | 0.392 |
| Q3ULD5 | Mccc2 | 0.004 | 0.036 | 0.000 | 0.315 | -0.197 | 1.080 | 0.893 | 0.981 |
| Q9DCC8 | Tomm20 | 0.641 | 0.917 | -0.090 | 0.350 | 0.000 | 0.068 | 0.058 | 0.398 |
| P55096 | Abcd3 | 0.092 | 0.232 | 0.042 | 0.000 | -0.929 | 1.360 | 1.060 | 0.053 |
| P32020 | Scp2 | 0.103 | 0.248 | 0.172 | 0.000 | -0.126 | 0.611 | 0.091 | 0.465 |
| P19096 | Fasn | 0.558 | 0.826 | 0.157 | 0.000 | -0.066 | 0.282 | -0.464 | -0.161 |
| Q8BH95 | Echs1 | 0.035 | 0.124 | 0.000 | 0.240 | -0.097 | 0.915 | 0.414 | 0.524 |
| P56391 | Cox6b1 | 0.015 | 0.076 | 0.029 | 0.000 | -0.083 | 0.406 | 0.174 | 0.409 |
| Q99N96 | Mrpl1 | 0.091 | 0.231 | 0.000 | -1.980 | 0.201 | 0.820 | 0.852 | 1.270 |

| | | | | | | | | | |
|---|---|---|---|---|---|---|---|---|---|
| Q99KE1 | Me2 | 0.952 | 1.000 | 0.000 | 0.140 | -0.205 | 0.190 | -0.212 | -0.072 |
| Q60931 | Vdac3 | 0.007 | 0.050 | 0.000 | 0.252 | -0.058 | 1.080 | 0.902 | 1.570 |
| P09925 | Surf1 | 0.000 | 0.013 | 0.069 | 0.000 | -0.073 | 0.682 | 0.590 | 0.722 |
| Q8BMD8 | Slc25a24 | 0.023 | 0.096 | -0.447 | 0.176 | 0.000 | 0.526 | 0.574 | 0.757 |
| P11352 | Gpx1 | 0.131 | 0.294 | 0.457 | 0.000 | -0.235 | -0.028 | -0.804 | -0.877 |
| P54116 | Stom | 0.556 | 0.825 | 0.677 | 0.000 | -0.243 | 0.446 | -0.539 | -0.245 |
| P99029 | Prdx5 | 0.996 | 1.000 | 0.234 | 0.000 | -0.266 | 0.051 | -0.201 | 0.121 |
| Q07417 | Acads | 0.730 | 1.000 | 0.089 | 0.000 | -0.125 | 0.435 | -0.537 | -0.263 |
| P12787 | Cox5a | 0.002 | 0.028 | 0.000 | 0.139 | -0.028 | 0.629 | 0.454 | 0.629 |
| Q9D0M3 | Cyc1 | 0.192 | 0.384 | 0.243 | 0.000 | -0.038 | 0.016 | 0.587 | 0.597 |
| Q80X85 | Mrps7 | 0.051 | 0.158 | 0.000 | 0.066 | -0.627 | 0.765 | 0.358 | 0.418 |
| P35486 | Pdha1 | 0.264 | 0.482 | 0.044 | 0.000 | -0.235 | 0.792 | 0.073 | -0.010 |
| Q1HFZ0 | Nsun2 | 0.102 | 0.247 | 0.289 | 0.000 | -0.086 | -0.029 | -1.110 | -0.849 |
| Q61171 | Prdx2 | 0.040 | 0.134 | 0.088 | 0.000 | -0.208 | -0.353 | -0.755 | -0.402 |
| Q8K3A0 | Hscb | 0.378 | 0.627 | 0.020 | 0.000 | -7.410 | -0.071 | 0.082 | -0.054 |
| Q9D6K5 | Synj2bp | 0.056 | 0.167 | 0.000 | 0.262 | -0.061 | 0.573 | 0.279 | 0.805 |
| Q61576 | Fkbp10 | 0.000 | 0.014 | 0.444 | -0.772 | 0.000 | 4.040 | 3.680 | 3.950 |
| Q9QZD8 | Slc25a10 | 0.003 | 0.030 | 0.000 | 0.214 | -0.415 | 1.190 | 1.260 | 1.510 |
| Q61733 | Mrps31 | 0.132 | 0.296 | 0.000 | 0.114 | -0.108 | 0.841 | -0.005 | 0.687 |
| Q9CQH3 | Ndufb5 | 0.003 | 0.032 | 0.000 | 0.202 | -0.151 | 0.682 | 0.684 | 0.812 |
| Q9Z1J3 | Nfs1 | 0.866 | 1.000 | 0.239 | 0.000 | -0.042 | 0.878 | -0.349 | -0.124 |
| P58059 | Mrps21 | 0.054 | 0.163 | 0.000 | 0.089 | -0.066 | 0.647 | 0.140 | 0.589 |
| P53395 | Dbt | 0.046 | 0.149 | 0.000 | 0.286 | -0.049 | 0.968 | 0.326 | 0.859 |
| Q9ESW4 | Agk | 0.183 | 0.372 | 0.000 | -0.572 | 0.077 | 0.527 | -0.141 | 0.703 |
| O89110 | Casp8 | 0.012 | 0.067 | 0.166 | -0.032 | 0.000 | -1.050 | -1.120 | -2.060 |
| Q9CZ83 | Mrpl55 | 0.003 | 0.031 | 0.173 | 0.000 | -0.267 | 0.903 | 0.758 | 0.928 |
| Q9DCT2 | Ndufs3 | 0.021 | 0.090 | -0.343 | 0.080 | 0.000 | 0.597 | 0.506 | 1.070 |
| P51660 | Hsd17b4 | 0.034 | 0.122 | 0.027 | 0.000 | -0.287 | 1.110 | 0.293 | 0.795 |
| Q8VCW8 | Acsf2 | 0.374 | 0.623 | 0.347 | -0.053 | 0.000 | 0.597 | -1.130 | -0.804 |
| Q8K1J6 | Trnt1 | 0.089 | 0.227 | -0.056 | 0.000 | 0.076 | -0.250 | -0.666 | -0.153 |
| Q9CPR5 | Mrpl15 | 0.032 | 0.119 | 0.144 | -1.190 | 0.000 | 1.030 | 0.900 | 1.190 |
| Q9DCW4 | Etfb | 0.322 | 0.558 | 0.252 | 0.000 | -0.128 | 0.867 | -0.052 | 0.291 |
| Q8VD26 | Tmem143 | 0.001 | 0.022 | -0.252 | 0.062 | 0.000 | 0.775 | 0.891 | 1.040 |
| Q9CQ54 | Ndufc2 | 0.001 | 0.022 | 0.044 | 0.000 | -0.180 | 0.889 | 0.645 | 0.923 |
| Q9CQ75 | Ndufa2 | 0.016 | 0.078 | -0.063 | 0.189 | 0.000 | 0.742 | 0.367 | 0.640 |
| P09671 | Sod2 | 0.023 | 0.096 | -0.061 | 0.348 | 0.000 | 0.715 | 0.476 | 0.682 |
| P08074 | Cbr2 | 0.001 | 0.019 | 0.000 | 0.042 | -1.580 | 5.280 | 4.340 | 4.610 |
| Q99LP6 | Grpel1 | 0.003 | 0.030 | 0.000 | 0.185 | -0.177 | 1.010 | 0.736 | 0.977 |
| Q9CR62 | Slc25a11 | 0.026 | 0.105 | 0.013 | 0.000 | -0.081 | 0.658 | 0.233 | 0.369 |
| P45952 | Acadm | 0.441 | 0.697 | 0.020 | 0.000 | -0.423 | 0.852 | -0.186 | -0.129 |
| Q80ZS3 | Mrps26 | 0.050 | 0.155 | -0.354 | 0.562 | 0.000 | 0.950 | 0.666 | 0.952 |
| Q9Z2Q5 | Mrpl40 | 0.003 | 0.031 | 0.154 | -0.056 | 0.000 | 0.713 | 0.525 | 0.546 |
| Q924T2 | Mrps2 | 0.014 | 0.073 | 0.000 | 0.348 | -0.330 | 0.736 | 1.060 | 0.937 |
| Q9CQA3 | Sdhb | 0.005 | 0.042 | 0.000 | 0.126 | -0.143 | 0.648 | 0.433 | 0.552 |
| Q9D3P8 | Plgrkt | 0.012 | 0.069 | -0.004 | 0.232 | 0.000 | 0.479 | 0.379 | 0.528 |

| | | | | | | | | | |
|---|---|---|---|---|---|---|---|---|---|
| P05202 | Got2 | 0.002 | 0.028 | 0.000 | 0.188 | -0.073 | 0.799 | 0.617 | 0.665 |
| O08756 | Hsd17b10 | 0.087 | 0.224 | -0.012 | 0.447 | 0.000 | 0.862 | 0.393 | 0.563 |
| Q64133 | Maoa | 0.000 | 0.011 | 0.151 | 0.000 | -0.761 | 3.640 | 3.580 | 3.770 |
| P22315 | Fech | 0.179 | 0.364 | 0.378 | -1.300 | 0.000 | 1.010 | 0.491 | 0.277 |
| Q9CPQ8 | Atp5l | 0.001 | 0.022 | -0.234 | 0.085 | 0.000 | 0.683 | 0.763 | 0.798 |
| Q3UMR5 | Mcu | 0.004 | 0.038 | 0.000 | 0.207 | -0.231 | 0.923 | 0.701 | 0.918 |
| Q9WUR2 | Eci2 | 0.392 | 0.641 | -0.002 | 0.400 | 0.000 | 0.637 | 0.088 | 0.274 |
| Q61578 | Fdxr | 0.099 | 0.243 | 0.000 | 0.172 | -0.061 | -0.043 | -0.801 | -0.540 |
| Q8JZQ2 | Afg3l2 | 0.015 | 0.076 | 0.000 | 0.089 | -0.109 | 0.828 | 0.358 | 0.590 |
| Q9D880 | Timm50 | 0.034 | 0.123 | 0.000 | 0.165 | -0.056 | 0.558 | 0.221 | 0.486 |
| Q9D855 | Uqcrb | 0.013 | 0.069 | 0.000 | 0.173 | -0.233 | 0.669 | 0.448 | 0.554 |
| Q5M8N4 | Sdr39u1 | 0.975 | 1.000 | -0.090 | 1.850 | 0.000 | -0.753 | 1.040 | 1.390 |
| P19536 | Cox5b | 0.287 | 0.512 | -0.201 | 0.104 | 0.000 | 0.204 | -0.054 | 0.216 |
| Q791V5 | Mtch2 | 0.007 | 0.048 | 0.000 | 0.565 | -0.017 | 1.160 | 1.140 | 1.190 |
| Q8BFP9 | Pdk1 | 0.386 | 0.634 | 0.043 | 0.000 | -0.101 | 0.101 | -0.625 | -0.165 |
| Q9EQ20 | Aldh6a1 | 0.368 | 0.615 | 0.300 | 0.000 | -0.150 | 1.460 | -0.061 | 0.226 |
| Q8K2M0 | Mrpl38 | 0.002 | 0.024 | 0.114 | 0.000 | -0.089 | 0.995 | 0.671 | 0.855 |
| P16125 | Ldhb | 0.000 | 0.013 | -0.212 | 0.132 | 0.000 | -1.330 | -1.570 | -1.550 |
| Q9DC50 | Crot | 0.119 | 0.276 | 0.000 | -0.124 | 0.034 | -0.050 | -0.367 | -0.340 |
| O35435 | Dhodh | 0.072 | 0.196 | 0.000 | 0.359 | -1.330 | 0.638 | 1.240 | 1.160 |
| O88441 | Mtx2 | 0.276 | 0.496 | 0.000 | 0.113 | -0.108 | 0.703 | -0.135 | 0.393 |
| Q8BJ03 | Cox15 | 0.015 | 0.075 | 0.000 | -0.638 | 0.346 | 1.240 | 1.100 | 1.810 |
| Q8R164 | Bphl | 0.031 | 0.114 | 0.197 | 0.000 | -0.133 | 0.714 | 0.305 | 0.778 |
| Q920E5 | Fdps | 0.104 | 0.252 | 1.020 | 0.000 | -0.336 | 1.880 | 0.878 | 1.120 |
| Q9DBF1 | Aldh7a1 | 0.000 | 0.009 | 0.000 | 0.318 | -0.191 | 2.700 | 2.460 | 2.520 |
| Q99MR8 | Mccc1 | 0.382 | 0.631 | 0.014 | 0.000 | -0.317 | 0.811 | -0.251 | 0.108 |
| Q9WVL0 | Gstz1 | 0.112 | 0.265 | 0.000 | 0.163 | -0.024 | 1.190 | 0.487 | 0.239 |
| Q922H2 | Pdk3 | 0.193 | 0.385 | 0.065 | 0.000 | -0.219 | 0.781 | -0.001 | 0.229 |
| Q9JMA2 | Qtrt1 | 0.001 | 0.019 | 0.000 | 0.059 | -0.158 | -0.700 | -0.758 | -0.891 |
| Q8VCX5 | Micu1 | 0.567 | 0.836 | -0.165 | 0.177 | 0.000 | 1.300 | -0.698 | 0.513 |
| Q99M87 | Dnaja3 | 0.029 | 0.112 | 0.003 | 0.000 | -0.329 | 0.713 | 0.216 | 0.625 |
| Q9CQ40 | Mrpl49 | 0.092 | 0.233 | 0.339 | 0.000 | -0.837 | 0.637 | 0.883 | 0.445 |
| Q8R3F5 | Mcat | 0.181 | 0.368 | 0.020 | 0.000 | -0.178 | 0.816 | -0.052 | 0.333 |
| Q9DCN2 | Cyb5r3 | 0.000 | 0.005 | 0.000 | 0.161 | -0.051 | 2.080 | 1.950 | 2.120 |
| Q9DB15 | Mrpl12 | 0.006 | 0.044 | 0.000 | 0.110 | -0.240 | 0.803 | 0.551 | 0.609 |
| Q99NB1 | Acss1 | 0.101 | 0.247 | 0.000 | 0.064 | -0.261 | -0.149 | -0.573 | -0.593 |
| Q9EQI8 | Mrpl46 | 0.017 | 0.081 | 0.194 | 0.000 | -0.083 | 1.000 | 0.471 | 0.691 |
| Q8C163 | Exog | 0.071 | 0.194 | -0.225 | 0.654 | 0.000 | 0.703 | 0.765 | 1.070 |
| Q5U458 | Dnajc11 | 0.066 | 0.185 | 0.771 | -0.042 | 0.000 | 1.780 | 1.580 | 0.657 |
| O08600 | Endog | 0.058 | 0.171 | -0.172 | 0.406 | 0.000 | -0.439 | -0.313 | -0.402 |
| O35459 | Ech1 | 0.045 | 0.146 | 0.000 | 0.120 | -0.262 | 0.291 | 0.279 | 0.264 |
| P47968 | Rpia | 0.160 | 0.337 | 0.718 | -0.577 | 0.000 | -0.741 | -2.220 | -0.389 |
| Q8VDC0 | Lars2 | 0.390 | 0.640 | 0.288 | 0.000 | -4.090 | 0.945 | -0.292 | -0.213 |
| Q3UHB1 | Nt5dc3 | 0.001 | 0.014 | 0.000 | 0.448 | -0.077 | 2.160 | 2.160 | 2.560 |
| Q9DCJ5 | Ndufa8 | 0.075 | 0.202 | 0.000 | 1.300 | -0.142 | 1.220 | 1.620 | 2.010 |

| | | | | | | | | | |
|---|---|---|---|---|---|---|---|---|---|
| Q9R257 | Hebp1 | 0.892 | 1.000 | 0.000 | 0.167 | -1.940 | 1.520 | -2.900 | -1.030 |
| Q921G7 | Etfdh | 0.009 | 0.055 | 0.011 | 0.000 | -0.217 | 0.785 | 0.394 | 0.573 |
| Q9JHS4 | Clpx | 0.557 | 0.825 | 0.335 | 0.000 | -0.328 | 1.540 | -0.635 | 0.363 |
| Q8BK72 | Mrps27 | 0.012 | 0.068 | 0.000 | 0.528 | -0.256 | 1.200 | 1.050 | 1.510 |
| Q8BW75 | Maob | 0.000 | 0.013 | 0.667 | 0.000 | -0.370 | 4.120 | 3.720 | 4.010 |
| Q922S4 | Pde2a | 0.073 | 0.199 | 0.280 | -0.058 | 0.000 | -0.909 | -2.760 | -0.790 |
| P48962 | Slc25a4 | 0.000 | 0.013 | 0.000 | 0.359 | -0.136 | 2.120 | 1.860 | 2.070 |
| Q9CQX2 | Cyb5b | 0.013 | 0.071 | -0.164 | 0.064 | 0.000 | 0.975 | 0.423 | 0.947 |
| Q9CPQ3 | Tomm22 | 0.010 | 0.061 | -0.029 | 0.000 | 0.186 | 1.040 | 0.523 | 0.946 |
| P70349 | Hint1 | 0.003 | 0.030 | -0.064 | 0.000 | 0.148 | -1.010 | -1.670 | -1.260 |
| P18155 | Mthfd2 | 0.034 | 0.122 | 0.000 | 0.357 | -0.941 | 0.795 | 1.290 | 1.380 |
| Q9DCZ4 | Apoo | 0.004 | 0.037 | 0.000 | 0.124 | -0.197 | 0.733 | 0.566 | 0.860 |
| Q9DCM0 | Ethe1 | 0.636 | 0.912 | 0.000 | 0.137 | -0.820 | 0.135 | -0.445 | 0.180 |
| Q99N93 | Mrpl16 | 0.172 | 0.355 | 0.000 | 0.020 | -0.274 | 0.737 | -0.068 | 0.326 |
| P62897 | Cycs | 0.205 | 0.401 | 0.000 | 0.081 | -0.533 | -0.300 | -0.469 | -0.708 |
| Q9D964 | Gatm | 0.821 | 1.000 | 0.770 | 0.000 | -0.380 | 0.678 | 0.053 | -0.044 |
| P31786 | Dbi | 0.001 | 0.019 | 0.000 | 0.100 | -0.167 | -1.290 | -1.720 | -1.290 |
| Q9DCU6 | Mrpl4 | 0.041 | 0.138 | 0.000 | 0.039 | -0.885 | 0.969 | 0.968 | 0.394 |
| Q9CR58 | Slc25a30 | 0.011 | 0.062 | 0.000 | 0.243 | -0.162 | 0.866 | 0.561 | 0.664 |
| Q791T5 | Mtch1 | 0.001 | 0.019 | 0.000 | 0.409 | -0.005 | 1.910 | 1.480 | 1.660 |
| Q9DB70 | Fundc1 | 0.256 | 0.470 | -0.517 | 1.650 | 0.000 | 2.470 | 0.712 | 1.270 |
| Q9CXZ1 | Ndufs4 | 0.012 | 0.067 | -0.105 | 0.323 | 0.000 | 0.881 | 0.660 | 1.120 |
| Q9JK81 | Myg1 | 0.012 | 0.068 | -0.941 | 0.000 | 0.003 | -1.670 | -1.890 | -1.610 |
| Q9CQN7 | Mrpl41 | 0.001 | 0.019 | -0.057 | 0.242 | 0.000 | 1.080 | 0.872 | 1.000 |
| O35129 | Phb2 | 0.003 | 0.030 | 0.000 | 0.062 | -0.164 | 0.922 | 0.591 | 0.718 |
| P97823 | Lypla1 | 0.168 | 0.350 | 0.000 | -0.250 | 0.196 | 0.058 | -1.140 | -0.961 |
| Q9CXW2 | Mrps22 | 0.536 | 0.803 | 0.837 | -1.070 | 0.000 | 1.960 | 1.660 | -1.400 |
| Q91VT4 | Cbr4 | 0.890 | 1.000 | 0.000 | 0.188 | -0.136 | 0.418 | -0.152 | -0.122 |
| Q9JKF7 | Mrpl39 | 0.494 | 0.759 | 0.000 | -3.790 | 0.010 | 1.100 | -0.144 | -1.440 |
| Q91VR2 | Atp5c1 | 0.035 | 0.125 | 0.127 | 0.000 | -0.028 | 1.150 | 0.535 | 0.475 |
| Q62425 | Ndufa4 | 0.219 | 0.419 | -0.214 | 0.210 | 0.000 | 0.195 | 0.073 | 0.541 |
| Q9D5T0 | Atad1 | 0.387 | 0.636 | 0.005 | -0.010 | 0.000 | 0.504 | -0.072 | 0.068 |
| Q9JKX6 | Nudt5 | 0.000 | 0.014 | 0.033 | -0.029 | 0.000 | -1.110 | -1.520 | -1.260 |
| Q8QZS1 | Hibch | 0.404 | 0.655 | 0.000 | 0.318 | -0.092 | 0.156 | 0.135 | 0.318 |
| Q99L04 | Dhrs1 | 0.470 | 0.729 | 0.040 | 0.000 | -0.227 | 0.315 | -0.898 | -0.478 |
| Q99KB8 | Hagh | 0.754 | 1.000 | 0.142 | 0.000 | -0.442 | -0.018 | -0.512 | -0.014 |
| Q9JLJ2 | Aldh9a1 | 0.081 | 0.213 | 0.030 | -0.088 | 0.000 | -0.239 | -1.410 | -0.779 |
| Q91VS7 | Mgst1 | 0.030 | 0.113 | 0.999 | 0.000 | -0.245 | 1.580 | 1.500 | 2.410 |
| Q91VM9 | Ppa2 | 0.070 | 0.193 | -0.028 | 0.474 | 0.000 | 0.613 | 0.457 | 0.874 |
| Q9D8P4 | Mrpl17 | 0.062 | 0.179 | -0.135 | 0.609 | 0.000 | 0.806 | 0.896 | 0.632 |
| P47791 | Gsr | 0.018 | 0.084 | 0.341 | 0.000 | -0.118 | -0.466 | -1.080 | -1.080 |
| Q9JLT4 | Txnrd2 | 0.839 | 1.000 | 0.000 | 0.351 | -0.041 | 0.125 | 0.029 | 0.247 |
| Q9CQ69 | Uqcrq | 0.004 | 0.038 | 0.000 | 0.165 | -0.022 | 0.638 | 0.470 | 0.464 |
| Q9CZU6 | Cs | 0.005 | 0.041 | 0.000 | 0.137 | -0.034 | 0.868 | 0.509 | 0.669 |
| O88967 | Yme1l1 | 0.017 | 0.081 | 0.000 | -0.237 | 0.021 | 0.822 | 0.328 | 0.838 |

| | | | | | | | | | |
|---|---|---|---|---|---|---|---|---|---|
| Q811U4 | Mfn1 | 0.420 | 0.674 | 0.000 | 1.260 | -0.739 | 2.090 | 1.140 | -0.275 |
| Q99J47 | Dhrs7b | 0.010 | 0.059 | 0.000 | 0.380 | -0.485 | 1.100 | 1.150 | 1.120 |
| Q9D6J5 | Ndufb8 | 0.039 | 0.132 | -0.884 | 0.149 | 0.000 | 0.958 | 0.797 | 0.597 |
| Q8BJ64 | Chdh | 0.375 | 0.623 | 0.970 | -0.135 | 0.000 | 0.659 | -1.190 | -0.567 |
| Q9CQN6 | Tmem14c | 0.066 | 0.186 | 0.146 | 0.000 | -0.210 | 0.760 | 0.282 | 0.302 |
| Q9CQL5 | Mrpl18 | 0.012 | 0.067 | -0.027 | 0.185 | 0.000 | 0.925 | 0.512 | 1.060 |
| Q8R2Q4 | Gfm2 | 0.142 | 0.309 | 0.000 | 0.144 | -0.128 | 0.042 | -1.100 | -0.903 |
| Q8JZU2 | Slc25a1 | 0.000 | 0.012 | 0.000 | 0.212 | -0.083 | 1.600 | 1.430 | 1.730 |
| Q8BU88 | Mrpl22 | 0.017 | 0.080 | 0.003 | -0.068 | 0.000 | 0.895 | 0.346 | 0.595 |
| P53702 | Hccs | 0.001 | 0.022 | -0.130 | 0.313 | 0.000 | 1.420 | 1.460 | 1.860 |
| P36552 | Cpox | 0.116 | 0.271 | 0.330 | 0.000 | -0.246 | -0.140 | -0.571 | -0.479 |
| Q9D0G0 | Mrps30 | 0.012 | 0.068 | 0.000 | 0.044 | -0.091 | 1.010 | 0.479 | 0.609 |
| Q9QYA2 | Tomm40 | 0.010 | 0.059 | 0.000 | 0.202 | -0.235 | 1.300 | 0.723 | 0.891 |
| Q8BYM8 | Cars2 | 0.412 | 0.665 | 0.076 | 0.000 | -0.350 | 0.743 | -0.258 | 0.119 |
| Q8BTX9 | Hsdl1 | 0.769 | 1.000 | 0.000 | 0.163 | -0.252 | 0.135 | -0.129 | 0.040 |
| Q8JZN7 | Rhot2 | 0.010 | 0.061 | 0.084 | 0.000 | -0.204 | 0.554 | 0.327 | 0.480 |
| Q9CXJ4 | Abcb8 | 0.014 | 0.074 | 0.000 | 0.642 | -0.065 | 1.350 | 1.020 | 1.280 |
| Q91WS0 | Cisd1 | 0.396 | 0.646 | -0.215 | 1.180 | 0.000 | 0.469 | 0.808 | 0.987 |
| O09174 | Amacr | 0.030 | 0.113 | 0.000 | 0.612 | -0.235 | 1.240 | 0.949 | 0.900 |
| Q9D7N3 | Mrps9 | 0.122 | 0.280 | 0.267 | -0.306 | 0.000 | 1.060 | 0.047 | 1.140 |
| Q8BWM0 | Ptges2 | 0.946 | 1.000 | -1.850 | 2.870 | 0.000 | 1.240 | 0.282 | -0.189 |
| Q99PU8 | Dhx30 | 0.416 | 0.670 | 0.000 | 0.380 | -0.875 | 1.330 | -0.133 | -0.053 |
| Q3URS9 | Ccdc51 | 0.007 | 0.048 | 0.194 | 0.000 | -0.221 | 0.635 | 0.568 | 0.670 |
| Q9DC70 | Ndufs7 | 0.005 | 0.040 | 0.000 | 0.110 | -0.184 | 0.684 | 0.466 | 0.575 |
| Q9EP89 | Lactb | 0.009 | 0.055 | 0.000 | 0.227 | -0.004 | 0.611 | 0.438 | 0.673 |
| Q9D3D9 | Atp5d | 0.126 | 0.287 | -0.224 | 0.095 | 0.000 | 0.573 | 0.056 | 0.267 |
| O88986 | Gcat | 0.640 | 0.917 | -0.398 | 0.000 | 0.308 | 0.250 | -0.442 | -0.349 |
| Q9JHI5 | Ivd | 0.333 | 0.571 | 0.000 | 0.127 | -0.011 | 0.544 | -0.052 | 0.212 |
| Q9CPP6 | Ndufa5 | 0.245 | 0.456 | -0.536 | 0.555 | 0.000 | 0.694 | 0.147 | 0.649 |
| O55125 | Nipsnap1 | 0.039 | 0.134 | -0.109 | 0.000 | 0.160 | 0.335 | 0.199 | 0.384 |
| Q8BX10 | Pgam5 | 0.212 | 0.409 | -0.303 | 0.066 | 0.000 | 0.400 | -0.081 | 0.256 |
| P48410 | Abcd1 | 0.804 | 1.000 | 0.401 | -0.082 | 0.000 | 0.518 | -0.244 | 0.258 |
| Q9WTP6 | Ak2 | 0.009 | 0.057 | 0.187 | 0.000 | -0.011 | -0.394 | -0.833 | -0.616 |
| Q8BG51 | Rhot1 | 0.015 | 0.076 | 0.000 | 0.146 | -0.220 | 0.857 | 0.556 | 0.462 |
| Q60759 | Gcdh | 0.450 | 0.708 | 0.000 | 0.355 | -0.371 | 0.773 | -0.053 | 0.092 |
| P20108 | Prdx3 | 0.006 | 0.046 | 0.214 | -0.058 | 0.000 | 0.778 | 0.509 | 0.675 |
| Q3U2A8 | Vars2 | 0.089 | 0.227 | 0.000 | 0.110 | -0.376 | 0.144 | 0.368 | 0.307 |
| Q80YD1 | Supv3l1 | 0.046 | 0.149 | 0.000 | 0.195 | -0.039 | 0.533 | 0.318 | 0.254 |
| Q99KK9 | Hars2 | 0.327 | 0.564 | 0.000 | 0.815 | -0.658 | 0.593 | 0.065 | 1.500 |
| Q91V12 | Acot7 | 0.020 | 0.089 | -0.038 | 0.193 | 0.000 | 0.736 | 0.449 | 0.396 |
| Q8BHF7 | Pgs1 | 0.017 | 0.080 | 0.000 | 1.120 | -0.175 | 2.030 | 1.790 | 2.010 |
| Q8K3J1 | Ndufs8 | 0.002 | 0.025 | 0.000 | 0.081 | -0.026 | 0.747 | 0.504 | 0.553 |
| Q64433 | Hspe1 | 0.001 | 0.018 | 0.000 | 0.100 | -0.080 | 1.090 | 0.779 | 0.991 |
| Q8K0D5 | Gfm1 | 0.253 | 0.467 | 0.000 | 0.253 | -0.248 | 1.020 | 0.078 | 0.217 |
| Q99N94 | Mrpl9 | 0.093 | 0.234 | 0.142 | 0.000 | -0.854 | 1.030 | 0.602 | 0.221 |

| | | | | | | | | | |
|---|---|---|---|---|---|---|---|---|---|
| Q8C3X2 | Ccdc90b | 0.005 | 0.040 | -0.005 | 0.303 | 0.000 | 1.660 | 1.020 | 1.250 |
| Q99N87 | Mrps5 | 0.000 | 0.012 | 0.039 | 0.000 | -0.040 | 1.140 | 0.934 | 0.888 |
| P51881 | Slc25a5 | 0.203 | 0.400 | 0.000 | 1.090 | -0.121 | 0.477 | 1.260 | 1.440 |
| Q8R1S0 | Coq6 | 0.178 | 0.364 | 0.000 | 0.621 | -0.148 | 0.936 | 0.515 | 0.414 |
| Q9DBL7 | Coasy | 0.134 | 0.298 | 7.440 | -0.978 | 0.000 | 7.090 | 7.330 | 7.010 |
| Q5HZI9 | Slc25a51 | 0.018 | 0.083 | -0.031 | 0.632 | 0.000 | 1.200 | 0.956 | 1.100 |
| Q91VC9 | Ghitm | 0.003 | 0.030 | 0.000 | 0.088 | -0.147 | 1.130 | 0.700 | 0.953 |
| Q8CAK1 | Iba57 | 0.322 | 0.558 | 0.036 | 0.000 | -0.449 | 0.439 | -0.205 | 0.176 |
| Q9D6K8 | Fundc2 | 0.013 | 0.071 | 0.000 | 0.474 | -0.255 | 1.690 | 1.410 | 0.958 |
| O08807 | Prdx4 | 0.256 | 0.470 | 0.038 | 0.000 | -0.002 | 0.439 | -0.070 | 0.262 |
| Q9QZ23 | Nfu1 | 0.022 | 0.094 | 0.000 | 0.235 | -0.727 | 1.080 | 0.755 | 1.350 |
| P51175 | Ppox | 0.909 | 1.000 | 0.000 | 0.082 | -0.711 | 0.651 | -0.730 | -0.373 |
| Q9JHR7 | Ide | 0.004 | 0.036 | 0.000 | 0.341 | -0.020 | -0.764 | -0.819 | -0.576 |
| Q9D1P0 | Mrpl13 | 0.522 | 0.787 | 0.000 | 0.348 | -0.702 | 0.959 | -0.679 | 0.596 |
| Q8K1M6 | Dnm1l | 0.892 | 1.000 | 0.042 | 0.000 | -0.313 | 0.147 | -0.185 | -0.310 |
| Q8K4X7 | Agpat4 | 0.005 | 0.042 | 0.000 | 0.300 | -0.338 | 1.030 | 0.953 | 1.130 |
| Q9CZL5 | Pcbd2 | 0.058 | 0.171 | 0.028 | 0.000 | -0.711 | 0.373 | 0.407 | 0.453 |
| P48771 | Cox7a2 | 0.957 | 1.000 | -0.456 | 0.000 | 0.309 | 0.089 | -0.409 | 0.125 |
| P70404 | Idh3g | 0.775 | 1.000 | 0.000 | 0.070 | -0.248 | 0.756 | -0.504 | -0.078 |
| Q9D6Y7 | Msra | 0.068 | 0.189 | 0.549 | -0.217 | 0.000 | -0.252 | -0.768 | -0.814 |
| P40630 | Tfam | 0.889 | 1.000 | 0.483 | 0.000 | -0.272 | 0.049 | 0.003 | 0.264 |
| P97386 | Lig3 | 0.578 | 0.849 | 0.068 | 0.000 | -0.173 | 0.026 | -0.596 | 0.058 |
| Q4VAE3 | Tmem65 | 0.014 | 0.072 | 0.000 | 0.487 | -0.336 | 1.190 | 1.030 | 1.570 |
| P56135 | Atp5j2 | 0.005 | 0.040 | 0.000 | 0.393 | -0.087 | 0.924 | 1.130 | 0.968 |
| Q9DCM2 | Gstk1 | 0.701 | 0.977 | 0.010 | 0.000 | -0.212 | 0.377 | -0.571 | -0.378 |
| Q9D0C4 | Trmt5 | 0.109 | 0.260 | -0.090 | 0.320 | 0.000 | -0.850 | -4.670 | -1.540 |
| Q924L1 | Letmd1 | 0.672 | 0.950 | -1.210 | 0.847 | 0.000 | -1.630 | 0.966 | -1.040 |
| Q9D8S4 | Rexo2 | 0.107 | 0.257 | -0.191 | 0.000 | 0.052 | 0.356 | 0.007 | 0.431 |
| Q7TSQ8 | Pdpr | 0.166 | 0.347 | 0.000 | 0.234 | -0.035 | 1.070 | 0.066 | 0.599 |
| Q9D7B6 | Acad8 | 0.170 | 0.353 | 0.000 | -1.970 | 0.001 | 1.380 | 0.114 | 0.357 |
| Q8K370 | Acad10 | 0.053 | 0.162 | 0.324 | 0.000 | -0.339 | 0.796 | 0.651 | 0.382 |
| P70677 | Casp3 | 0.003 | 0.030 | -0.016 | 0.138 | 0.000 | -1.190 | -2.040 | -1.570 |
| P61922 | Abat | 0.014 | 0.073 | 0.000 | 1.570 | -1.050 | 4.160 | 3.100 | 3.540 |
| Q9CQZ6 | Ndufb3 | 0.002 | 0.025 | -0.037 | 0.054 | 0.000 | 0.800 | 0.535 | 0.580 |
| Q9D6M3 | Slc25a22 | 0.001 | 0.016 | 0.000 | 0.088 | -0.280 | 1.510 | 1.170 | 1.410 |
| Q9D2R6 | Coa3 | 0.017 | 0.080 | 0.000 | 0.304 | -0.248 | 1.120 | 0.611 | 0.904 |
| P47934 | Crat | 0.395 | 0.644 | 0.000 | -3.600 | 0.145 | 2.180 | -1.640 | 0.743 |
| Q91WK1 | Spryd4 | 0.045 | 0.146 | 0.000 | 0.231 | -0.354 | 0.951 | 0.398 | 0.566 |
| Q9CQE1 | Nipsnap3b | 0.179 | 0.365 | 0.000 | 0.874 | -0.677 | 0.769 | 0.963 | 0.692 |
| Q9WVD5 | Slc25a15 | 0.005 | 0.042 | -0.508 | 0.364 | 0.000 | 1.270 | 1.750 | 1.660 |
| Q91Z53 | Grhpr | 0.060 | 0.176 | 0.000 | -0.087 | 0.077 | -0.125 | -0.652 | -0.656 |
| Q91VA6 | Poldip2 | 0.121 | 0.278 | 0.000 | 0.211 | -1.280 | 1.200 | 0.544 | 0.335 |
| Q9Z1P6 | Ndufa7 | 0.004 | 0.037 | -0.044 | 0.000 | 0.010 | 1.000 | 0.554 | 0.732 |
| P58281 | Opa1 | 0.111 | 0.263 | -0.010 | 0.051 | 0.000 | -0.089 | -0.563 | -0.194 |
| Q8VDT9 | Mrpl50 | 0.179 | 0.365 | 0.195 | -2.670 | 0.000 | 0.890 | 0.464 | 0.716 |

| | | | | | | | | | |
|---|---|---|---|---|---|---|---|---|---|
| Q8K215 | Lyrm4 | 0.019 | 0.086 | 0.000 | 0.224 | -0.231 | 0.718 | 0.439 | 0.602 |
| Q9JLZ3 | Auh | 0.243 | 0.453 | 0.028 | 0.000 | -0.092 | 0.540 | -0.170 | 0.570 |
| Q8BGA9 | Oxa1l | 0.039 | 0.133 | 0.000 | -1.550 | 0.191 | 1.400 | 1.020 | 1.330 |
| Q8BUY5 | Timmdc1 | 0.012 | 0.067 | 0.000 | 0.616 | -0.099 | 1.080 | 1.290 | 1.470 |
| Q9ERB0 | Snap29 | 0.440 | 0.697 | 0.870 | -3.570 | 0.000 | 0.006 | 0.284 | 0.520 |
| O54918 | Bcl2l11 | 0.580 | 0.851 | 0.000 | 0.199 | -0.024 | 0.457 | -0.129 | 0.176 |
| Q9D8Y1 | Tmem126a | 0.000 | 0.011 | 0.000 | 0.009 | -0.058 | 1.380 | 1.100 | 1.350 |
| Q3TL44 | Nlrx1 | 0.072 | 0.196 | 0.000 | 0.222 | -0.158 | 0.479 | 0.242 | 0.303 |
| Q9CZN8 | Qrsl1 | 0.705 | 0.982 | 0.000 | 0.121 | -0.527 | 0.510 | -0.341 | -0.174 |
| Q91VN4 | Chchd6 | 0.012 | 0.068 | 0.000 | 0.253 | -1.110 | 1.520 | 1.500 | 1.590 |
| Q9WU56 | Pus1 | 0.903 | 1.000 | 2.900 | -0.702 | 0.000 | 2.620 | 0.251 | -1.290 |
| Q71RI9 | Kyat3 | 0.016 | 0.079 | 0.000 | 0.041 | -0.711 | 0.821 | 0.680 | 0.805 |
| O88587 | Comt | 0.014 | 0.072 | -0.457 | 0.392 | 0.000 | 1.030 | 0.947 | 1.110 |
| Q9D338 | Mrpl19 | 0.721 | 1.000 | 0.000 | -0.168 | 0.031 | 1.100 | -0.582 | -0.080 |
| Q8BIP0 | Dars2 | 0.009 | 0.056 | 0.571 | -0.292 | 0.000 | 1.550 | 1.440 | 2.180 |
| Q99LB2 | Dhrs4 | 0.046 | 0.147 | 0.000 | 0.173 | -0.424 | 1.020 | 0.369 | 0.601 |
| Q8C5H8 | Nadk2 | 0.914 | 1.000 | 0.228 | 0.000 | -0.004 | 0.494 | -0.258 | -0.095 |
| Q9CQE3 | Mrps17 | 0.067 | 0.187 | 0.000 | 1.020 | -0.199 | 0.968 | 1.420 | 1.540 |
| Q9JKL4 | Ndufaf3 | 0.124 | 0.284 | 0.000 | -1.040 | 0.623 | 0.737 | 0.694 | 1.040 |
| Q8BHC4 | Dcakd | 0.017 | 0.081 | 0.466 | 0.000 | -1.560 | 2.210 | 1.810 | 2.370 |
| Q9D1I6 | Mrpl14 | 0.431 | 0.687 | 0.279 | 0.000 | -0.206 | 0.925 | -0.016 | 0.044 |
| P41216 | Acsl1 | 0.926 | 1.000 | 0.000 | 0.371 | -0.869 | 0.044 | 0.090 | -0.768 |
| P52503 | Ndufs6 | 0.095 | 0.237 | -0.325 | 0.000 | 0.739 | 0.967 | 0.626 | 1.040 |
| Q3U186 | Rars2 | 0.906 | 1.000 | 0.398 | -0.283 | 0.000 | 1.060 | -0.833 | -0.334 |
| Q3U5Q7 | Cmpk2 | 0.171 | 0.354 | 0.545 | 0.000 | -0.063 | 0.143 | -0.649 | -0.640 |
| Q8BK08 | Tmem11 | 0.001 | 0.019 | -0.043 | 0.285 | 0.000 | 1.080 | 0.954 | 1.080 |
| Q60649 | Clpb | 0.990 | 1.000 | 0.421 | 0.000 | -0.108 | 0.340 | -0.304 | 0.289 |
| Q3TC33 | Ccdc127 | 0.031 | 0.114 | -0.180 | 0.197 | 0.000 | 0.651 | 0.308 | 0.506 |
| Q9CY73 | Mrpl44 | 0.009 | 0.057 | 0.000 | 0.358 | -0.175 | 0.827 | 0.770 | 0.815 |
| Q80U63 | Mfn2 | 1.000 | 1.000 | 0.000 | 0.057 | -0.064 | -0.674 | 0.298 | 0.369 |
| Q9DC71 | Mrps15 | 0.711 | 0.987 | -0.194 | 0.000 | 0.044 | 0.841 | -0.657 | 0.193 |
| Q9CR59 | Gadd45gip1 | 0.130 | 0.293 | 0.000 | 0.151 | -0.194 | 0.895 | 0.031 | 0.578 |
| Q8BVU5 | Nudt9 | 0.155 | 0.330 | 0.000 | -0.351 | 0.224 | 0.896 | 2.140 | 0.088 |
| Q2TPA8 | Hsdl2 | 0.861 | 1.000 | 0.190 | -0.464 | 0.000 | 0.705 | -0.463 | -0.284 |
| Q9D8S9 | Bola1 | 0.460 | 0.719 | 0.000 | -2.370 | 0.187 | 1.050 | -0.214 | -0.646 |
| Q9CR68 | Uqcrfs1 | 0.007 | 0.048 | 0.000 | 0.074 | -0.106 | 0.656 | 0.355 | 0.524 |
| Q66GT5 | Ptpmt1 | 0.058 | 0.172 | 0.000 | 0.711 | -0.229 | 1.030 | 0.782 | 0.979 |
| Q9CQZ5 | Ndufa6 | 0.233 | 0.438 | 0.000 | 0.678 | -0.132 | 0.410 | 0.479 | 0.879 |
| Q99N89 | Mrpl43 | 0.003 | 0.030 | 0.000 | 0.185 | -0.319 | 1.180 | 0.974 | 1.330 |
| P56382 | Atp5e | 0.000 | 0.013 | 0.000 | 0.115 | -0.081 | 0.914 | 0.754 | 0.858 |
| Q9WV84 | Nme4 | 0.027 | 0.105 | 1.300 | -1.640 | 0.000 | 3.340 | 2.630 | 2.720 |
| Q9D1B9 | Mrpl28 | 0.304 | 0.534 | 0.000 | 0.671 | -1.410 | 0.836 | 0.095 | 0.635 |
| Q8BIG7 | Comtd1 | 0.937 | 1.000 | 0.000 | 0.166 | -0.276 | 0.673 | -0.609 | -0.276 |
| Q5IRJ6 | Slc30a9 | 0.003 | 0.030 | 0.000 | 0.577 | -0.728 | 2.360 | 2.540 | 2.810 |
| Q924D0 | Rtn4ip1 | 0.540 | 0.807 | 0.000 | 0.155 | -0.927 | 0.162 | -2.980 | -0.090 |

| | | | | | | | | | |
|---|---|---|---|---|---|---|---|---|---|
| Q9D023 | Mpc2 | 0.034 | 0.121 | 0.000 | 0.091 | -0.908 | 0.954 | 0.558 | 0.976 |
| Q8K2Y7 | Mrpl47 | 0.048 | 0.152 | 0.000 | 0.273 | -0.157 | 1.070 | 0.895 | 0.325 |
| Q9CPQ1 | Cox6c | 0.008 | 0.052 | 0.000 | 0.148 | -0.102 | 0.358 | 0.542 | 0.504 |
| Q99N85 | Mrps18a | 0.010 | 0.060 | 0.176 | -0.091 | 0.000 | 0.951 | 0.506 | 0.708 |
| P63030 | Mpc1 | 0.002 | 0.028 | 0.158 | -0.024 | 0.000 | 1.540 | 0.979 | 1.210 |
| Q921N7 | Tmem70 | 0.029 | 0.111 | -0.295 | 3.890 | 0.000 | 5.300 | 5.780 | 6.340 |
| Q9CYK1 | Wars2 | 0.461 | 0.720 | -0.361 | 0.518 | 0.000 | 0.617 | -0.291 | 0.913 |
| Q61586 | Gpam | 0.000 | 0.012 | 0.000 | -0.259 | 0.263 | 2.000 | 1.940 | 2.100 |
| Q8BYL4 | Yars2 | 0.539 | 0.807 | 0.319 | -1.240 | 0.000 | 0.125 | -1.890 | -0.671 |
| Q9ERI6 | Rdh14 | 0.189 | 0.379 | 0.000 | 0.132 | -0.126 | -0.072 | -0.191 | -0.680 |
| Q9D6U8 | Fam162a | 0.000 | 0.013 | -0.044 | 0.195 | 0.000 | 1.030 | 0.901 | 0.963 |
| Q9D125 | Mrps25 | 0.125 | 0.285 | 5.570 | -3.410 | 0.000 | 6.160 | 5.650 | 5.610 |
| Q9DC29 | Abcb6 | 0.117 | 0.272 | 0.022 | -0.201 | 0.000 | -0.174 | -0.821 | -0.399 |
| Q9CZ57 | Nsun4 | 0.430 | 0.686 | 0.129 | -1.190 | 0.000 | 0.559 | -0.895 | 1.310 |
| Q9DAT5 | Trmu | 0.965 | 1.000 | 0.330 | -0.121 | 0.000 | 0.746 | -0.391 | -0.095 |
| Q9D1R1 | Tmem126b | 0.441 | 0.698 | 0.146 | -0.371 | 0.000 | -0.573 | 0.779 | 0.798 |
| Q3TQB2 | Foxred1 | 0.796 | 1.000 | -0.086 | 0.930 | 0.000 | 0.212 | 0.095 | 0.873 |
| Q9CQ91 | Ndufa3 | 0.595 | 0.868 | -0.286 | 0.062 | 0.000 | 0.320 | -0.326 | 0.166 |
| Q99N95 | Mrpl3 | 0.067 | 0.187 | 0.000 | 0.558 | -0.395 | 1.000 | 0.576 | 0.857 |
| Q9DB10 | Smdt1 | 0.087 | 0.224 | -0.402 | 0.296 | 0.000 | 0.541 | 0.458 | 0.326 |

**Supp. Table 2.** Mitochondrial proteome was detected in $CD4^+$ T cells isolated from old mice after mito-transfer and in non-manipulated $CD4^+$ T cells from old mice. $CD4^+$ T cells from young and old mice, and from old mice after mito-transfer were cultured for 4 h before processing for mass spectrometry analysis. Data expressed as median protein Log2 fold change of $CD4^+$ T cells from old mice, from 3 individual old mice (paired experiment)

| Uniprot.ID | Gene.ID | P. Val | FDR | O1 | O2 | O3 | OM1 | OM2 | OM3 |
|---|---|---|---|---|---|---|---|---|---|
| Q8JZN5 | Acad9 | 0.215 | 0.412 | 0.000 | 0.150 | -0.210 | 0.570 | 0.120 | 0.080 |
| Q9Z0X1 | Aifm1 | 0.917 | 1.000 | 0.000 | 0.040 | -0.030 | 0.160 | -0.130 | 0.010 |
| Q3TQB2 | Foxred1 | 0.796 | 1.000 | -0.090 | 0.930 | 0.000 | 0.210 | 0.100 | 0.870 |
| Q99LC3 | Ndufa10 | 0.006 | 0.047 | 0.000 | 0.010 | -0.250 | 0.720 | 0.460 | 0.860 |
| Q7TMF3 | Ndufa12 | 0.011 | 0.064 | -0.160 | 0.000 | 0.140 | 0.790 | 0.550 | 1.110 |
| Q9ERS2 | Ndufa13 | 0.056 | 0.168 | 0.040 | -0.180 | 0.000 | 0.650 | 0.140 | 0.880 |
| Q9CQ75 | Ndufa2 | 0.016 | 0.078 | -0.060 | 0.190 | 0.000 | 0.740 | 0.370 | 0.640 |

| | | | | | | | | | |
|---|---|---|---|---|---|---|---|---|---|
| Q9CQ91 | Ndufa3 | 0.595 | 0.868 | -0.290 | 0.060 | 0.000 | 0.320 | -0.330 | 0.170 |
| Q9CPP6 | Ndufa5 | 0.245 | 0.456 | -0.540 | 0.560 | 0.000 | 0.690 | 0.150 | 0.650 |
| Q9CQZ5 | Ndufa6 | 0.233 | 0.438 | 0.000 | 0.680 | -0.130 | 0.410 | 0.480 | 0.880 |
| Q9Z1P6 | Ndufa7 | 0.004 | 0.037 | -0.040 | 0.000 | 0.010 | 1.000 | 0.550 | 0.730 |
| Q9DCJ5 | Ndufa8 | 0.075 | 0.202 | 0.000 | 1.300 | -0.140 | 1.220 | 1.620 | 2.010 |
| Q9DC69 | Ndufa9 | 0.004 | 0.037 | -0.070 | 0.100 | 0.000 | 0.740 | 0.440 | 0.640 |
| Q9JKL4 | Ndufaf3 | 0.124 | 0.284 | 0.000 | -1.040 | 0.620 | 0.740 | 0.690 | 1.040 |
| Q9DCS9 | Ndufb10 | 0.006 | 0.043 | 0.000 | 0.250 | -0.110 | 0.630 | 0.920 | 0.810 |
| O09111 | Ndufb11 | 0.001 | 0.018 | 0.020 | -0.200 | 0.000 | 0.580 | 0.570 | 0.650 |
| Q9CQZ6 | Ndufb3 | 0.002 | 0.025 | -0.040 | 0.050 | 0.000 | 0.800 | 0.540 | 0.580 |
| Q9CQC7 | Ndufb4 | 0.005 | 0.039 | 0.000 | 0.250 | -0.030 | 0.720 | 0.760 | 0.570 |
| Q9CQH3 | Ndufb5 | 0.003 | 0.032 | 0.000 | 0.200 | -0.150 | 0.680 | 0.680 | 0.810 |
| Q9CR61 | Ndufb7 | 0.020 | 0.088 | 0.000 | -0.040 | 0.110 | 0.720 | 0.440 | 1.080 |
| Q9D6J5 | Ndufb8 | 0.039 | 0.132 | -0.880 | 0.150 | 0.000 | 0.960 | 0.800 | 0.600 |
| Q9CQJ8 | Ndufb9 | 0.003 | 0.033 | -0.200 | 0.220 | 0.000 | 0.740 | 0.740 | 0.810 |
| Q9CQ54 | Ndufc2 | 0.001 | 0.022 | 0.040 | 0.000 | -0.180 | 0.890 | 0.650 | 0.920 |
| Q91VD9 | Ndufs1 | 0.002 | 0.025 | 0.000 | 0.200 | -0.070 | 0.850 | 0.650 | 0.790 |
| Q91WD5 | Ndufs2 | 0.006 | 0.045 | 0.000 | 0.410 | -0.120 | 0.950 | 0.910 | 1.040 |
| Q9DCT2 | Ndufs3 | 0.021 | 0.090 | -0.340 | 0.080 | 0.000 | 0.600 | 0.510 | 1.070 |
| Q9CXZ1 | Ndufs4 | 0.012 | 0.067 | -0.110 | 0.320 | 0.000 | 0.880 | 0.660 | 1.120 |
| P52503 | Ndufs6 | 0.095 | 0.237 | -0.330 | 0.000 | 0.740 | 0.970 | 0.630 | 1.040 |
| Q9DC70 | Ndufs7 | 0.005 | 0.040 | 0.000 | 0.110 | -0.180 | 0.680 | 0.470 | 0.570 |
| Q8K3J1 | Ndufs8 | 0.002 | 0.025 | 0.000 | 0.080 | -0.030 | 0.750 | 0.500 | 0.550 |
| Q91YT0 | Ndufv1 | 0.001 | 0.019 | 0.000 | 0.030 | -0.080 | 0.820 | 0.550 | 0.690 |
| Q9D6J6 | Ndufv2 | 0.049 | 0.154 | 0.000 | 0.340 | -0.180 | 0.720 | 0.360 | 0.860 |
| Q8BUY5 | Timmdc1 | 0.012 | 0.067 | 0.000 | 0.620 | -0.100 | 1.080 | 1.290 | 1.470 |
| Q9D8Y1 | Tmem126a | 0.000 | 0.011 | 0.000 | 0.010 | -0.060 | 1.380 | 1.100 | 1.350 |
| Q9D1R1 | Tmem126b | 0.441 | 0.698 | 0.150 | -0.370 | 0.000 | -0.570 | 0.780 | 0.800 |
| Q921N7 | Tmem70 | 0.029 | 0.111 | -0.290 | 3.890 | 0.000 | 5.300 | 5.780 | 6.340 |
| Q8K2B3 | Sdha | 0.017 | 0.081 | 0.000 | 0.000 | -0.200 | 0.720 | 0.300 | 0.450 |
| Q9CQA3 | Sdhb | 0.005 | 0.042 | 0.000 | 0.130 | -0.140 | 0.650 | 0.430 | 0.550 |
| Q9D0M3 | Cyc1 | 0.192 | 0.384 | 0.240 | 0.000 | -0.040 | 0.020 | 0.590 | 0.600 |
| Q9D855 | Uqcrb | 0.013 | 0.069 | 0.000 | 0.170 | -0.230 | 0.670 | 0.450 | 0.550 |
| Q9CZ13 | Uqcrc1 | 0.015 | 0.076 | 0.080 | 0.000 | -0.050 | 0.660 | 0.290 | 0.460 |
| Q9DB77 | Uqcrc2 | 0.140 | 0.306 | 0.000 | 0.130 | -0.100 | 0.560 | 0.050 | 0.310 |
| Q9CR68 | Uqcrfs1 | 0.007 | 0.048 | 0.000 | 0.070 | -0.110 | 0.660 | 0.350 | 0.520 |
| Q9CQ69 | Uqcrq | 0.004 | 0.038 | 0.000 | 0.170 | -0.020 | 0.640 | 0.470 | 0.460 |
| Q9D2R6 | Coa3 | 0.017 | 0.080 | 0.000 | 0.300 | -0.250 | 1.120 | 0.610 | 0.900 |
| Q8BJ03 | Cox15 | 0.015 | 0.075 | 0.000 | -0.640 | 0.350 | 1.240 | 1.100 | 1.810 |
| P19783 | Cox4i1 | 0.004 | 0.038 | 0.100 | -0.020 | 0.000 | 0.710 | 0.420 | 0.700 |
| P12787 | Cox5a | 0.002 | 0.028 | 0.000 | 0.140 | -0.030 | 0.630 | 0.450 | 0.630 |
| P19536 | Cox5b | 0.287 | 0.512 | -0.200 | 0.100 | 0.000 | 0.200 | -0.050 | 0.220 |
| P56391 | Cox6b1 | 0.015 | 0.076 | 0.030 | 0.000 | -0.080 | 0.410 | 0.170 | 0.410 |
| Q9CPQ1 | Cox6c | 0.008 | 0.052 | 0.000 | 0.150 | -0.100 | 0.360 | 0.540 | 0.500 |
| P48771 | Cox7a2 | 0.957 | 1.000 | -0.460 | 0.000 | 0.310 | 0.090 | -0.410 | 0.130 |

| | | | | | | | | | |
|---|---|---|---|---|---|---|---|---|---|
| Q62425 | Ndufa4 | 0.219 | 0.419 | -0.210 | 0.210 | 0.000 | 0.200 | 0.070 | 0.540 |
| P09925 | Surf1 | 0.000 | 0.013 | 0.070 | 0.000 | -0.070 | 0.680 | 0.590 | 0.720 |
| Q03265 | Atp5a1 | 0.002 | 0.025 | 0.000 | 0.050 | -0.110 | 0.800 | 0.540 | 0.810 |
| P56480 | Atp5b | 0.009 | 0.057 | 0.000 | 0.260 | -0.160 | 0.720 | 0.570 | 0.810 |
| Q91VR2 | Atp5c1 | 0.035 | 0.125 | 0.130 | 0.000 | -0.030 | 1.150 | 0.540 | 0.470 |
| Q9D3D9 | Atp5d | 0.126 | 0.287 | -0.220 | 0.090 | 0.000 | 0.570 | 0.060 | 0.270 |
| P56382 | Atp5e | 0.000 | 0.013 | 0.000 | 0.120 | -0.080 | 0.910 | 0.750 | 0.860 |
| Q9DCX2 | Atp5h | 0.025 | 0.101 | 0.000 | 0.470 | -0.340 | 1.070 | 1.120 | 0.720 |
| P97450 | Atp5j | 0.033 | 0.121 | 0.000 | 0.130 | -0.630 | 1.010 | 0.420 | 1.140 |
| P56135 | Atp5j2 | 0.005 | 0.040 | 0.000 | 0.390 | -0.090 | 0.920 | 1.130 | 0.970 |
| Q9CPQ8 | Atp5l | 0.001 | 0.022 | -0.230 | 0.080 | 0.000 | 0.680 | 0.760 | 0.800 |
| Q9DB20 | Atp5o | 0.001 | 0.019 | 0.000 | 0.130 | -0.070 | 0.900 | 0.670 | 0.900 |
| O35143 | Atpif1 | 0.013 | 0.069 | 0.260 | 0.000 | -0.320 | 1.590 | 1.150 | 0.810 |
| Q921N7 | Tmem70 | 0.029 | 0.111 | -0.290 | 3.890 | 0.000 | 5.300 | 5.780 | 6.340 |

**Supp. Table 3.** Proteins related to ETC were detected in CD4$^+$ T cells isolated from old mice after mito-transfer and in non-manipulated CD4$^+$ T cells from old mice. CD4$^+$ T cells from young and old mice, and from old mice after mito-transfer were cultured for 4 h before processing for mass spectrometry analysis. Data expressed as median protein Log2 fold change of CD4$^+$ T cells from old mice, from 3 individual old mice (paired experiment)

| Uniprot ID | Gene ID | P.Val | FDR | O1 | O2 | O3 | OM1 | OM2 | OM3 |
|---|---|---|---|---|---|---|---|---|---|
| Q9R1C7 | Prpf40a | 0.057 | 0.169 | 0.000 | 0.036 | -0.175 | -0.196 | -0.679 | -0.769 |
| Q9ES52 | Inpp5d | 0.069 | 0.191 | -0.054 | 0.183 | 0.000 | -0.248 | -0.967 | -0.380 |
| Q9ERK4 | Cse1l | 0.041 | 0.138 | 0.073 | -0.029 | 0.000 | -0.350 | -1.430 | -0.960 |
| Q9DBG6 | Rpn2 | 0.000 | 0.012 | 0.000 | 0.145 | -0.216 | 1.580 | 1.360 | 1.520 |
| Q9D358 | Acp1 | 0.010 | 0.060 | 0.000 | 0.034 | -0.377 | -0.763 | -1.300 | -1.250 |
| Q9CX99 | Grap | 0.077 | 0.207 | -0.922 | 0.152 | 0.000 | -1.990 | -2.090 | -0.687 |
| Q99L45 | Eif2s2 | 0.540 | 0.807 | 0.342 | -0.259 | 0.000 | 0.266 | -0.631 | -0.179 |
| Q99JR1 | Sfxn1 | 0.018 | 0.083 | 0.000 | 0.180 | -0.179 | 0.403 | 0.408 | 0.393 |
| Q99JF8 | Psip1 | 0.090 | 0.230 | -0.095 | 0.000 | 0.152 | -0.474 | -0.546 | -0.054 |
| Q921F2 | Tardbp | 0.080 | 0.211 | 0.004 | 0.000 | -0.021 | -0.142 | -0.728 | -0.349 |
| Q8QZY9 | Sf3b4 | 0.531 | 0.798 | -2.010 | 0.478 | 0.000 | -3.370 | -0.510 | -0.237 |
| Q8K4I3 | Arhgef6 | 0.010 | 0.058 | -0.259 | 0.000 | 0.183 | -0.812 | -1.510 | -1.100 |

| | | | | | | | | | |
|---|---|---|---|---|---|---|---|---|---|
| Q8K2Z4 | Ncapd2 | 0.437 | 0.694 | 0.059 | -0.356 | 0.000 | 0.217 | -0.607 | -0.761 |
| Q8K1I7 | Wipf1 | 0.146 | 0.315 | -0.001 | 0.000 | 0.402 | -0.311 | -0.419 | 0.062 |
| Q8CC88 | Vwa8 | 0.942 | 1.000 | 0.000 | 0.045 | -0.313 | 0.847 | -0.752 | -0.481 |
| Q8C3J5 | Dock2 | 0.006 | 0.047 | 0.009 | 0.000 | -0.011 | -0.628 | -1.100 | -0.665 |
| Q8C2K5 | Rasal3 | 0.018 | 0.083 | -0.082 | 0.141 | 0.000 | -0.512 | -1.320 | -0.924 |
| Q8BZN6 | Dock10 | 0.210 | 0.406 | -0.532 | 0.000 | 0.228 | -0.481 | -1.600 | -0.303 |
| Q8BK67 | Rcc2 | 0.253 | 0.466 | 0.487 | -0.092 | 0.000 | 0.079 | -0.473 | -0.172 |
| Q8BH59 | Slc25a12 | 0.037 | 0.127 | -0.084 | 0.153 | 0.000 | 0.368 | 0.204 | 0.480 |
| Q8BGW0 | Themis | 0.127 | 0.288 | -1.320 | 0.000 | 0.443 | -1.890 | -1.260 | -1.100 |
| Q80SU7 | Gvin1 | 0.117 | 0.273 | 0.231 | 0.000 | -0.019 | 0.045 | -0.881 | -0.733 |
| Q78ZA7 | Nap1l4 | 0.009 | 0.056 | -0.293 | 0.000 | 0.003 | -0.919 | -1.530 | -0.967 |
| Q6ZQ38 | Cand1 | 0.009 | 0.057 | -0.021 | 0.028 | 0.000 | -0.389 | -0.813 | -0.557 |
| Q6PDI5 | Ecm29 | 0.133 | 0.297 | 0.000 | -0.153 | 0.001 | 0.008 | -1.570 | -1.600 |
| Q6PB66 | Lrpprc | 0.035 | 0.125 | 0.000 | 0.121 | -0.292 | 0.941 | 0.428 | 0.430 |
| Q62351 | Tfrc | 0.848 | 1.000 | 1.330 | 0.000 | -0.491 | 1.130 | -0.397 | -0.350 |
| Q62077 | Plcg1 | 0.003 | 0.033 | -0.125 | 0.000 | 0.471 | -1.190 | -1.370 | -1.750 |
| Q61823 | Pdcd4 | 0.011 | 0.065 | -0.098 | 0.000 | 0.177 | -1.090 | -2.030 | -1.100 |
| Q61081 | Cdc37 | 0.002 | 0.027 | 0.019 | 0.000 | -0.009 | -0.837 | -1.370 | -1.050 |
| Q60932 | Vdac1 | 0.006 | 0.046 | 0.000 | 0.313 | -0.052 | 1.240 | 1.010 | 1.700 |
| Q60931 | Vdac3 | 0.007 | 0.050 | 0.000 | 0.252 | -0.058 | 1.080 | 0.902 | 1.570 |
| Q60787 | Lcp2 | 0.009 | 0.057 | -0.096 | 0.845 | 0.000 | -1.390 | -1.550 | -1.040 |
| Q60631 | Grb2 | 0.005 | 0.041 | -0.091 | 0.403 | 0.000 | -1.080 | -1.510 | -0.944 |
| Q3UUV5 | Skap1 | 0.002 | 0.023 | -0.314 | 0.000 | 0.236 | -1.930 | -2.300 | -1.630 |
| Q3UPF5 | Zc3hav1 | 0.070 | 0.193 | -0.342 | 0.000 | 0.012 | -0.376 | -0.579 | -0.367 |
| Q3UND0 | Skap2 | 0.206 | 0.403 | 0.518 | 0.000 | -0.624 | -0.258 | -0.599 | -1.130 |
| Q3TBT3 | Tmem17 3 | 0.649 | 0.925 | -0.395 | 0.164 | 0.000 | -0.291 | -0.323 | 0.074 |
| Q1HFZ0 | Nsun2 | 0.102 | 0.247 | 0.289 | 0.000 | -0.086 | -0.029 | -1.110 | -0.849 |
| Q03526 | Itk | 0.005 | 0.041 | 0.620 | -0.355 | 0.000 | -2.150 | -1.480 | -1.840 |
| Q01965 | Ly9 | 0.539 | 0.807 | 0.248 | -0.106 | 0.000 | -0.213 | -0.306 | 0.254 |
| P97370 | Atp1b3 | 0.009 | 0.056 | 0.000 | 0.096 | -0.048 | 0.414 | 0.419 | 0.703 |
| P70218 | Map4k1 | 0.001 | 0.015 | -0.001 | 0.000 | 0.020 | -0.901 | -1.290 | -1.110 |
| P70168 | Kpnb1 | 0.204 | 0.400 | -0.040 | 0.088 | 0.000 | -0.081 | -0.545 | -0.064 |
| P68254 | Ywhaq | 0.006 | 0.046 | -0.139 | 0.000 | 0.050 | -0.858 | -1.630 | -1.230 |
| P63101 | Ywhaz | 0.001 | 0.015 | 0.000 | -0.021 | 0.113 | -1.140 | -1.610 | -1.370 |
| P63037 | Dnaja1 | 0.340 | 0.580 | 0.072 | 0.000 | -0.107 | 0.131 | -0.492 | -0.294 |
| P62259 | Ywhae | 0.027 | 0.105 | 0.000 | 0.068 | -0.183 | -0.420 | -1.160 | -0.884 |
| P61982 | Ywhag | 0.016 | 0.078 | 0.076 | 0.000 | -0.065 | -0.399 | -0.977 | -0.678 |
| P61290 | Psme3 | 0.001 | 0.021 | 0.000 | 0.420 | -0.049 | -1.120 | -1.150 | -1.340 |
| P54775 | Psmc4 | 0.018 | 0.083 | -0.123 | 0.000 | 0.106 | -0.391 | -0.884 | -0.580 |
| P49718 | Mcm5 | 0.432 | 0.688 | 0.140 | 0.000 | -0.032 | 0.135 | -0.802 | -0.001 |
| P43404 | Zap70 | 0.038 | 0.131 | -0.465 | 0.041 | 0.000 | -0.669 | -0.946 | -0.589 |
| P42227 | Stat3 | 0.103 | 0.250 | 0.000 | -0.155 | 0.131 | -0.089 | -0.977 | -0.688 |

| | | | | | | | | | |
|---|---|---|---|---|---|---|---|---|---|
| P42225 | Stat1 | 0.005 | 0.039 | -0.079 | 0.271 | 0.000 | -0.603 | -0.992 | -0.866 |
| P39688 | Fyn | 0.582 | 0.853 | -0.633 | 0.103 | 0.000 | -0.287 | -0.529 | -0.169 |
| P39054 | Dnm2 | 0.326 | 0.562 | 0.075 | -0.001 | 0.000 | 0.448 | -0.098 | 0.271 |
| P36371 | Tap2 | 0.787 | 1.000 | -0.225 | 0.273 | 0.000 | -0.153 | -0.078 | 0.133 |
| P27870 | Vav1 | 0.049 | 0.154 | 0.325 | 0.000 | -0.114 | -0.272 | -1.240 | -1.350 |
| P26450 | Pik3r1 | 0.468 | 0.727 | 0.000 | 0.958 | -0.100 | -0.021 | 0.030 | 0.038 |
| P26039 | Tln1 | 0.041 | 0.137 | 0.059 | 0.000 | -0.282 | -0.358 | -1.020 | -1.000 |
| P25206 | Mcm3 | 0.066 | 0.185 | 0.280 | 0.000 | -0.196 | -0.205 | -1.110 | -0.943 |
| P24161 | Cd247 | 0.811 | 1.000 | -0.617 | 0.000 | 0.247 | -0.627 | 0.147 | 0.420 |
| P24063 | Itgal | 0.054 | 0.163 | -0.246 | 0.010 | 0.000 | -0.641 | -0.514 | -0.247 |
| P22682 | Cbl | 0.000 | 0.013 | -0.024 | 0.000 | 0.154 | -1.190 | -1.550 | -1.380 |
| P22646 | Cd3e | 0.501 | 0.766 | -0.349 | 0.000 | 0.371 | -0.475 | -0.277 | 0.156 |
| P19783 | Cox4i1 | 0.004 | 0.038 | 0.100 | -0.024 | 0.000 | 0.711 | 0.422 | 0.696 |
| P11942 | Cd3g | 0.167 | 0.349 | -0.367 | 0.000 | 0.062 | -0.670 | -0.495 | -0.158 |
| P11352 | Gpx1 | 0.131 | 0.294 | 0.457 | 0.000 | -0.235 | -0.028 | -0.804 | -0.877 |
| P08113 | Hsp90b1 | 0.004 | 0.035 | 0.176 | 0.000 | -0.151 | 1.090 | 0.696 | 0.949 |
| P04235 | Cd3d | 0.281 | 0.504 | -0.353 | 0.097 | 0.000 | -0.435 | -0.393 | -0.084 |
| P01851 | Tcb2 | 0.061 | 0.176 | -0.290 | 0.000 | 0.247 | -0.718 | -0.420 | -0.372 |
| O89100 | Grap2 | 0.049 | 0.154 | -0.257 | 0.110 | 0.000 | -0.559 | -2.360 | -1.760 |
| O54957 | Lat | 0.507 | 0.771 | -0.255 | 0.567 | 0.000 | 0.081 | 0.134 | 0.862 |
| O54734 | Ddost | 0.002 | 0.025 | 0.000 | 0.042 | -0.469 | 1.490 | 1.120 | 1.220 |

**Supp. Table 4.** Proteins related to TCR signalosome were detected in CD4$^+$ T cells isolated from old mice after mito-transfer and in non-manipulated CD4$^+$ T cells from old mice. CD4$^+$ T cells from young and old mice, and from old mice after mito-transfer were cultured for 4 h before processing for mass spectrometry analysis. Data expressed as median protein Log2 fold change of CD4$^+$ T cells from old mice, from 3 individual old mice (paired experiment)
